# Genome writing and Targeted Delivery of the *NKX6-3/ANK1* gene cluster and its Type 2 Diabetes GWAS Variants to Human iPSCs

**DOI:** 10.64898/2026.01.04.697539

**Authors:** Noor Chalhoub, Arushi Varshney, Weimin Zhang, Skyler Uhl, Jon M. Laurent, Colleen McLoughlin, Hannah Ashe, Xingrui Mou, Nathan Dale, Kiran Ramnarine, Daniel Paull, Jordan Goldberg, Matthew T. Maurano, Ran Brosh, David Fenyö, Filippo Cipriani, Stephen C.J. Parker, Jef D. Boeke

## Abstract

Genome-wide association studies (GWAS) identified over 600 loci containing single-nucleotide polymorphisms (SNPs) associated with type 2 diabetes (T2D), most of which reside in non-coding regions. Among the set of T2D SNPs, linking causal genome variants to disease risk experimentally has remained a challenge; however, advances in synthetic mammalian genome writing techniques now enable the delivery of multiple haplotypes to human induced pluripotent stem cells (hiPSCs) to create a series of isogenic cell lines that can be differentiated and phenotyped *in vitro*. Here, to begin efforts in dissecting a T2D GWAS locus, we engineered an *NKX6-3/ANK1* gene cluster knockout hiPSC line and introduced a landing pad facilitating the delivery of synthetic haplotype payloads. We built four haplotypes, including several that are not observed in nature, containing risk SNPs spanning the *NKX6-3/ANK1* gene cluster using a method called “variant Switching Auxotrophic markers for Integration” (vSwAP-In), and integrated them precisely into hiPSCs. *NKX6-3/ANK1* deletion blocked pancreatic progenitor and skeletal muscle differentiation, suggesting that *NKX6-3* and *ANK1* are required for early pancreatic and skeletal muscle development, and perhaps related to the existence of two nonoverlapping sets of SNPs in linkage disequilibrium that associate with the expression of the two adjacent genes. When *NKX6-3*/*ANK1* T2D “Risk” haplotypes were reintroduced, skeletal muscle and pancreatic progenitor differentiation capabilities were restored. *ANK1* expression was elevated in the *ANK1* Risk and All-Risk haplotypes compared to the *NKX6-3* Risk and Non-Risk haplotypes, establishing a functional experimental platform to examine risk SNP clusters in their native contexts. Overall, this work establishes a platform for the dissection of GWAS loci using synthetic haplotype genomics in hiPSCs.

**Significance Statement:** Genome-wide association studies have been used to identify disease-associated SNPs; however, most SNPs lie in non-coding regions, making functional experimentation difficult to perform. Using vSwAP-In, a yeast-based DNA variant-building method, and mSwAP-In, a mammalian genome engineering approach, we establish a platform for functional GWAS dissection in hiPSCs. This platform allows us to build DNA harboring virtually any combination of disease-risk SNPs, allowing for functional characterization of SNPs without the limitations of linkage disequilibrium. We demonstrate this approach using a Type 2 diabetes GWAS gene cluster, *NKX6-3/ANK1*.

## Introduction

Type 2 diabetes (T2D) has become widely accepted as a polygenic disease with multiple genetic factors influencing an individual’s risk (1). Genome-wide association studies (GWAS) have been used to study complex diseases, such as T2D, by identifying genetic variants associated with disease risk (2). Single-nucleotide polymorphisms (SNPs) and other genetic variants associated with Type 2 diabetes (T2D) risk have been identified through GWAS conducted across various populations (3) with approximately 600 risk loci, and over 1,200 independent signals reported (4). Statistical association is influenced by the allele frequency of variants within the population, and the correlation of variants across the population, known as linkage disequilibrium (LD) (5, 6). GWAS SNPs are often found in non-coding regions and are typically in high LD with surrounding SNPs, making it difficult to pinpoint causal SNPs in the genomic locus that may be contributing to disease risk (2, 5-7).

The *NKX6-3/ANK1* gene cluster, located on chromosome 8, has been reported as a T2D risk locus, with independent signals or sets of risk SNPs identified mainly in European and East Asian populations (1, 8-11). Most studies have shown how these SNPs interact in the context of T2D-related tissues through expression quantitative trait loci (eQTL) experiments computationally (9, 12-15), while other studies investigated the overexpression of a short, muscle-specific isoform of *ANK1* known as *sANK1* and the role of select SNPs outside their native contexts (16, 17). There are currently no functional studies examining T2D-risk SNPs at the *NKX6-3/ANK1* locus in their native context.

Here, we report a platform in which fully synthetic variant haplotypes can be engineered in yeast (*S. cerevisiae*) and delivered to hiPSCs to study GWAS loci. We developed a method called vSwAP-In (variant Switching of Auxotrophic markers Progressively for Integration) using similar principles we previously reported (18) to build genomic payloads containing risk SNP combinations that are not observed in nature, allowing us to study risk SNPs without the limitations of LD in natural populations. We show here that these haplotype payloads can be delivered as “Big DNA” molecules to hiPSCs using the mSwAP-In delivery method (19, 20). In mSwAP-In, Big DNA payloads of up to ∼200 kb are delivered to mouse embryonic stem cells (mESCs) harboring a previously delivered Landing Pad. In this case, the native locus is deleted and replaced with a marker cassette (MC1) acting as a Landing Pad that encodes positive and negative selection markers, and a fluorescent tag. Given that mSwAP-In relies on homologous recombination (HR) after Cas9-mediated double-strand break, a conserved mechanism across different mammalian systems, here we extend the mSwAP-In technology to hiPSCs.

Using this platform, we produced a series of isogenic hiPSC lines consisting of a parental cell line, a gene cluster knockout cell line, and a series of synthetic haplotype-containing cell lines at the *NKX6-3/ANK1* T2D GWAS locus. The intent of this approach is to ultimately dissect the locus at a single-base-pair resolution. By performing this work in hiPSCs, we were able to differentiate cells harboring the various natural and synthetic haplotypes into T2D-relevant cell types and measure how each haplotype might affect gene expression and/or differentiation potential towards disease- and locus-relevant cell types. This workflow can serve as a model for functional and experimental dissection of important GWAS disease loci.

## Results

### Identifying Candidate Risk SNPs at the *NKX6-3/ANK1* locus

To determine informative haplotypes to build, we combined data from European and East Asian GWAS and European eQTL for both *ANK1* and *NKX6-3* and identified SNPs that are in high LD (*r^2^ >* 0.6) with the lead SNPs representing two independent signals at the locus, denoted in asterisks (Fig. 1*A* *SI Appendix,* Tables S1-S2). SNPs that fall into high LD in all three categories (European GWAS, East Asian GWAS, and European eQTL) were selected, consistent with the hypothesis that the causal SNP(s) of the locus must be located within the intersection. A total of 38 SNPs were identified across ∼33 kb, with half colocalizing with an *NKX6-3* pancreatic islet eQTL signal, and the other half with an *ANK1* skeletal muscle eQTL signal (D’ = 0.956, R² = 0.072 between two GWAS signal lead SNPs) (Fig. 1*A*, *SI Appendix,* Table S1). We also performed cross-tissue annotations in pancreatic islets and skeletal muscle to identify promoter activity and chromatin accessibility signatures at each SNP by comparing open chromatin and chromatin states in the respective tissues (Fig. S*1A-D*).

**Figure 1:**
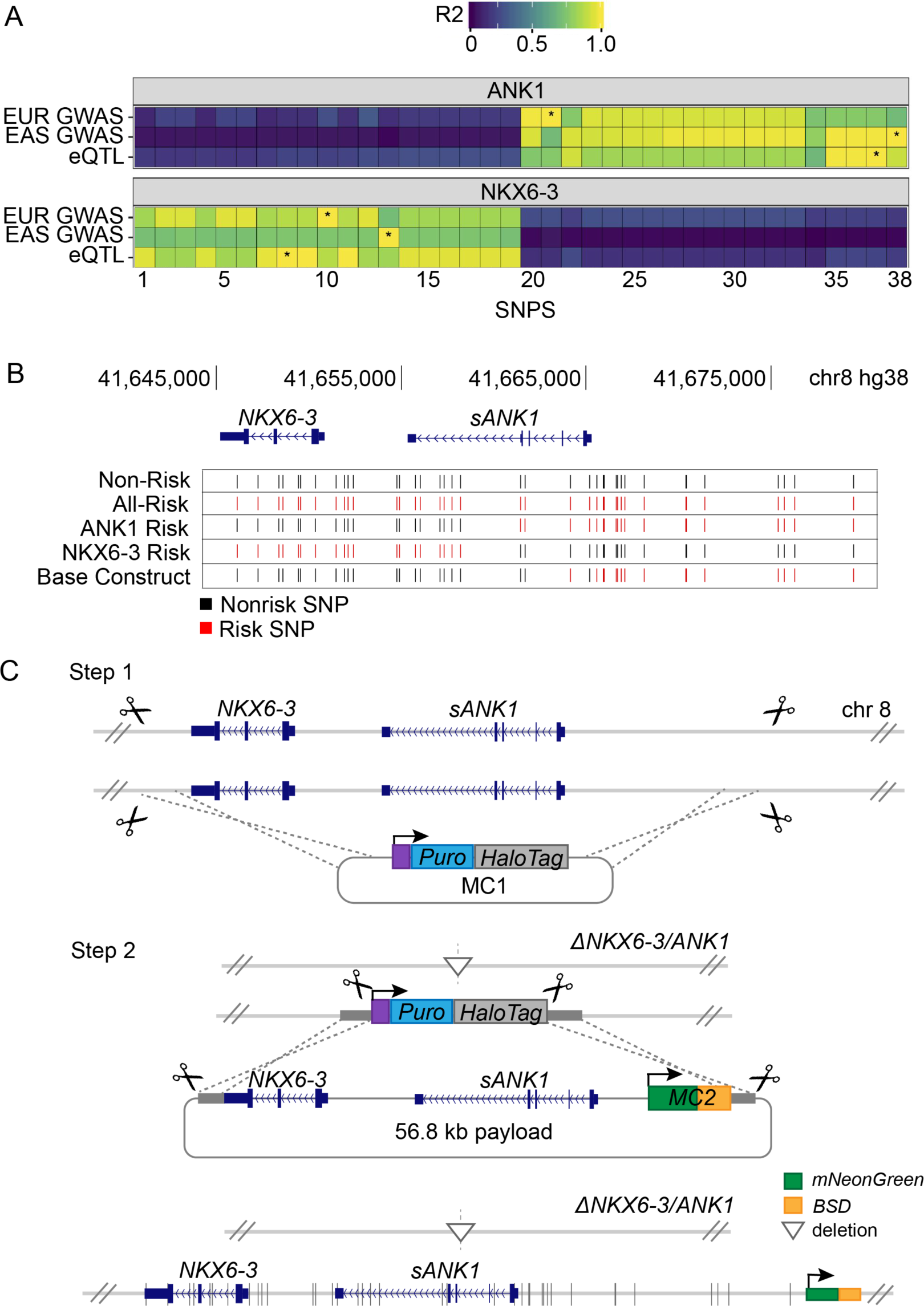
*NKX6-3/ANK1* cluster haplotypes. **A.** Data from European and East Asian GWAS and eQTL studies were combined to identify SNPs with *r>0.5* of the lead SNPs (*). A total of 38 SNPs were identified, half of which colocalize to *ANK1,* and the other half colocalizing with *NKX6-3*. A full list of SNPs can be found in *SI appendix* Table S1. The two sets of risk SNPs are defined in this paper as the *NKX6-3* risk SNPs (left side) and the *ANK1* risk SNPs (right side). **B**. Genome Browser snapshot of SNP locations and the series of isogenic haplotypes that were built: Non-Risk, All-Risk, ANK1-Risk, NKX6-3-Risk, and Base Construct. Risk SNPs are denoted in red, and nonrisk SNPs are denoted in black. Base construct reflects reference genome. **C**. Diagram depicting overall scheme for mSwAP-In in hiPSCs at the *NKX6-3/ANK1* locus. Step 1 uses two Cas9-gRNAs to delete the endogenous locus and replace it with a marker cassette (MC1) containing *PuroR-HaloTag*. Step 2 shows the delivery of a 56.8-kb haplotype payload (68 kb total plasmid size) containing *NKX6-3, ANK1*, and a second marker cassette (MC2) containing *mNeonGreen* (green box) and blasticidin resistance gene (orange box) along with Cas9-gRNAs that target MC1 and the payload. The inverted triangle symbolizes the NHEJ-mediated deletion of the *NKX6-3/ANK1* cluster in one allele.

Based on the above bioinformatic analyses, we developed and implemented a strategy to design, build, and deliver several initial synthetic haplotypes: All Risk, Non-Risk, *ANK1* (only) risk SNPs (referred to as ANK1-Risk), and *NKX6-3* (only) risk SNPs (referred to as NKX6-3-Risk), as well as a Base Construct that allowed us to engineer the synthetic haplotypes. The haplotypes have the following haplotype frequencies: All-Risk haplotype freq. = 0.517; Non-Risk haplotype freq. = 0.156; the ANK1-Risk and NKX6-3-Risk haplotypes are rare or unobserved in reference populations (Fig. 1*B*, *SI Appendix*, Tables S1-S2). These synthetic haplotypes will be delivered to the endogenous *NKX6-3/ANK1* locus in hiPSCs using a two-step mSwAP-In approach so as to ensure any *NKX6-3/ANK1* signal is coming from the integrated synthetic haplotypes (Fig. 1*C*, Steps 1–2), which will be further described in the following sections.

### Building Haplotype Payloads

To build the synthetic haplotype payloads, a 52.8-kb segment from a bacterial artificial chromosome (BAC) containing human *NKX6-3/ANK1* genomic sequence was digested *in vitro* using CRISPR-Cas9 RNPs (21, 22) and verified by pulsed-field gel electrophoresis (*SI Appendix*, Fig. S2*A*). The digested BAC was transformed into Yeast Assemblon Vector (YAV) pLM1050, which can be shuttled between yeast and *E. coli*, using linker DNA to direct yeast HR (18, 19, 23, 24) (Fig. *1B*, Fig. *2A*). The yeast transformants were screened via junction PCR, and 54% were found to have both junctions, suggesting efficient assembly of the digested BAC and assembly vector. (Fig. 2*B*, Fig. S2*B*). Four such clones chosen at random were recovered in *E. coli* for plasmid extraction and sequencing verification, and all were found to contain the 52.8-kb BAC segment with no sequence errors (Fig. 2*C*). A secondary marker cassette, MC2, was subsequently inserted ∼1.6 kb upstream from the 3’ end of the digested BAC using CRISPR-Cas9 (25), and sequence-verified (Fig. S2*C*-S*2D*). The resulting payload is referred to as “Base Construct,” and shares the same genotype as the reference genome (Fig. 1*B*, Fig. *2A*).

**Figure 2:**
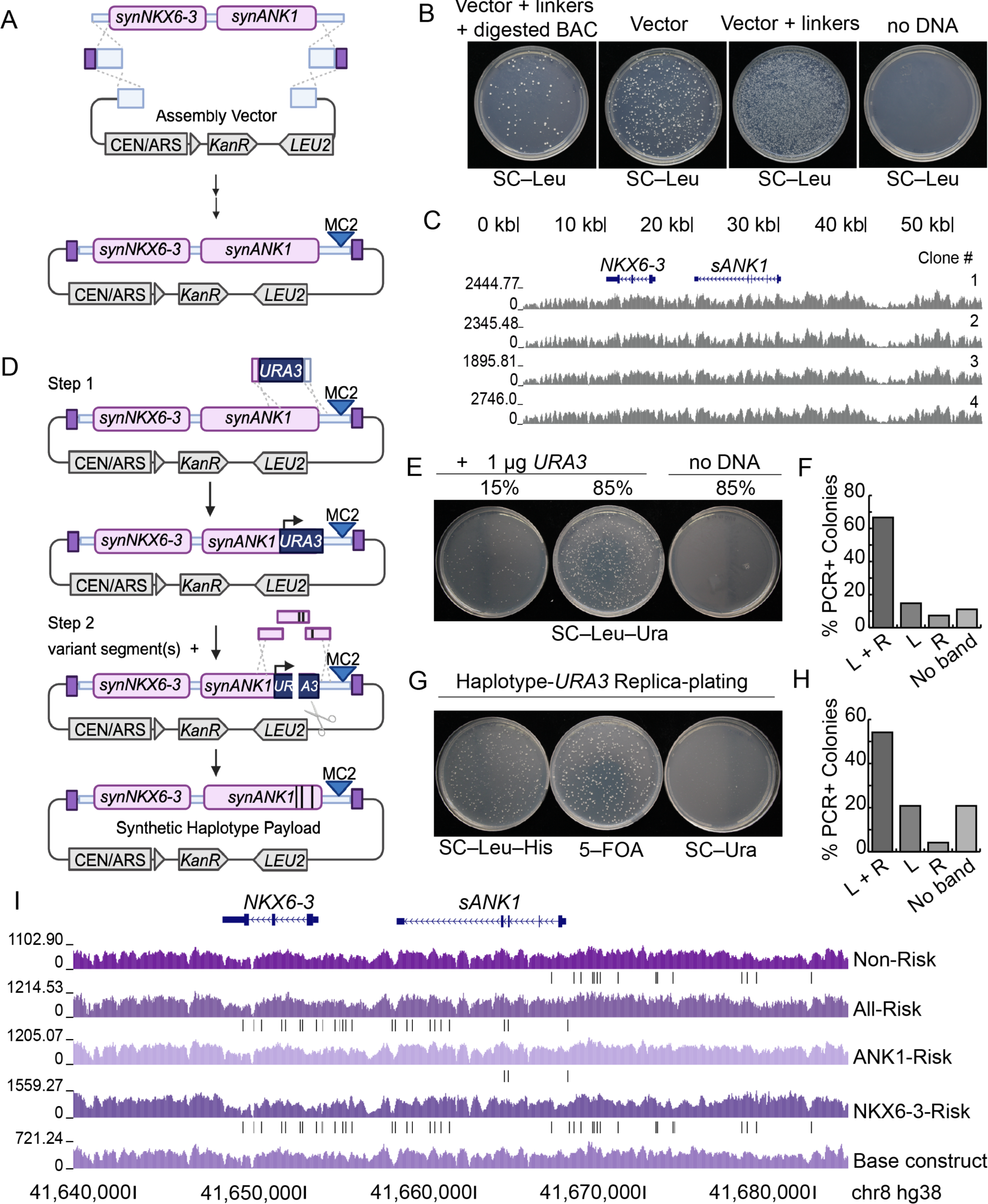
Haplotype building. **A.** Base Construct assembly performed by digesting BAC RP11-111B9 using *in-vitro* CRISPR digestion, and transforming into yeast along with linker DNAs containing homology to BAC digest and a linearized assembly vector. MC2 was then cloned towards the 3’ end of the digested BAC. **B.** Yeast transformation of Base Construct performed with all components, vector only, vector and linker DNA, and no DNA. 26 of 48 colonies tested from the “complete” plate had the expected base construct structure. **C.** Sequencing coverage of Base Construct payload clones spanning ∼52 kilobases of the *NKX6-3/ANK1* locus **D.** Diagram depicting vSwAP-In method. First, a *URA3* segment containing > 40 bp of flanking terminal homologies replaces the region to be edited in the haplotype payload. Next, variant segments are transformed, swapping out the *URA3* marker. **E.** vSwAP-In transformation plates showing the *URA3* was successfully inserted in the payload plasmid by selective plating on SC–Leu–Ura medium. **F.** Amplicon analysis. Percentage of colonies screened (n=27) at random for left (L) and right (R) junctions of *URA3* transformants after single-colony purification. **G.** vSwAP-In amplicon analysis. Post-transformation of variant segments, many yeast colonies survived on 5-FOA, showing high *URA3* loss efficiency. **H.** Percentage of vSwAP-In colonies screened (n=24) containing left (L), right (R), or no junctions of vSwAP-In transformants after single-colony purification. **I.** Sequencing coverage of synthetic haplotype payloads containing various risk and non-risk SNPs built using a combination of vSwAP-In and *in yeasto* CRISPR-Cas9 editing methods. Black vertical lines mark SNPs. A full list of SNPs present in the haplotype payloads can be found in *SI Appendix* Tables S5-8.

To create the various haplotypes, we developed and deployed a modified, *in yeasto* plasmid-building method called vSwAP-In, which relies on yeast HR to seamlessly integrate DNA variants in a targeted region of interest (Fig. 2*D*). The vSwAP-In method is a two-step editing process that uses positive/negative selectable auxotrophic markers such as *URA3* to select for colonies that have successfully integrated or lost DNA segments, based on a previously developed method for engineering yeast extrachromosomal DNA (18). First, *URA3* flanked by 40-60 bp of terminal homologies to the region of interest is used to replace the wild-type region on the payload (Fig. 2*D*, Step 1, Fig. 2*E*–2*F*). Next, variant segments are co-transformed with a *URA3*-targeting Cas9 expression plasmid to then replace the *URA3* (Fig. 2*D*, Step 2). Transformants are selected for *URA3* loss using 5–FOA media (26) and further confirmed to have lost *URA3* on SC–Ura (Fig. 2*G*). Transformants are then PCR-screened for *URA3* junctions (Fig. 2*H*, *SI Appendix* Table S4). The haplotype payloads were constructed using this vSwAP-In method to make the majority of SNP changes, and *in yeasto,* CRISPR-based editing to introduce minor SNP changes (24, 25, 27). vSwAP-In was found to be successful, and relatively efficient, with roughly 50–65% of screened clones containing correct junctions for both *URA3* transformations and variant segment transformations (Fig. 2*F*, Fig. 2*H*). Several *URA3*-containing precursors, and haplotypes harboring internal deletions were made during the construction of the four main haplotype payloads, but were not used for downstream analyses (Fig. S3). The constructed haplotype payloads were then recovered into *E. coli* and sequence verified for correct SNP configurations (Fig. 2*I*). A list of SNPs present in the All-Risk, ANK1-Risk, NKX6-3-Risk, and Non-Risk haplotypes relative to the reference genome (Fig. 2*I*, Fig. 1*B*) can be found in *SI Appendix* Tables S5–S8.

### Establishing and characterizing MC1 Landing pad hiPSCs with *NKX6-3/ANK1* deletion

To deliver haplotype payloads into hiPSCs, a marker cassette (MC1) was inserted so as to eliminate the endogenous *NKX6-3/ANK1* locus, ensuring that upon synthetic haplotype integration, *NKX6-3/ANK1* transcription is solely from the synthetic haplotypes harboring the various risk SNP combinations. To establish the hiPSC precursor cell line (*NKX6-3^-/-^ ANK1^-/-^*) for haplotype building, we inserted an MC1 landing pad at the *NKX6-3/ANK1* locus by following a general transfection and selection timeline (Fig. 3*A*). MC1 donor DNA was modified to contain flanking homologies to the *NKX6-3/ANK1* locus, and two Cas9-gRNA plasmids were cloned to target low-activity regions based on ATAC-seq data from islets and skeletal muscle tissue (19) (Fig. 1*C*, Step 1), Fig. S4*A*). The MC1 and the two Cas9-gRNAs plasmids were transfected into AS0041 hiPSC parental cells, and selected with an antibiotic for 6 days (Fig. S4*B*-S4*C*). The cells were incubated with the HaloTag ligand Janelia Fluor® 549 (JF549) to visualize established colonies and allow for single-cell sorting of cells with MC1 integrated (Fig. 3*B*). Of the cells transfected with both the MC1 and the two Cas9-gRNAs plasmids, 37.3% were found to be JF549+ compared to 23.45% when only MC1 was transfected (Fig. 3*C*, Fig. S5*A*), indicating a ∼2 fold enrichment of correctly integrated MC1. Out of 288 cells that were single-cell sorted, 93 clones successfully grew after undergoing another round of antibiotic selection, and one of the 93 clones was found to contain both left and right junctions by PCR genotyping and to lack *NKX6-3/ANK1* (Fig. S5*B*). This clone (#73, hereinafter referred to as AS0041^MC1^) was verified by targeted Capture sequencing (Capture-Seq) (22, 28), and was confirmed to have lost *NKX6-3/ANK1* in both alleles, suggesting either hemizygous or biallelic integration of MC1 (Fig. 3*D*). To determine zygosity, the entirety of the MC1 cassette was amplified from AS0041^MC1^ genomic DNA with primers that bind the flanking genomic regions. Two distinct amplicons were detected: one at ∼5 kb representing full-length, single-copy correctly integrated MC1, and one at ∼2 kb, consistent with a genomic deletion of the region between the two gRNA targets (Fig. S5*C).* Sanger sequencing confirmed that the latter amplicon corresponded to a complete deletion of the *NKX6-3/ANK1* locus initiated by the cutting by Cas9-gRNA and deletion formation via non-homologous end joining (Fig. S5*D).* The deletion of the *NKX6-3/ANK1* locus in the AS0041^MC1^ cell line did not have a noticeable effect on cell growth, which is unsurprising given that *NKX6-3* and *ANK1* are not detectably expressed in human embryonic stem cells, which are similar to hiPSCs.

**Figure 3:**
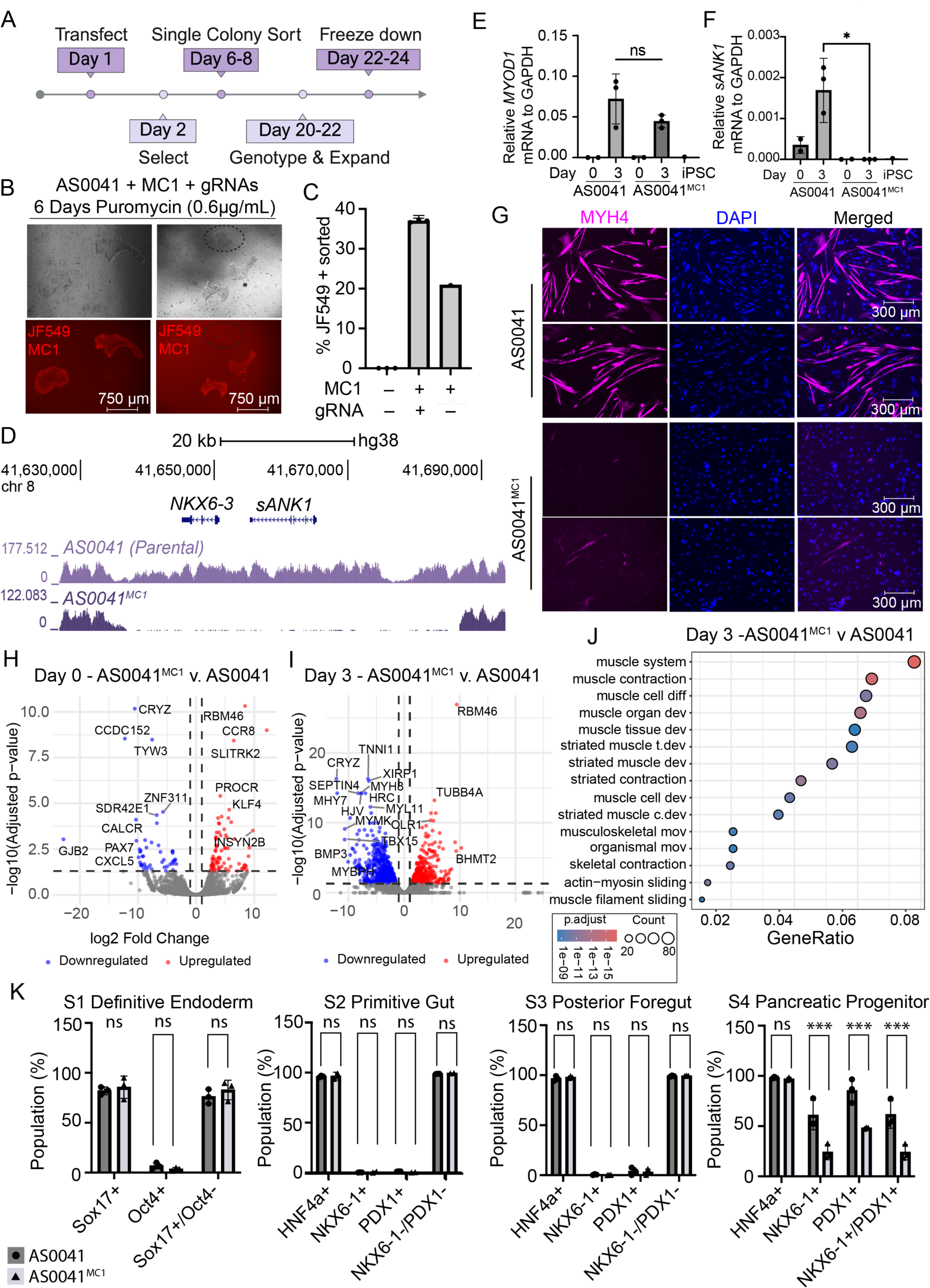
*NKX6-3/ANK1* Cluster knockout line. **A.** Timeline for establishing a landing pad hiPSC line. **B.** Images of AS0041 hiPSCs transfected with an MC1 plasmid after 6 days of puromycin selection (0.6 µg/mL). Cells were incubated with the JF549 HaloTag ligand for 1 hour prior to imaging and subsequent flow sorting. **C**. Percentages of MC1-harboring (JF549+) AS0041 hiPSCs following transfection with an MC1 plasmid in the presence or absence of a Cas9-gRNAs expression plasmid. JF549+ hiPSCs were single-cell sorted into three 96-well plates to establish monoclonal cell populations. **D.** Capture-sequencing reads mapped to hg38. Results show a complete deletion of *NKX6-3/ANK1* from AS0041^MC1^. **E.** Immunofluorescence of AS0041 and AS0041^MC1^ hiPSCs following skeletal muscle differentiation. Cells were stained for MYH4, a late-stage skeletal muscle marker, and DAPI for nuclei. **F and G.** qRT-PCR analysis of *MYOD1* (F) and *sANK1* (G) for the indicated hiPSC lines at days 0 and 3 of skeletal muscle differentiation. Expression was normalized to *GAPDH* mRNA level. **H.** Volcano plot of differentially expressed genes at Day 0 prior to differentiation of AS0041^MC1^ compared to parental cell line. Early stage muscle cell transcription factors are downregulated, while those that promote skeletal muscle development were upregulated. Data are plotted with adjusted p-value cut-off of 0.05 and absolute log2fold change of 1. **I.** Volcano plot of differentially expressed genes 3 days post-differentiation of AS0041^MC1^ compared to parental cell line. Genes that play important roles in late-stage skeletal muscle development were downregulated, while genes involved in metabolism were upregulated. Data are plotted with adjusted p-value cut-off of 0.05 and absolute log2fold change of 1. **J.** Gene ontology (GO) Enrichment analysis of biological processes of AS0041^MC1^ compared to parental cell line 6 days post-differentiation. Genes involved in muscle development and processes, and structural function were enriched as a result of *NKX6-3/ANK1* deletion, plotted by the proportion of input genes annotated to the GO term by the total number of input genes (GeneRatio). **K**. Flow cytometry analysis of percentage of AS0041 and AS0041^MC1^ cells expressing various differentiation markers for each stage of pancreatic progenitor differentiation. No significant differences between the two lines were observed except at Stage 4, where AS0041^MC1^ showed a significant decrease in NKX6-1+ and PDX1+ cells, *p* = 0.0006 (NKX6-1+); 0.0005 (PDX1+); 0.0004 (NKX6-1+/PDX1+). *P* values were calculated using a 2-way ANOVA (Šídák’s multiple comparisons test).

Next, we evaluated the effect of deleting *NKX6-3/ANK1* on muscle differentiation by infecting both AS0041^MC1^ and the parental cell line, AS0041, with a lentivirus overexpressing *MYOD1* (29). Cells were cultured in skeletal muscle medium containing a cocktail of inhibitors to promote commitment to the myogenic progenitor state until they were ready for *MYOD1*-lentiviral infection. Prior to infection, cells were collected as “Day 0”. Cells were also collected on “Day 3” and “Day 6” post *MYOD1* lentiviral infection. Quantitative measurements of both *MYOD1* and *sANK1* (short *ANK1*) mRNA levels showed that AS0041^MC1^ cells did not express *sANK1* but expressed comparable levels of *MYOD1* compared to AS0041 (Fig. 3*F-G*). At the end of the muscle differentiation, AS0041 and AS0041^MC1^ cells were stained with myosin heavy-chain 4 (MYH4), a late-stage muscle marker. We found that AS0041^MC1^ failed to properly differentiate and form myotubes, as expected, because *sANK1* is known to play an early role in the organization of the sarcoplasmic reticulum during skeletal muscle development to create skeletal muscle fibers (30) (Fig. 3*E*). We also constructed single gene deletions for *NKX6-3* and *sANK1* and we showed that whereas the *sANK1* deletion failed to differentiate into muscle cells, *NKX6-3* deletion cells differentiated normally into muscle cells (not shown).

In addition to the role *sANK1* plays in the organization of skeletal muscle development, some studies show that *sANK1* expression is directly regulated by MYOD, as *sANK1*’s alternative promoter contains two conserved MYOD binding sites, potentially defining an important *sANK1* enhancer (31–33). To gain deeper insight into global gene expression changes, bulk-RNA sequencing was performed on cells collected at Days 0, 3, and 6 of differentiation. Genes crucial for early stages of skeletal muscle differentiation, *PAX7, CALCR,* and *KLF4,* were seen to be differentially expressed on Day 0 in AS0041^MC1^ compared to AS0041(31, 34, 35) (Fig. 3*H*). Both *PAX7* and *CALCR* maintain the skeletal muscle stem cell state, and their downregulation suggests decreased muscle stemness; however, the upregulation of *KLF4* suggests enhanced myogenic progenitor proliferation. Three days and six days post differentiation via *MYOD1* overexpression resulted in differential expression of genes involved in mid-stage and late-stage muscle development, such as the myosin heavy and light chain family (*MYH8, MYH7*, and *MYL11*), myomaker *(MYMK)*, RNA-binding protein *MBNL3,* and other genes related to muscle structure and function and such as *SEPTIN4,* myosin binding protein-H (*MYBPH*), and T-box transcription factor-15 (*TBX15)* (36–40) (Fig. 3*I*, Fig. S7*A*). Other notable differentially expressed genes observed were some related to insulin signaling, such as *DUSP9 and APOB,* important for glucose uptake within muscle cells (41, 42), and calcium ion regulation, such as *SRL* and *HRC*, important for muscle development and regeneration (43, 44) (Fig. S7*A*). A Gene Ontology (GO) analysis was performed to identify the biological processes in which these differentially expressed genes play important roles. Cells from Days 3 and 6 showed genes enriched specifically in skeletal muscle processes and development, suggesting that the loss of *NKX6-3/ANK1,* and presumably *sANK1* due to its known role in skeletal muscle, is critical for proper muscle differentiation and function (Fig. 3*J*, Fig. S7*B*).

The *NKX6-3* gene is a part of the NKX family of homeodomain transcription factors responsible for numerous developmental processes in the central nervous system and digestive and endocrine systems (45–49). Closely related *NKX6-1* plays an important role in the formation and maintenance of insulin-producing beta cells (50–52), and some studies suggest *NKX6-3* is expressed in islets (14, 53, 54), though its role is not understood. To investigate *NKX6-3*’s role in pancreatic cells, we differentiated AS0041 and AS0041^MC1^ into pancreatic progenitors (see Materials and Methods), and marker-positive cell populations were assessed by flow cytometry. We observed that both cell lines showed comparable levels of SOX17 and OCT4 in the definitive endoderm stage (Stage 1). However, by the pancreatic progenitor stage (Stage 4), AS0041^MC1^ cells show significantly lower levels of NKX6-1 and PDX1 compared to parental cells (*p* = 0.0004), suggesting that *NKX6-3/ANK1* locus may be critical for the transition into pancreatic progenitor cell types (Fig. 3*K*). We attempted to quantify *NKX6-3* expression in these cells at the various stages via RT-qPCR, but we were unable to detect *NKX6-3* mRNA at any of the stages (Fig. S8*A-D*), and neither in adult islets nor β-cell organoid samples (Fig. S8*E-G*).

### Establishing an Isogenic Series of Synthetic Haplotype iPSC lines

To test the various haplotypes for T2D risk, we delivered the haplotypes (Fig. 2*I*) to AS0041^MC1^ cells using mSwAP-In (19). The haplotype payloads were equipped with a second marker cassette (MC2) containing a blasticidin selection and mNeonGreen fluorescence marker cassette to be “swapped” with MC1. After 6 days of blasticidin selection, round, polyclonal colonies formed, containing a mix of cells with integrated MC2 and those without (Fig. 4*A*, Fig. S9*A-C*). Presumably, the residual MC1-containing cells persisted due to reduced drug penetration as a result of colony-based protection. We first used nucleofection to integrate the haplotype payloads. To purify the cells that had an MC1 to MC2 swap, hiPSCs were single-cell sorted to ensure monoclonality and then screened via junction PCR to identify clones that successfully integrated the synthetic *NKX6-3/ANK1* haplotype payloads. Using nucleofection methods, 26.03% of transfected cells were mNeonGreen+, and 20% of PCR-screened colonies post single-cell sorting contained both left and right payload junctions (Fig. S9*D-F*). We then compared nucleofection and lipofection methods, and found that lipofection methods resulted in more mNeonGreen+ cells than nucleofection (Fig. *S10A-B*). Single-cell sorting of lipofection and nucleofection conditions confirmed that there was a higher percentage of mNeonGreen+ cells in the lipofection condition (40-46%) as compared to nucleofection (29-30%) (Fig. S10*C*-S10*D*). Candidate clones were also screened for left and right PCR junctions, with 45-50% lipofected candidates with left and right junctions, and 25-50% of nucleofected candidates with left and right junctions. The lipofection method was used to integrate the remaining haplotype payloads (Fig. S11*A-F*). Prior to sequencing, candidate clones from each haplotype underwent repeated PCR screening to ensure MC1 loss and lack of backbone integration (Fig. S12). Candidate clones were further verified via Capture-seq (22, 28) to ensure integration of synthetic *NKX6-3/ANK1* haplotype payloads in the correct genomic context (Fig. 4*B*).

**Figure 4:**
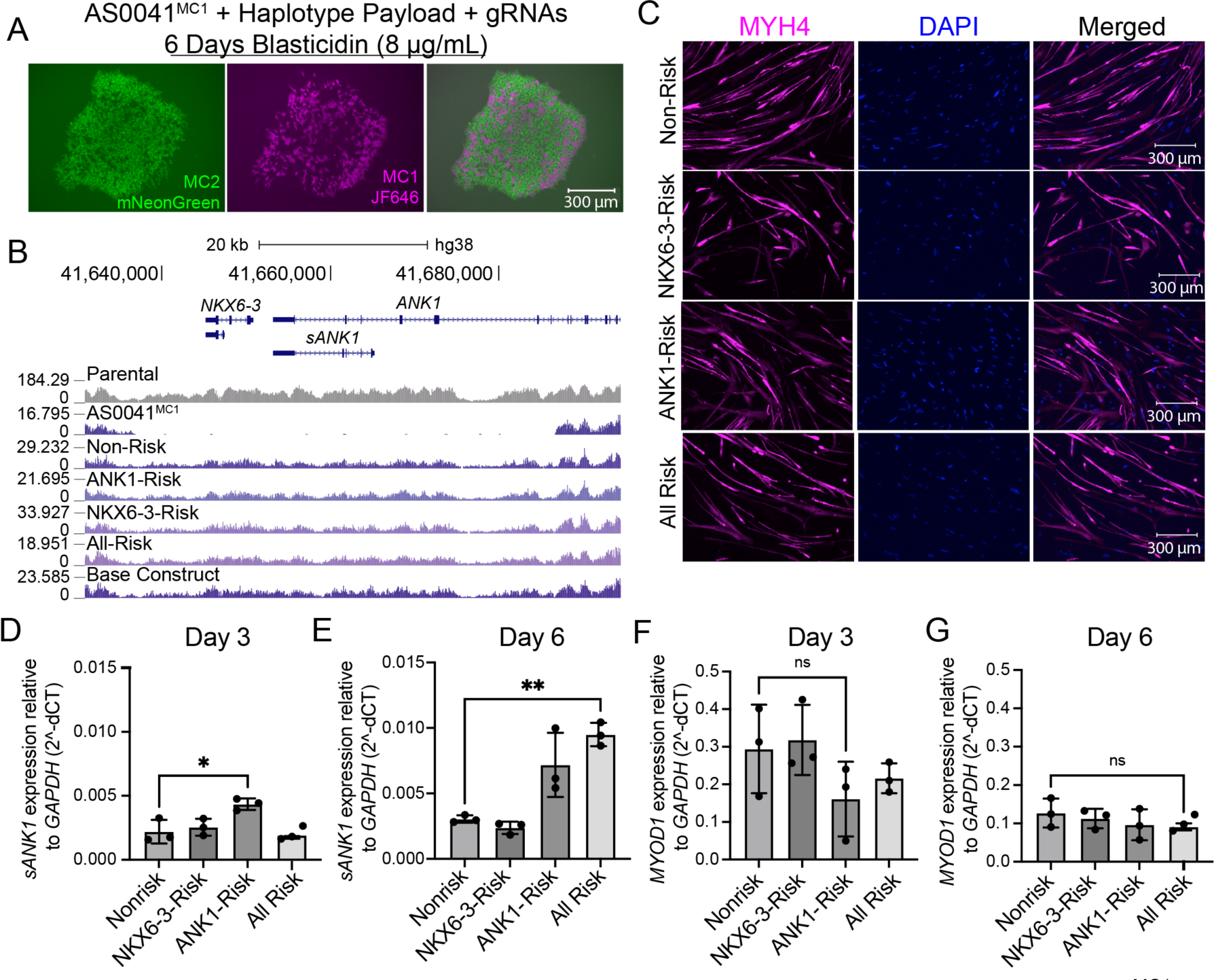
An isogenic set of hiPSCs with a series of synthetic haplotypes. **A**. AS0041^MC1^ cells transfected with a haplotype payload and two Cas9-gRNA expression plasmids after 6 days of blasticidin selection. Cells that did not integrate the payload remain tagged with JF646 HaloTag Ligand, while cells that integrated the MC2-containing payload DNA gain mNeonGreen expression. **B.** Capture-seq data demonstrating successfully integrated haplotype payloads into AS0041^MC1^ cells with reads aligned to the hg38 human genome, showing coverage consistent with hemizygous reintegration of *NKX6-3/ANK1* locus. **C.** Mature myotubes 14-days post-MYOD1 lentiviral infection stained with MYH4 to identify late-stage skeletal muscle cells, and DAPI to identify nuclei. **D.** qRT-PCR measurements of *ANK1* 3 days post lentivirus infection show ANK1-Risk haplotype to have higher expression level compared to Non-Risk (*p* = 0.0388, Student’s t-test). Each dot represents a technical replicate of the differentiation. **E.** qRT-PCR measurements of *ANK1* 6 days post lentivirus infection show All-Risk haplotype to have higher expression levels compared to Non-Risk (*p* = 0.0034, Student’s t-test) Each dot represents a technical replicate of the differentiation. **F-G.** qPCR measurements of *MYOD1* 3 and 6 days post-infection showed no significant expression differences between haplotypes. Each dot represents a technical replicate of the differentiation.

To assess the different T2D risk allele combinations, the haplotype hiPSCs were differentiated into skeletal muscle cells and pancreatic progenitors. Notably, the newly delivered synthetic DNAs restored both pancreatic and muscle phenotypes that were lost in the AS0041^MC1^ knockout cell line, proving that these phenotypes are encoded on the synthetic haplotypes delivered and did not arise from unrelated clonal variability. There were no apparent qualitative differences in skeletal muscle differentiation capabilities, regardless of haplotype (Fig. 4*C*). Similarly, we performed pilot pancreatic progenitor differentiations on the All Risk and *NKX6-3* Risk haplotypes and observed that reintroduction of *NKX6-3/ANK1* haplotypes restored levels of pancreatic progenitor transcription factors NKX6-1+ and PDX1+ cells to levels similar to the parental cell line, as seen in Stage 4 pancreatic progenitor cells (Fig. S13*A-D)*.

According to eQTL studies, *sANK1-*associated T2D risk alleles correlate with increased expression of *sANK1* (9, 12); however, no functional studies have been performed with these risk SNPs together in their native context. To see how these different risk SNP combinations contribute to increased T2D risk, we examined expression of *sANK1* across the various haplotypes. We first measured *sANK1* mRNA levels at different stages of myogenic differentiation, and found that *sANK1* was most highly expressed at Day 6 (Fig. S13*E*). We then collected cells at Days 0, 3, and 6, and found that the All Risk and ANK1*-*Risk haplotypes showed increased levels of *sANK1* expression (ANK1-Risk Day 3: *p* = 0.038; All-Risk Day 6: *p =* 0.0034) compared to the Non-Risk haplotype at Days 3 and 6 (Fig. 4*D-E*, Fig S13*F*), despite expressing similar levels of *MYOD1* (Fig 4*F-G*, Fig S13*G*). These findings are consistent with T2D GWAS and eQTL studies previously reported at this locus.

### Characterization of Risk compared to Non-Risk Haplotypes for T2D-associated phenotypes

To further investigate the impact T2D risk SNPs have on the transcriptional landscape of the differentiated skeletal muscle cells, we conducted RNA-sequencing on each haplotype at Days 0, 3, and 6. We first examined gene expression trajectories throughout differentiation to identify patterns between the All-Risk, ANK1-Risk, and NKX6-3-Risk haplotypes compared to the Non-Risk haplotype. *ANK1* expression level was found to increase from Day 0 to Day 3, and slightly drop in expression at Day 6 (Fig. 5*A*). The ANK1-Risk haplotype showed higher *ANK1* expression compared to the Non-Risk haplotype (*p* = 0.0497), supporting previous findings (9, 12). Additionally, *ANK1* expression level remained high at Day 6 in ANK1-Risk and All-Risk haplotypes compared to the NKX6-3-Risk and Non-Risk haplotypes. Interestingly, the All-Risk haplotype expressed significantly higher early-stage myogenic transcription factors *PAX3* and *PAX7* genes at Days 0 and 3 compared to all the other haplotypes (*PAX7: p* = 0.024 (Non-Risk), 0.036 (NKX6-3-Risk), 0.047 (ANK1-Risk); *PAX3*: *p* = 0.036 (Non-Risk), 0.021 (ANK1-Risk) (Fig. 5*B-C*). The expression level of these genes decreases after Day 3, but still remains higher than the other haplotypes on Day 6. Consistent with RT-qPCR data (Fig. 4*F-G*), all haplotypes expressed similar levels of *MYOD1* on Days 3 and 6 (Fig. 5*D*), as well as *MYOG*, an essential skeletal muscle transcription factor that codes for myogenin (Fig. 5*E*). Finally, the All-Risk vs Non-Risk haplotype comparison revealed the largest differences among the haplotype comparisons done, and thus we focused analyses on this.

**Figure 5.**
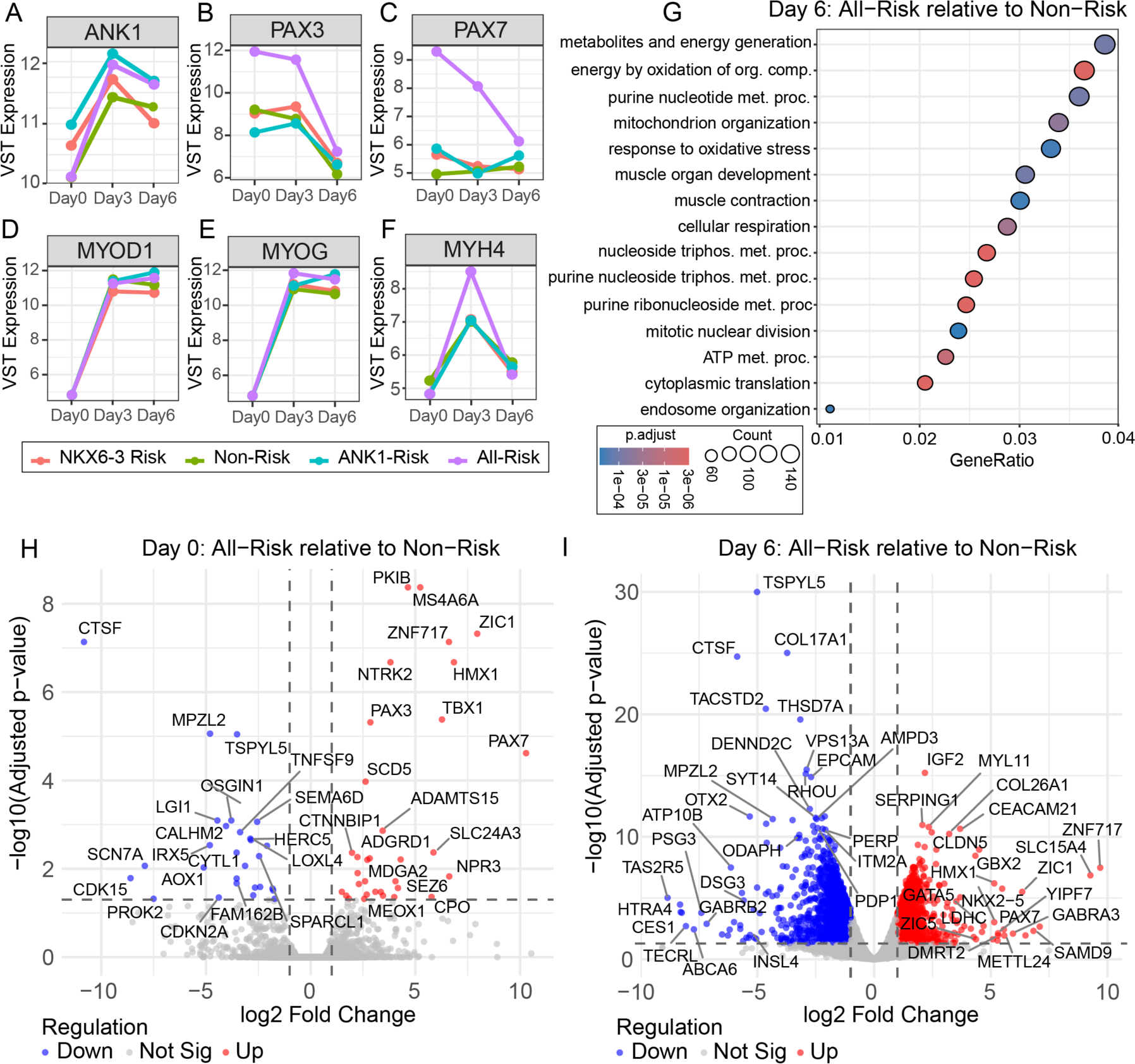
Gene expression changes specific to All-Risk vs Non-Risk haplotypes: A-F. Variance stabilizing transformation (VST) expression line graphs of select myogenic-related genes throughout differentiation on Days 0, 3, and 6 NKX6-3-Risk (red), Non-Risk (green), ANK1-Risk (blue), and All-Risk (purple) haplotypes. *P* values were calculated using a 2-way ANOVA. *ANK1*: *p* = 0.0497 (Non-Risk vs. ANK1-Risk); *PAX3*: *p* = 0.036 (Non-Risk vs. All-Risk), 0.021 (ANK1-Risk vs. All-Risk); *PAX7: p* = 0.024 (Non-Risk vs. All-Risk), 0.036 (NKX6-3-Risk vs. All-Risk), 0.047 (ANK1-Risk vs. All-Risk). **G.** Gene Ontology (GO) analysis for biological processes of the All-Risk haplotype compared to the Non-Risk haplotype at Day 6 of differentiation, showing genes enriched in various skeletal muscle functions and energy metabolism plotted by the proportion of input genes annotated to the GO term by the total number of input genes (GeneRatio). Adjusted p-value cut-off set to 0.05, and absolute log-fold change cut off set to 1. Abbreviations are as follows: org. = organic; comp. = compounds; met. = metabolic; proc. = process. **H-I.** Volcano plot showing differentially expressed genes at Day 0 (**H**) and Day 6 post *MYOD1* lentiviral infection (**I**) between the All-Risk haplotype vs. the Non-Risk haplotype, prior to infection with *MYOD1*. Downregulated genes (blue) and upregulated genes (red) plotted with adjusted p-value cut-off of 0.05 and absolute log2fold change of 1.

Additional myogenic gene expression trajectories of structural and functional genes such as myosin heavy chain genes (MYHs) and myosin light chain genes (MYLs) were also examined. The majority of them did not show temporal differences in gene expression levels or patterns among the haplotypes. However, the gene expression trajectories of *MYH1*, *MYL6*, and *MYL9*, showed differences between all of the risk-SNP containing haplotypes compared to the Non-Risk haplotype (Fig S14*A*, S14*B*). For *MYH1*, the Non-Risk haplotype shows a steady increase of gene expression level between Days 0 and 6, whereas the other haplotypes show a stark pattern of expression difference, with gene expression level going up from Days 0 to 3, and then down at Day 6 (Fig S14*A*). Expression trajectories for *MYL6* showed all of the risk SNP-containing haplotypes with increased gene expression throughout the differentiation, while the Non-Risk haplotype had a slight decrease in expression at Day 6 (Fig. S14*B*). In addition to *MYH* and *MYL* genes, we also examined several structural proteins of the sarcomere, including *TNNT1*, *TNNT2*, *ACTA1*, and *DES*, as the sarcomere plays an important role in stabilizing the muscle during contractile function (55). While there were no apparent differences identified in the expression of *TNNT1*, *TNNT2*, and *DES*, a very dramatic difference was observed in *ACTA1* gene expression, with the All-Risk haplotype showing delayed expression until Day 6 (Fig. S14*C*). These findings suggest that the risk-associated SNPs indeed influence the transcriptional landscape of a set of functional genes in skeletal muscle.

To assess the functional consequences of the haplotypes on insulin signaling, we examined the temporal expression of key pathway components during skeletal muscle differentiation. The All-Risk haplotype exhibited altered expression of multiple insulin signaling genes, including reduced *IRS1* expression at Days 3 and 6 (Fig.S15*A*), but no difference in *IRS2* expression (Fig. S15*B*). The All-Risk haplotype also showed signficant expression pattern differences of PI3K and AKT signaling pathway genes compared to the Non-Risk haplotype (Fig. S15*C-F*) (*PIK3CA*: *p*-value = 0.0217; *PIK3CB*: *p*-value = 0.0252; *AKT1*: *p*-value = 0.0346). *SLC2A4* (GLUT4) expression followed similar temporal patterns across the haplotypes at Days 3 and 6, but the All-Risk haplotype exhibited lower gene expression on Day 0 compared to the other haplotypes (Fig. S15*G*). Unexpectedly, *GSK3B*, which encodes glycogen synthase kinase 3B and is typically regulated by *AKT*, was reduced at Day 6 in the All-Risk and ANK1-Risk haplotypes compared to the Non-Risk and NKX6-3-Risk haplotypes (56, 57) (Fig. S15*H*). Together, these data indicate that risk SNPs across the *NKX6-3/ANK1* locus are associated with alterations in insulin signaling gene regulation.

To better understand the differences between the All-Risk and Non-Risk haplotype, we performed a GO enrichment analysis on Day 6 samples, where the differences were most prominent. All-Risk cells were enriched for biological processes related to energy metabolism, mitochondrial function, oxidation of organic compounds, and muscle organ development and contraction (Fig. *5G*). We also plotted differentially expressed genes of the All-Risk haplotype relative to the Non-Risk haplotype on Day 0 and Day 6. At Day 0, All-Risk cells in the myogenic progenitor state (Day 0) exhibited upregulation in early-stage myogenic genes *PAX3* and *PAX7* compared to the Non-Risk haplotype (Fig. 5*H*). *PAX3* and *PAX7* work with other myogenic regulatory factors (MRFs) such as *MYOD1*, *MYOG*, *MYF5* (myosin factor 5), and *MRF4* to promote the differentiation of progenitor cells into mature skeletal muscle cells (58, 59). *ZIC1*, a zinc-finger protein that works alongside *PAX3* during early skeletal muscle development, was also upregulated at Day 0, suggesting the All-Risk haplotype may have a larger persistence of myogenic progenitor identity compared to the Non-Risk haplotype. This upregulation trend persisted to Day 6, as *PAX7* and *ZIC1* are still upregulated in the All-Risk haplotype compared to the Non-Risk haplotype (Fig. 5*I*). Confirmation of elevated *PAX3* and *PAX7* expression levels in the All-Risk haplotype was shown by measuring relative mRNA levels on Day 3 and Day 6 (Fig. S16*A-D*). We also observed several genes encoding transcription factors such as *ITM2a* (downregulated) and *DMRT2* (upregulated), which are downstream targets of *PAX3* in the myogenesis signaling cascade, as well as upregulation of *IGF2*, which activates pro-differentiation pathways via PI3K and AKT, suggesting the All-Risk haplotype may be entering later stages of myogenesis prior to the Non-Risk haplotype, despite no observed qualitative differences in skeletal muscle differentiation. Genes downregulated in All-Risk cells were found to be involved in energy metabolism and lipid metabolism, such as *PDP1, ABCA6, ATP10B, SSTR1*, and *CES1*, and in calcium homeostasis, such as *SYT14, OXTR*, and *AMPD3*. *PDP1* and *SYT14* were found to be expressed in the ANK1-Risk, but not the NKX6-3 Risk *SI Appendix* Dataset S3). The remaining genes were not differentially expressed in either the ANK1-Risk or the NKX6-3-Risk haplotypes (Fig. 5*I*, Fig. S17*A-B*). Thus, our results indicate that the All-Risk haplotype shows the most perturbations in gene expression of those important in myogenic structure and function, insulin signaling, and energy metabolism, suggesting that risk SNPs across the *NKX6-3/ANK1* locus have a combinatorial effect on T2D risk.

## Discussion

We have described a platform for experimentally dissecting GWAS loci using vSwAP-In, a simple DNA building method in yeast, and mSwAP-In, a Big DNA delivery method. We have exemplified this strategy by dissecting the *NKX6-3/ANK1* T2D GWAS locus in hiPSCs. vSwAP-In proved to be an efficient and scarless method for introducing variants to establish synthetic haplotype payloads of the *NKX6-3/ANK1* locus. The ability to build both natural and unnatural haplotypes is crucial for dissecting GWAS loci, as many SNPs are typically in high LD, complicating experimental validation of causal variants in patient-derived cells (5, 60). hiPSCs harboring the synthetic haplotypes can be expanded and differentiated to disease-relevant tissues. Here, we demonstrated the successful integration of several isogenic *NKX6-3/ANK1* haplotype variants: the All-Risk, the ANK1-Risk, the NKX6-3-Risk, and the Non-Risk haplotypes, using the mSwAP-In procedure, previously only reported on in mESCs (19, 20). Each haplotype variant contained different combinations of candidate risk SNPs in high LD with the lead SNPs identified in European and East Asian GWAS and eQTL data to better understand candidate risk SNPs in their genomic contexts. This technology should be extendable to other human loci to build cell line variants, similar to our findings in mESCs (19, 23, 61).

The landing pad hiPSC line AS0041^MC1^ represents a combined *NKX6-3/ANK1* gene cluster knockout system. This line demonstrated a drastic reduction in the ability to differentiate into both skeletal muscle cells and pancreatic progenitors, presumably due to the lack of *sANK1* and *NKX6-3,* respectively. Studies in mouse models have previously characterized functional effects of *sAnk1* knockouts (16, 17, 30), and did not find murine *Nkx6-3* to be expressed in pancreatic cell types (46), but instead it was reportedly expressed in gastric cell types (45–47). We were unable to detect *NKX6-3* expression, perhaps due to the extremely low abundance of the mRNA in islet samples (62). Our results suggest that *NKX6-3* might play a role in pancreatic cell differentiation, as implied by the significant downregulation in the AS0041^MC1^ hiPSCs of genes encoding pancreatic developmental transcription factors NKX6-1 and PDX1, which are required for islet and β-cell development (50, 63, 64). Further experiments are required to provide direct evidence for the expression of *NKX6-3,* or for its direct requirement for pancreatic differentiation. Specifically, the single deletions of the *NKX6-*3 and *sANK1* genes would shed light on this.

We also showed that reintegration of *NKX6-3/ANK1* haplotype variants harboring T2D risk SNPs restored differentiation capability into both skeletal muscle and pancreatic progenitor cells, proving the necessity of these genes in the differentiation processes. We then began characterizing the risk SNPs of the locus by quantifying *ANK1* gene expression differences in the haplotypes that were differentiated into skeletal muscle. We found that cells harboring the All-Risk and ANK1-Risk haplotypes had substantially higher levels of *ANK1* expression compared to the Non-Risk and NKX6-3-Risk haplotypes at Days 3 and Days 6 of differentiation, consistent with previous GWAS/eQTL studies (9, 12). The ANK1-Risk haplotype can be further dissected and differentiated to narrow down possible causal risk SNPs that maintain elevated *ANK1* expression levels. In addition to elevated *ANK1* levels, the All-Risk haplotype showed elevated expression levels of *PAX3* and *PAX7*, both early-stage myogenic transcription factors, suggesting that risk SNPs may directly affect skeletal muscle differentiation. In addition, the All-Risk haplotype was shown to have upregulated gene expression of early-stage myogenic factors, such as *PAX3* and *PAX7*. Further experimentation is required to uncover the roles *PAX3* and *PAX7* may play in the context of T2D risk. Perturbations to gene expression of genes important for skeletal muscle function, insulin signaling, and metabolism examined in the All-Risk haplotype, suggest that collectively, risk SNPs across the *NKX6-3/ANK1* locus may be playing a broader role in T2D risk. Overall, our results suggest that the *NKX6-3/ANK1* cluster plays an important role in pancreatic differentiation and skeletal muscle differentiation, and our work shows the first functional analysis of a series of GWAS haplotypes of the *NKX6-3/ANK1* cluster to help define the genomic basis of this T2D risk locus. In summary, haplotype DNAs produced synthetically and reintroduced into human iPSCs can faithfully recapitulate haplotype-specific patterns observed in patient-derived pancreatic islets and skeletal muscle tissue, providing insights into the behavior of uncommon natural haplotypes.

## Materials and Methods

Additional information is available in *SI Appendix*

### Base Construct Building

BAC RP11-111B9 (BACPAC) was digested using *in vitro* CRISPR, releasing a fragment corresponding to the genomic coordinates chr8:41,635,845-41,688,341 (hg38). To perform an *in vitro* CRISPR digest, crRNAs (*SI Appendix* Table S9) were diluted to 0.1 nM using Nuclease-Free Duplex Buffer (Integrated DNA Technologies (IDT): 11-05-01-12), and combined with 0.1 nM Alt-R® CRISPR-Cas9 tracrRNA. Oligos were mixed and diluted to 1 µM final concentration using Nuclease-Free Duplex Buffer and annealed at 98°C for 2 minutes, 95°C for 30 seconds, and decreasing 5°C/30 seconds to 20°C. The digest contained 25 nM of each crRNA-tracrRNA, 0.1 µM Cas9 Nuclease, S. pyogenes (New England Biolabs (NEB): M0386S), 1 *µ*g RP11-111B9, and 1X Buffer 3.1 (NEB: B7203S) in 40 µL. The reaction was incubated at room temperature for 10 minutes, and then at 37°C overnight. Digest was verified on a 1% low-melting agarose gel with a Lambda-PFG ladder (NEB: N0341S) and run on a BioRAD pulsed field gel apparatus. The digested BAC was co-transformed with linker DNAs (*SI Appendix* Table S3) into a linearized yeast assemblon vector (18) and yeast cells were selected on SC–Leu plates for two days at 30°C. The transformants were single-colony purified and screened with left and right junction assays (*SI Appendix* Table S4) using GoTaq Green (Promega: M7122) according to the manufacturer’s protocol. MC2 was cloned ∼1.6 kb away from the 3’ end of the digested BAC into the yeast strain base construct using the Ellis Lab Toolkit (25). Golden Gate Assembly was performed to create a sgRNA expression vectors using pWS082 and annealed oligos ((*SI Appendix* Table S9). The reaction was recovered into Top10 *E.coli* and digested with EcoRV (NEB: R3195S) to linearize the sgRNA expression cassette. The Cas9-sgRNA gap repair vector, pWS158, was digested with Esp3I (NEB: R0734S) and gel-purified. MC2 was amplified and gel-purified from a derivative of an MC2-based plasmid (19) using Q5 Master Mix (NEB: M0492L), and primers with homology to the base construct (*SI Appendix* Table S4). The base construct was grown to an OD = 0.5 and cells were made competent using lithium acetate and resuspended in a final volume of 30 µL per transformation. The following components were transformed to insert MC2: 50 ng digested gap-repair vector, 90 ng linearized sgRNA expression vector, and 400 ng of MC2 in a total of 30 µL. The transformation was plated onto SC–Leu–Ura plates, as the base construct backbone contains a *LEU2* marker, and Cas9-sgRNA gap repair vector contains a *URA3* marker. Yeast colonies were single-colony purified and screened via colony junction PCR using GoTaq Green (Promega: M7123).

### Building Haplotype Payloads

To build haplotype variants, either vSwAP-In or CRISPR-based editing such as the Ellis Lab Toolkit (25), or CREEPY (24) were performed. vSwAP-In was performed by PCR amplifying a *URA3* marker with 40 bp of homologies to the target region, and transformed into BY4741 yeast carrying the base construct. A second transformation step was performed by cotransforming a *URA3*-targeting Cas9-gRNA plasmid and segments containing SNP variants with overlapping homology. The Ellis Lab Toolkit was performed using similar methods to the previously described MC2 insertion, but with different annealed oligos and donor DNA. A full list of sgRNAs, oligos and donor DNA sequences can be found in Tables S2 and S3. Resulting haplotype yeast plasmids were recovered into TransforMax electrocompetent EPI300 E.coli (Biosearch Technologies: EC02T110) by miniprepping yeast plasmids according to manufacturer instructions (Zymo Research: D2004). 2 µL of the resulting miniprepped plasmid were transformed into EPI300 diluted with sterile water (1:4) and plated onto selective LB–Kan plates to select for recovered transformants. Transformants were then selected at random and miniprepped using the ZymoPure Plasmid Miniprep Kit (Zymo Research: D4210) for sequencing to validate recovered haplotype payload. Once sequencing confirms the recovery of the haplotype payload, plasmids were grown up using either the ZymoPure Maxiprep or Midiprep Kit following manufacturer’s instructions (Zymo Research: D4202, D4200).

### Cloning of Cas9-gRNA expression plasmids for hiPSC engineering

gRNAs targeting the *NKX6-3/ANK1* locus (for MC1 integration) were cloned into pX330 (Addgene: 48137) using BbsI-mediated Golden Gate Assembly (65). Oligos orNCH001 and orNCH002 were annealed to form the left-side gRNA-targeting upstream of endogenous *NKX6-3*. Oligos orNCH003 and oNCH004 were annealed to produce the right-side cutting downstream of *sANK1*. gRNAs targeting MC1 for payload deliveries were cloned into pX333 (Addgene: 64073), which contains two independent gRNA expression cassettes followed by a Cas9 cassette. Oligos orNCH009 and oNCH010 were annealed to form the gRNA targeting the left side of MC1 and outside the left homology arms of the synthetic payload, and orNCH011 and orNCH012 were annealed to target the right side of MC1 and outside the right homology arms of synthetic payload using Golden Gate Assembly in a two-step process using BbsI (NEB: R3539S) and BsaI (NEB: R3733S) (65). A full list of gRNAs used for integration are listed in *SI Appendix* Table S9.

### Delivering Haplotype Payloads into hiPSCs

#### Establishing the Landing Pad Cell Line

Cell line AS0041 is a hiPSC line derived from normal glucose-tolerant male-donor fibroblast cells. The fibroblast cells were previously reprogrammed using Stemgent StemRNA 3^rd^ Gen Reprogramming Kit (Reprocell: 00-0076) to create the AS0041 hiPSC line.

#### Routine culturing conditions

Prior to seeding, 6-well plates were coated with Geltrex (Gibco: A1413201) according to manufacturer’s instructions. Coated plates were incubated at 37°C for at least 1 hour prior to cell plating. AS0041 cells were seeded in 2 mL of StemFlex medium (Gibco: A3349401) + 1 µM Thiazovivin (STEMCELL Technologies: 7225) at a density of 3 x 10^5^ cells/well. Medium was changed every day, or double-fed (4 mL per well) for 2 days under a weekend-free protocol using StemFlex medium. Cells were routinely passaged once every 4-5 days, with a doubling time of approximately one day. For routine passaging, cells were treated with StemPro Accutase (Gibco: A1110501) and incubated at 37°C for 5-6 minutes to dissociate cells. The StemPro Accutase was neutralized using StemFlex medium + 1 µM Thiazovivin and cells were gently pipetted 2-3 times to maintain cell clusters. Cells were then spun down at 200*g* for 4 minutes and plated at a density of 3 x 10^5^ cells/well in a fresh, Geltrex-coated 6-well plate containing 2 mL of StemFlex medium + 1 µM Thiazovivin per well. The next day, medium was aspirated completely and replaced with 2 mL of StemFlex medium.

#### Nucleofection

AS0041 cells were pretreated with StemFlex + 10 µM of Thiazovivin 1 hour prior to nucleofection. Medium was completely aspirated, and cells were treated with 1 mL of StemPro Accutase and incubated at 37°C for 5-6 minutes to dissociate cells. The StemPro Accutase was neutralized using an equal volume of StemFlex medium + 10 µM Thiazovivin. Cells were gently resuspended to form a single-cell suspension and spun down at 200*g* for 4 minutes. Cells were then washed with DPBS -/- (no Ca^2+^/Mg^2+^) and counted using The Countess 3 (Invitrogen: A49862). For each transfection condition, 1-2 million cells were placed into individual 1.5 mL tubes and spun down again at 200*g* for 4 minutes. Pelleted cells were resuspended with 50 µL of the Lonza P3 Primary Cell Nucleofector Solution (Lonza: V4LP-3002). The DNA used for nucleofection is as follows: 7 µg of MC1 and 1 µg of each Cas9-gRNA cloned into pX330 for MC1 delivery, or 1 µg of the haplotype payload DNA, and 1 µg of dual gRNA-Cas9 cloned into pX333 for payload delivery. The DNAs were diluted in 50 µL of the nucleofector solution as well. The diluted DNA and cells were then thoroughly mixed, placed into the Lonza 4D nucleocuvettes, and transfected using program CB-150 on the Lonza 4D Nucleofector (AAF-1003X). Cells were plated onto 1 well of a 6-well plate coated with rhLaminin-521 (Gibco: A29248) containing 2 mL of StemFlex medium + 10 µM Thiazovivin. Puromycin (for MC1) or blasticidin (BSD) selection began 24 hours post-transfection at 0.6 µg/mL, or 8 µg/mL, respectively, for 6 days.

#### Lipofection

This protocol was adapted from Blanch-Asensio et al., *Nature Protocols* (2024) (66) and used for haplotype payload deliveries. AS0041^MC1^ cells were seeded in 12-well rhLaminin-521 coated plates containing 1 mL of StemFlex medium + 1 µM Thiazovivin at a density of 2 x 10^5^ cells per well for each transfection condition. When cells reached 60-80% confluency, they were washed with PBS. 1 mL of Opti-MEM medium was added to each well. The cells were lipofected using the Lipofectamine Stem Transfection Reagent (Invitrogen: STEM00003). DNA vectors (700 ng of haplotype payload DNA and 1 µg of dual gRNA-Cas9 were diluted in Opti-MEM for a total of 50 µL per transfection. Using a separate tube, 4 µL of Lipofectamine Stem Reagent was diluted in 46 µL of Opti-MEM for each reaction, and then mixed with the diluted DNA vectors. The mixture was incubated at room temperature for 10 minutes, and then added dropwise to the cells. The mixture was swirled and the cells were incubated at 37°C + 5% CO_2_ for 4 hours. 1 mL of warm Stemflex medium was then added to each well and the cells were returned to the incubator. The next day, a full medium exchange with warm StemFlex medium was performed to allow cells to recover. Two days post-lipofection, cells were cultured in StemFlex medium + 8 µg/mL blasticidin to begin selecting for payload integration. Selection was maintained for 5-6 days.

#### Single-cell Sorting and expansion of hiPSC clones

hiPSCs were allowed to recover for 1-1.5 weeks prior to cell sorting to ensure enough drug-resistant cells were present in the plate. Nucleofected or lipofected cells were pretreated with HaloTag Ligand JF549 or JF646 (Promega: GA1110, GA1120) up to 1 day prior to sorting to identify cells that contained MC1. For isolating cells harboring MC1, JF549/JF646+ cells were sorted. For isolating cells harboring correctly-integrated synthetic haplotype payloads, mNeonGreen+ and JF549/JF646-cells were sorted. hiPSCs were single-cell sorted into Geltrex-coated 96-well plates containing 100 µL of StemFlex medium + 1:20 dilution of CloneR2. Sorting was performed on the Sony SH800S Cell Sorter using a 100 µm sorting chip (Sony: LE-C3210). Four days after sorting, 100 µL of warm StemFlex medium was added to each well. Two days later, a full medium exchange was performed by aspirating the existing medium and replacing with 100 µL of fresh StemFlex medium. Cells were then selected again in StemFlex medium supplemented with 0.6 µg/mL puromycin for MC1 or 8 µg/mL blasticidin for haplotype payloads, and medium was changed every 2-4 days until large, established colonies were formed.

#### Verification of candidate hiPSC clones

Established colonies in the 96-well plates were dissociated using 0.5 mM EDTA in DPBS -/-, and gently pipetted to form cell clusters. 50% of the cell mixture was placed into a 96-well PCR plate for genotyping, and the remaining cells were placed into a Geltrex-coated 12-well plate containing 1 mL of StemFlex medium + 1 µM Thiazovivin for expansion. To extract crude gDNA, cells were resuspended in water supplemented with 0.3 µg/mL Proteinase K and lysed at 37°C for 1 hour, following heat inactivation at 99°C for 10 minutes. For MC1 verification, PCR screening was performed by amplifying the left and right junctions of MC1, the inner portion of MC1, and the genomic region of *ANK1* (*SI Appendix* Table S4). One candidate out of 93 clones contained bands amplifying both junctions and not *ANK1*. Several candidates contained only one junction, and not the other. Candidates that passed PCR-genotyping were expanded to 6-well plates, gDNA extracted using the QIAamp DNA Mini Kit (Qiagen: 51306) and sequence verified with Capture-seq for MC1 integration using the NextSeq 500. (67). Baits using the vector backbone of delivered payloads were used to capture sequences that would indicate backbone integration within the cells. For payload verification, a general PCR screening was performed by amplifying the left and right junctions of the payload, outside of the homology arms, the inner portion of MC1, and the genomic region of *ANK1*, where 25-50% of screened payloads contained both left and right junctions, no MC1, and contained *ANK1* (*SI Appendix* Table S4). Several clones containing both junctions were selected at random and expanded to 6-well plates, and gDNA extracted using the QIAamp DNA Mini Kit. PCR-verification was repeated for all junctions, *ANK1*, MC1, and vector backbone integration prior to be sent for Capture-seq (*SI Appendix* Table S4). Baits using the vector backbone and MC1 were used to capture sequences that would indicate backbone integration, and MC1 retention.

### Differentiation and Characterization of Skeletal Muscle Cells

#### Vector creation and Lentivirus production

Lentivirus vectors were created using an EF1α-MYOD1-Hygro plasmid (Addgene: Plasmid #120464). Lentivirus was produced by transfecting HEK-293T cells cultured in 10 10 cm plates using DMEM + 10% FBS (GemCell: 100-500-500) with the following ratios of DNA per plate: 3.5 µg lentivirus vector, 6 µg psPAX2 and 3 µg pMD2.G. The DNA mixture was diluted in 970 µL Opti-MEM and 30 µL Lipofectamine 2000. The mixture was incubated for 15-20 minutes and added drop-wise onto HEK-293T cells and gently swirled. Transfected cells were incubated for 2 days before recovering the virus. Two days post-transfection, medium containing the virus was removed and spun down at 200*g* for 5 minutes to settle dead cells. The supernatant was carefully removed and filtered through a 0.45 µm filter into a Millipore Amicon Ultra Centrifugal Filter (MilliporeSigma: UFC900396). Samples were then spun down at 4000*g* for 30 minutes at 4°C. Virus was quantified using Lenti-X GoStix Plus (Takara Bio: 631280)) which measures the p24 capsid protein levels (ng/mL). To quantify the multiplicity of infection (MOI), AS0041 cells were plated in a 6-well plate and infected with varying amounts of lentivirus. Survival was measured post-hygromycin selection using PrestoBlue. Samples with < 40% survival were used for calculating viral titer transduction units per mL of virus (TU/mL). The amount of lentivirus used to infect a 10 cm plate at an MOI = 10 was calculated by using the ratio of TU from the PrestoBlue assay to lentiviral concentration (ng) from the Lenti-X GoStix assay and the difference between the culture plate surface area (from a 6-well to a 10 cm) multiplied by 10. These calculations yielded ∼2 µg of lentivirus needed per 10 cm plate. Each batch of 10 x 10 cm plates used to make lentivirus yielded between 40-65 µg of lentivirus.

#### Myoblast differentiation

Myoblast differentiation protocol was adapted from Young et al., 2016) (29). Cells were plated in 10 cm plates coated with rhLaminin-521 (Gibco: A29248) at a density of ∼2 x 10^6^ cells per plate in SMC4 medium. Cells were then infected with MYOD1 lentivirus once they reached 60-80% confluency at an MOI=10. Polybrene (10 µg/mL) was used to enhance infection efficiency. Cells were spinoculated at 1250 rpm for 90 minutes at 32°C. The next day, cells were treated with hygromycin (50 µg/mL) to select for MYOD1 cassette integration. Two days later, cells were split onto 6-well Geltrex-coated plates containing basal medium (no bFGF) + 10 µM Thiazovivin at a density of ∼1×10^5^ cells/cm^2^. The next day, cells were induced in high-glucose DMEM + 15% FBS. Four days after induction, cells were differentiated in low glucose DMEM + 5% horse serum + 3 µg/mL doxycycline for 5-7 days until myotubes formed. Medium was changed daily. Roughly 2-3 million cells were collected on Day 0 (prior to infection), 72 hours (2 days post-selection) and 144 hours (2 days of high glucose induction). Cells from 0h and 72h were collected by dissociating cells with TrypLE and neutralizing with equal amounts of medium. 144h cells were collected by washing cells with DPBS and using a cell scraper to gently collect myoblast cells. Cells from all timepoints were spun at 300 *g* for 3 minutes and pelleted were resuspended in 50% FBS, 40% DMEM, and 10% DMSO and frozen at -80°C to maintain cell integrity. 10% of the cells at each timepoint were collected for qRT-PCR analysis.

Basal media: DMEM/F-12 (Gibco: 21041025), 20% Knock-out serum replacement (Gibco: 10828-010), 1% Non-essential amino acids (NEAA) (Gibco: 11140-050), 1% GlutaMAX (Gibco: 35050-061), 100µM beta-mercaptoethanol (Gibco: 21985023).

SMC4: Basal media, 5 µM Thiazovivin (STEMCELL Technologies: 72252), 0.4 µM PD0325901 (STEMCELL Technologies: 72184), 2 µM SB431542 (Selleck Chemicals: S1067), 1 µM CHIR99021 (Tocris: 4423), 10ng/mL human bFGF (Gibco: 13256-029).

#### Analyses of Differentiated Myoblast Cells

Myoblast cells were collected at 0h, 72h, and 144h post lentiviral infection. Differentiations of myoblast cells were collected at various early timepoints throughout the differentiation process for *sANK1* expression. Roughly 200,000-300,000 cells were collected for qRT-PCR analysis.

#### RNA Extraction

RNA from 0.5-1×10^6^ differentiated myoblasts at 0hr, 72h, and 144h post infection was extracted using TRIzol Reagent (Invitrogen 15596026) and Direct-zol RNA Miniprep Kit (Zymo Research: R2052) according to product instructions. RNA from 1-2×10^5^ differentiated pancreatic progenitor cells at Stages 1-4 was extracted using the Qiagen RNeasy Plus Kit (Qiagen: 74134).

#### qRT-PCR of Myoblast and Pancreatic Progenitor Cells

Extracted total RNA (∼500 ng) samples were reverse transcribed using SuperScript IV Reverse Transcriptase according to the manufacturer’s instructions (Invitrogen: 18090010). cDNA was diluted 1:10 with RNAse-free water and used as input for RT-qPCR using SYBR Green PCR Master Mix (Applied Biosystems: 4309155). The RT-qPCR reaction was set up by aliquoting 5 µL of 2X SYBR Green PCR Master Mix, and 4.5 µL of RNAse-free water into a 384-well plate. An Echo 550 liquid handler was used to dispense 0.5 µL of diluted cDNA and 50 nL of premixed qPCR primer pairs (50 µM) in each well. RT-qPCR reactions were performed on a LightCycler 1536 Real-Time PCR Thermal Cycler (Roche: 05334276001). A list of RT-qPCR primers can be found in *SI Appendix* Table S10.

#### RNA Quality Control and cDNA Synthesis

Total RNA quality and concentration were evaluated using the Agilent Bioanalyzer system with the RNA 6000 Pico Kit (Agilent Technologies: G2938-90035). Complementary DNA (cDNA) was generated from 1 ng of total RNA using the SMART-Seq mRNA HT Kit (Takara Bio: 634796). The synthesis process included 12 cycles of PCR amplification as per the manufacturer’s instructions.

#### Library Preparation and Quality Assessment

The resulting cDNA was quantified using the Quant-iT PicoGreen dsDNA Assay Kit (Thermo Fisher Scientific: P7589). Samples were subsequently diluted to a working concentration of 0.3 ng/µL. Sequencing libraries were prepared using the Nextera XT DNA Library Preparation Kit (Illumina: FC-131-1024). During library construction, 12 cycles of PCR were performed to amplify the fragments and incorporate unique dual barcodes. The integrity and fragment size distribution of the final libraries were verified using the Agilent 4200 TapeStation with High Sensitivity D1000 ScreenTape (Agilent Technologies: 5067-5584). Library concentrations were further validated with the Quant-iT system to facilitate precise normalization.

#### Pooling and Next-Generation Sequencing

Individual libraries were pooled in equimolar ratios, with calculations based on the molarity derived from TapeStation fragment analysis and Quant-iT concentration data. The final library pool was quantified by quantitative PCR (qPCR) using the KAPA Library Quantification Kit (Roche: 07960140001) with KAPA HiFi HotStart DNA Polymerase. Sequencing was performed on the Illumina NovaSeq X Plus system. Data were generated using a paired-end 50-cycle sequencing strategy.

#### RNA-Seq analysis on Myoblast samples

Fastq files were downloaded and read quality was visualized. Reads were then aligned to human genome and gene counts were generated using STAR (v2.7.7a). Gene count files were then used for DESeq2 analysis, and data was normalized to the Non-Risk samples. Normalized data from DESeq2 were filtered using p-adjusted values < 0.05 and the absolute value of the log2 fold change > 1. GO enrichment analysis (68, 69) was then performed and plotted. Volcano plots were made by subsetting the normalized data from DESeq2 into “All-Risk”, “ANK1-Risk”, and “NKX6-3-Risk” and filtering for upregulated genes (log2 fold change > 1) and downregulated genes (log2 fold change < -1) and by p-adjusted values < 0.05. Plots were then made using ggplot (70), and data from Day 0, Day 3, and Day 6 of each haplotype were plotted individually. Gene trajectories were made by performing DESeq2 on gene count files, and then taking the variance stabilizing transformation (vst) of the DESeq2 results for genes expressed on Day 0, Day 3, and Day 6. Line graphs were then plotted using ggplot (70).

#### Immunostaining of Skeletal Muscle Cells

Cells were fixed using Formalin (4% final concentration) and incubated for 20 minutes at room temperature. Cells were then permeabilized using 0.3% Triton-X 100 for 10 minutes at room temperature. 10% Normal Goat Serum (Thermo Scientific: 50062Z) was used to block cells for 1 hour. After blocking, cells were treated with 2 µg/mL MF20 (Invitrogen: 50-6503-82) in 1% Normal Goat Serum and 0.1% Triton-X 100 and incubated overnight at 4°C with rocking. The following day, cells were washed with PBS twice to remove residual antibody. Cells were then stained with DAPI (1:1000) and incubated at room temperature for 15 minutes. Cells were then washed three times with PBS and imaged.

### Adaptation of iPSC lines for Pancreatic Progenitor Differentiation

Engineered hiPSCs delivered to NYSCF were not immediately usable on NYSCF’s automated cell culture platforms. To adapt the hiPSCs to NYSCF’s culture conditions and to generate sufficient inventory of each cell line, cells were initially thawed, expanded, and frozen in 2D barcoded Matrix tubes (Thermo Scientific: 3741) as previously reported (71), though here we used StemFlex medium (Gibco: A3349401) rather than mTeSR1.

#### iPSC expansion

All liquid handling steps were performed using NYSCF’s custom Hamilton STAR systems except for thawing, which was performed manually.

Matrix tubes containing frozen hiPSCs were removed from liquid nitrogen storage and transferred on dry ice prior to thawing. Vials were placed into a 37°C water bath for ∼90 seconds before resuspension in StemFlex medium (Gibco: A3349401) containing 10% CloneR (STEMCELL Technologies: 05888) to neutralize the cryopreservative. The cells were transferred into 15 mL conical tubes, to which additional StemFlex + 10% CloneR was added to ensure complete neutralization of the dissociation reagent (total volume per conical tube: 15 mL). A 10 µL aliquot of each cell suspension was taken for cell counting using a dead/total count application on an Opera Phenix High-Content Screening System (herein referred to as Opera Phenix; Revvity: HH14001000). The cell suspensions were centrifuged at 800 rpm for 4 minutes. The supernatant was aspirated without disturbing the cell pellet, and cells were resuspended in StemFlex medium + 10% CloneR and plated on 12-well Cultrex-coated plates (Corning: 3513) at a density of 3 x 10^5^ cells per well. Each well was topped up with additional StemFlex + 10% CloneR to a total volume of 1 mL as needed. After thawing and plating, the hiPSCs were cultured for 48 hours at 37°C in 5% CO_2_ and StemFlex + 10% CloneR without changing medium. From 48 hours onward, cells were fed daily with StemFlex + 1X penicillin/streptomycin (pen/strep; Gibco: 15070-063).

Upon reaching 60-80% confluence, the cells were passaged into three identical Cultrex-coated 12-well plates. Briefly, culture medium was aspirated without disturbing the cell layer, and wells were washed once with 500 µL of Accutase (Sigma-Aldrich: A6964). After washing, 500 µL of fresh Accutase was added to each well, and cells were incubated at 37°C in 5% CO_2_. Once fully dissociated, 500 µL of StemFlex + 10 µM Y-27632 (Selleck Chemicals: S1049) + 1X pen/strep was added to each well to neutralize the Accutase (Accutase-neutralization ratio: 1:1). The cell suspensions were transferred into an intermediate 96-well round-bottom plate (Corning: 3958) and centrifuged at 120*g* for 5 minutes. Following centrifugation, the supernatant was aspirated, and the cells were resuspended in 1 mL of StemFlex + Y-27632 + pen/strep. A 10 µL aliquot of the cell suspension was taken for cell counting using a dead/total count application on the Opera Phenix. After counting, the cells were seeded into the destination 12-well plates at a specified target density (2.5-3.0 x 10^5^ cells per well), with the volume transferred dependent on the cell count. All wells were topped up with additional StemFlex + Y-27632 + pen/strep to a total volume of 1 mL. The destination plates were returned to the incubator for a minimum of 24 hours. After 24 hours, a half-feed was performed with StemFlex + pen/strep, followed by daily media changes. Upon reaching 60-80% confluence, the cells were again passaged into identical Cultrex-coated 12-well plates.

### Automated generation of iPSC-derived pancreatic progenitor cells

Human iPSC-derived pancreatic progenitor cells were generated following an in-house 7-stage differentiation protocol for hiPSC-derived pancreatic islets, which was carried out until the end of Stage 4. The differentiation started on Day 0 with a normalized passage of hiPSCs into Cultrex-coated 12-well plates at a cell density of 8.5 x 10^5^ cells per well, as described above. After the destination plates were seeded, the cells were incubated with StemFlex + Y-27632 + pen/strep overnight at 37°C and 5% CO_2_ to ensure the formation of a 100% confluent iPSC monolayer.

Subsequently, the cells were cultured with Stage 1 medium for 4 days (Day 1 to Day 4), with daily medium exchange, at 37°C and 5% CO_2_ for definitive endoderm differentiation. Then, the cells were cultured with Stage 2 medium for 2 days (Day 5 to Day 6), with daily medium exchange, at 37 °C and 5% CO_2_ for primitive gut differentiation. Afterwards, the cells were cultured with Stage 3 medium for 3 days (Day 7 to Day 9) with daily medium exchange, at 37 °C and 5% CO_2_ for posterior foregut differentiation, followed by culturing with Stage 4 medium for 4 days (Day 10 to Day 13), with daily medium exchange, at 37°C and 5% CO_2_ for pancreatic progenitor differentiation. All medium exchanges were performed using NYSCF’s custom Hamilton STAR systems, with minimal human intervention other than transporting plates to and from the incubator and preparing medium. All cell differentiation media were freshly supplemented with their respective growth factors on the day of medium exchange. On the first day of each distinct stage in the differentiation, cells were washed once with the new stage basal medium before adding the final, supplemented differentiation medium.

Media compositions for each stage are as follows, Stage 1 (definitive endoderm) media: commercially available definitive endoderm differentiation kit (STEMCELL Technologies: 05110), specifically, on Stage 1 Day 1, 100X Supplement CJ and 100X Supplement MR from the kit were added into the 1X Definitive Endoderm Basal Medium at a dilution of 1:100 to prepare the final media. On Stage 1 Day 2 through Day 4, only 100X Supplement CJ was added to the 1X Definitive Endoderm Basal Medium at a dilution of 1:100 to prepare the final media. Stage 2 (primitive gut) media: Stage 2 basal [RPMI 1640 medium, GlutamMAX™ supplement (Gibco: 61870127), 0.5X B27 (B27™ Supplement 50X, Serum-Free, Fisher Scientific: 17504-044), 1X pen/strep], supplemented with 50 ng/mL KGF/FGF-7 (R&D Systems 251-KG). Stage 3 (posterior foregut) media: Stage 3 basal [DMEM, High Glucose, GlutaMAX™ supplement (Thermo Fisher Scientific: 10566-024), 0.5X B27, 1X pen/strep], supplemented with 0.25 µM KAAD-cyclopamine (Reprocell: 04-0028), 2.00 µM retinoic acid (Reprocell: 04-0021), 0.25 µM LDN-193189 (Reprocell: 04-0074). Stage 4 (pancreatic progenitor) media: Stage 4 basal [DMEM, High Glucose, GlutaMAX™ supplement, 0.5X B27, 1X pen/strep], supplemented with 50 ng/mL recombinant hEGF (R&D Systems: 236-EG), 10 mM nicotinamide (Sigma-Aldrich: N0636), 50 ng/mL KGF/FGF-7. All media were sterile filtered with a 0.22 µm filter (EMD Millipore: SCGP00525; Nalgene: 569-0020) prior to cell culture.

### Preparation of samples for flow cytometry, RT-qPCR, and sequencing analyses

At the end of each stage of the differentiation, samples were collected for flow cytometry, RT-qPCR, and sequencing analysis as follows. Culture medium was aspirated without disturbing the cell layer, and wells were washed once with 500 µL of dissociation reagent (Stages 0, 1: Accutase; Stages 2-4: TrypLE Select Enzyme 1X [Gibco: 12563011]). After washing, 500 µL of the respective dissociation reagent was added, and cells were incubated at 37°C in 5% CO_2_.

Once fully dissociated, the cell suspensions were transferred into 15 mL conical tubes, and the dissociation reagent was neutralized by adding 4.5 mL of DMEM/F-12, GlutaMAX™ supplement (Gibco: 10565-018) + 10% fetal bovine serum (FBS; Gibco: 10082-147), for a total volume of 5 mL. 10 µL of each neutralized cell suspension was taken for cell counting using a dead/total count application on the Opera Phenix. From each neutralized cell suspension, 500 µL was transferred to a TruTaper 96-well block (Analytical Sales & Services: 968810GC) for flow cytometry analysis as described in the following section, and 250 µL was transferred to a Matrix tube. These Matrix tubes were centrifuged at 600*g* for 5 minutes, after which the supernatant was aspirated to leave dry cell pellets for downstream RT-qPCR analysis. The Matrix tubes were then stored at -80°C.

The remaining cell suspensions in the conical tubes were delegated for sequencing analyses. These conical tubes were centrifuged at 800 rpm for 4 minutes, after which the supernatant was aspirated and cells were resuspended in 700 µL of CELLBANKER 1 freezing medium (Amsbio: 11910). The cell suspensions were immediately transferred into Matrix tubes, which were then placed into a CoolBox 96F System (Biocision: BCS-147) and stored at -80°C. After 24 hours, the Matrix tubes were transferred from -80°C to liquid nitrogen storage.

### Flow Cytometry Analysis

All reagent applications and removals were performed using the NYSCF robotic systems.

### Live staining, fixation and permeabilization

Following the sample preparation process described above, the TrueTaper block containing live cell flow cytometry samples was centrifuged at 130*g* for 5 minutes. After aspirating the supernatant, samples were resuspended in 1mL per sample 1X DPBS (Gibco: 14190-144) containing fixable live/dead dye (BD Biosciences, green: L34970A, violet: L34964A) at dilution 1:2000. Cell suspensions were incubated in dye suspension in the dark at room temperature for 30 minutes, then washed by 4x dilution with 1X DPBS before centrifugation, supernatant removal and resuspension in fixative. Cells were fixed for 12 minutes in the dark at room temperature in 1mL 4% paraformaldehyde (PFA; Electron Microscopy Solutions: 15710-S)/1X DPBS, washed by 5x dilution with 1X DPBS before centrifugation (fixed cell suspension 300*g*), supernatant removal, and resuspension in *Flow buffer*. Samples were permeabilised by treating with 500 μL of *Flow buffer* (1X DPBS, 0.1% Triton-X-100 [Sigma-Aldrich: 93443], 5% normal donkey serum [Jackson ImmunoResearch Laboratories: 017-000-121]), incubated for 45 minutes in the dark at room temperature. A pool of 10 µL of each sample was retained as an unstained control for FACS. Samples were stained for analysis of the specific differentiation stage as described below.

### Sample staining

For each stage of the differentiation, antibody cocktails were prepared and applied. Each sample was prepared with about 2.5 x 10^5^ fixed cells in a 100 μL working volume of Flow buffer. Samples were incubated with primary antibodies in the dark at 4°C overnight. Samples were washed by 10x dilution with Flow buffer before resuspension in secondary antibody for 2 hours. After washing, samples were resuspended in a 300 µL 1X DPBS for flow cytometry event collection.

*Stage 0*

Primary antibodies used were Oct4-BV421 (BD Biosciences: 565644) at a 1:50 dilution, and Rabbit Nanog mAb (Cell Signaling Technologies: #D73G4) at a 1:100 dilution, and then incubated with a secondary antibody donkey-anti rabbit Alexa Fluor®647 (Invitrogen: A-41573) at a 1:2000 dilution for 2 hours.

*Stage 1*

Antibodies used were Oct4-BV421 (BD Biosciences :565644) at a 1:50 dilution, and Sox17-AF647 (BD Biosciences: 562594) at a 1:100 dilution for 2 hours.

*Stages 2, 3, and 4*

Antibodies used were PE conjugated anti h/mPDX1 (R&D Systems: IC2419P) at 20 µL/1M cells, AlexaFluor 647 conjugated anti human NKX6.1 (BD Pharmingen: 563338) at a 1:200 dilution, and AlexaFluor 488 conjugated anti human HNF4-alpha (Abcam: AB216897) at a 1:100 dilution.

FACS event collection and analysis were carried out on an Attune NxT (Thermo Fisher Scientific) using onboard software.

## Author Contributions

N.C., A.V., W.Z., J.M.L., F.C., S.J.P., and J.D.B. designed research; N.C., A.V., W.Z., and S.U. performed research; A.V., C.M., H.A., M.T.M., and S.J.P. performed sequencing; N.C., A.V., W.Z., R.B., S.J.P., and J.D.B. analyzed data; X.M., N.D., K.R., D.P., J.G., and F.C. performed pancreatic differentiations; A.V., W.Z., S.U., R.B., D.F., S.J.P., and J.D.B. contributed insight and knowledge, N.C., R.B., and J.D.B. wrote the paper; all authors reviewed the final manuscript.

## Acknowledgments

We thank members of our laboratories for their support and suggestions to this study. This study was supported by the National Human Genome Research Institute’s Centers of Excellence in Genomic Science (NHGRI–CEGS) Grant RM1HG009491 to J.D.B., NIH Institutional Training Grant T32GM136542 to N.C, NIH grants UM1DK126185 and R01DK117960 to S.J.P. We thank Mike Erdos and colleagues at NHGRI for providing the iPSC line.

