## Supplemental figures and tables for "Genome writing and Targeted Delivery of the *NKX6-3/ANK1* gene cluster and its Type 2 Diabetes GWAS Variants to Human iPSCs": SI Appendix containing Figures S1-S17, and Tables S1-S11_V5for BioRXiv.pdf

Noor Chalhoub et al.

### Figures

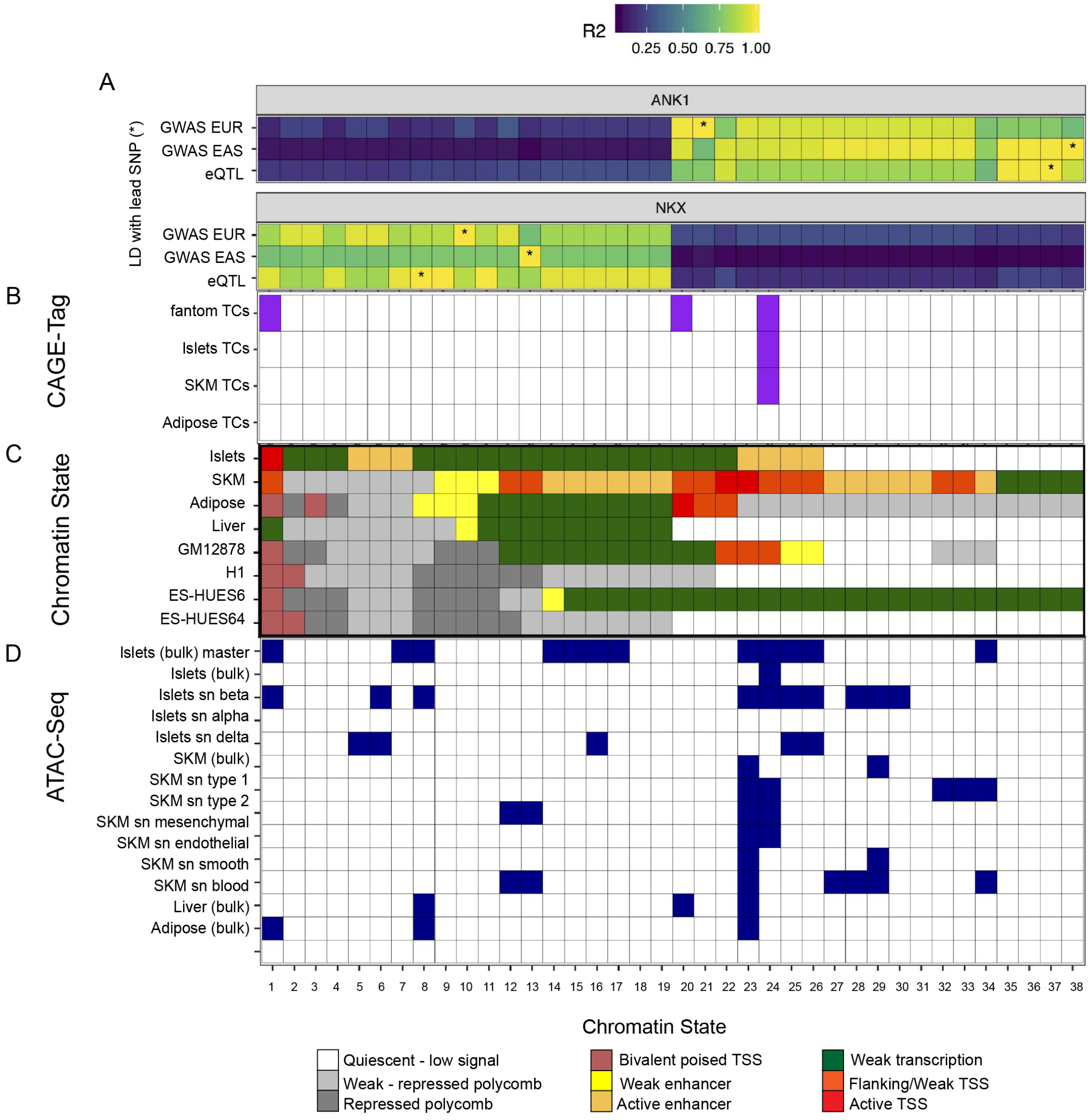

**Fig. S1. A.** Functional annotation overlaps with eQTL SNPs nominate regulatory variants at the ANK1-NKX6-3 locus. From top: LD $r^2$ (R2) with the lead SNP (marked with *) for European T2D GWAS, East Asian T2D GWAS, and the colocalized ANK1 eQTL in skeletal muscle and NKX6-3 eQTL in pancreatic islets. **B.** SNP overlap with CAGE tag clusters from the FANTOM study, or those identified in islets, skeletal muscle and adipose tissues is shown in purple. **C.** SNP overlap with chromatin (ChromHMM) states in relevant tissues and cell types. **D.** SNP overlap with bulk (orange label) and single nucleus (blue label) ATAC-seq peaks for in relevant tissues and cell types.

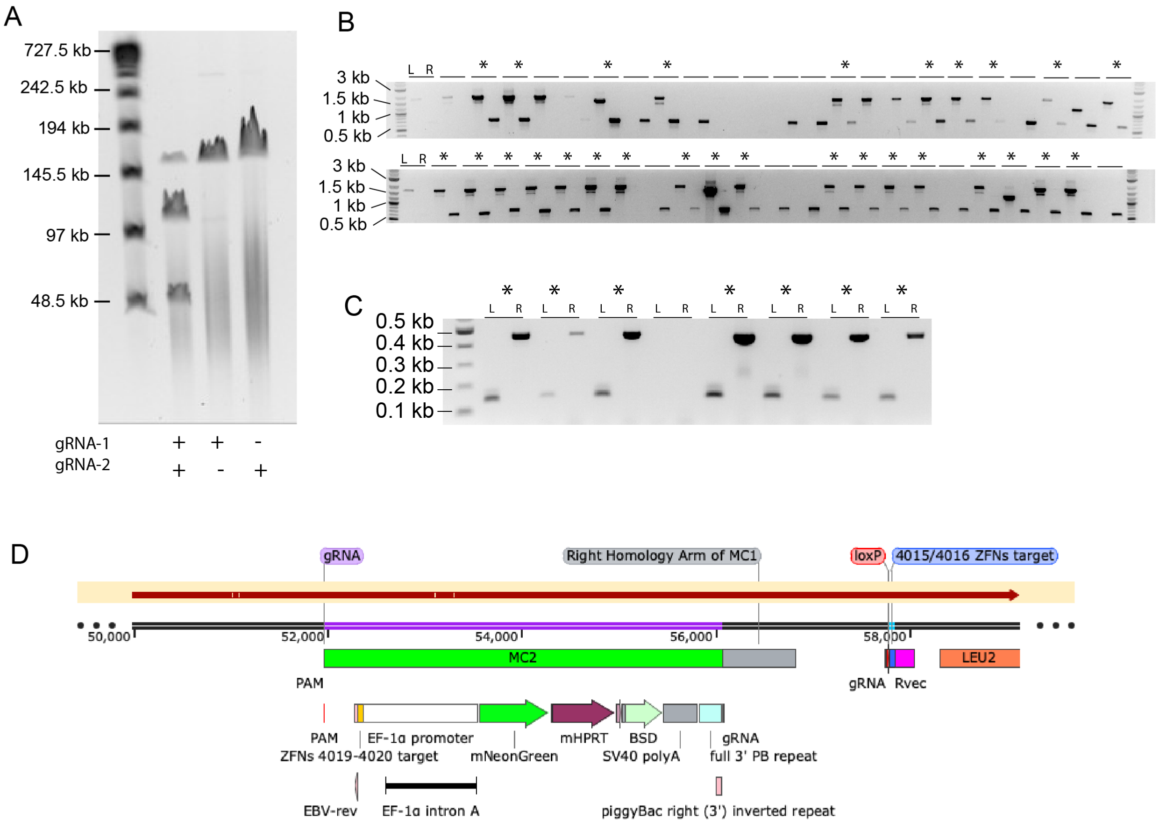

**Fig. S2. A.** Pulsed-field gel electrophoresis of bacterial artificial chromosome (BAC) RP11-111B9 digested *in vitro* using 2 CRISPR-Cas9 RNPs, resulting in a 52.8-kb segment used for base construct assembly. **B.** Left (L) and right (R) junction PCR screening of single-colony purified assembled base construct yeast transformants. 26 out of 48 colonies contained both left and right junctions (*). **C**. Left (L) and right (R) junction PCR screening of MC2 insertion from single-colony purified yeast containing the base construct payload. Seven of eight colonies contained both junctions (*). **D.** Snapshot of SnapGene file showing alignment of Base Construct plasmid sequenced by Nanopore following MC2 cloning. Alignment shows that MC2 has been successfully inserted.

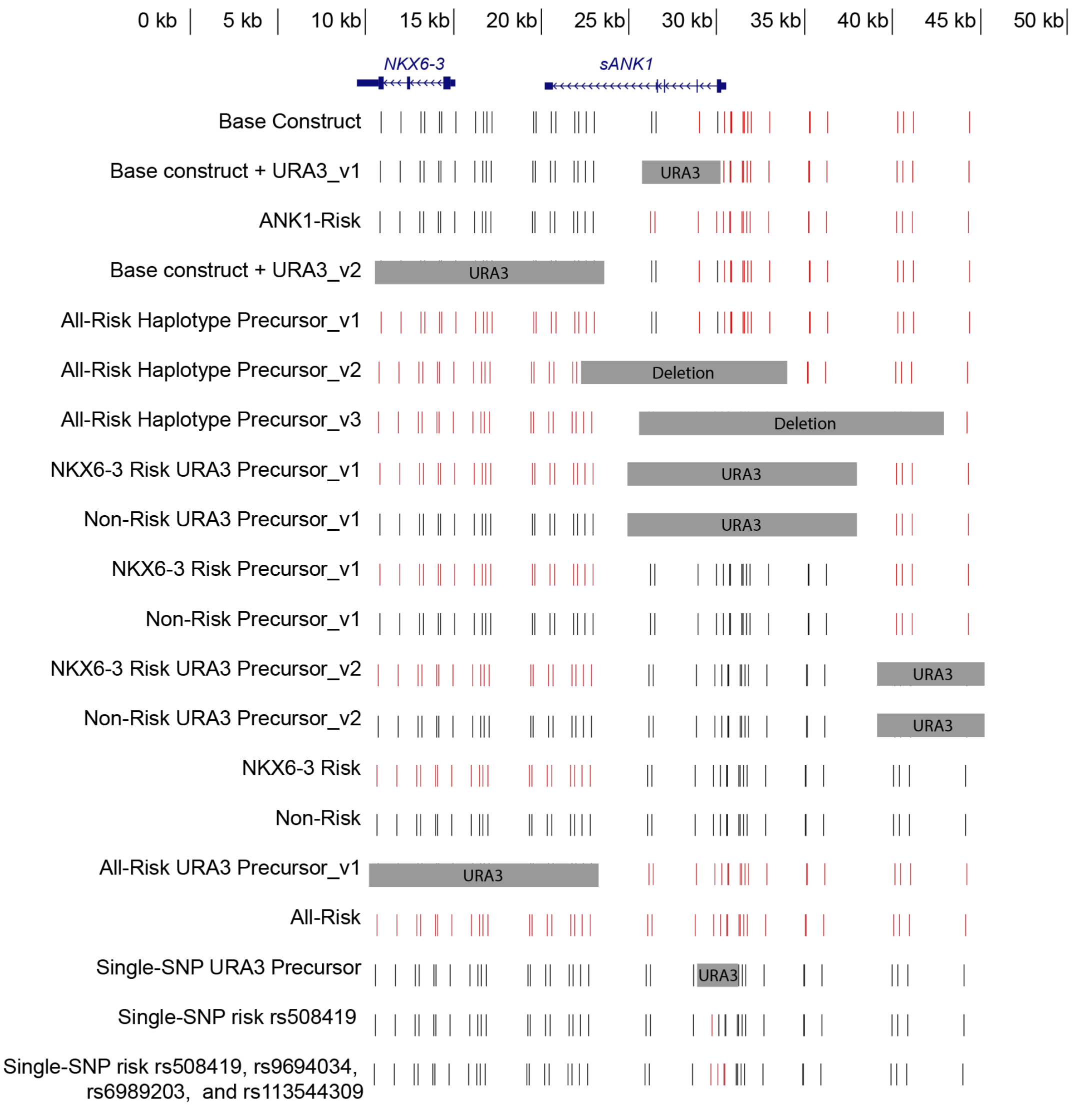

**Fig S3**. Sequential building of the haplotype variants in which first a Base-Construct was created, and *URA3* was transformed to create precursors which later became the ANK1-Risk and NKX6-3 Risk haplotypes. The All-Risk haplotype was created by transforming a URA3 marker replacing the NKX6-3 SNPs into the ANK1-Risk haplotype, and swapping it with segments PCR-amplified from the NKX6-3-Risk haplotype. Additional targeted risk-SNP haplotypes were made by performing vSwAP-In using the Non-Risk haplotype, and introducing a gene fragment that is risk for rs508419 or rs508419, rs9694034, rs6989203, and rs11354430; however, these constructs were not used for downstream experiments.

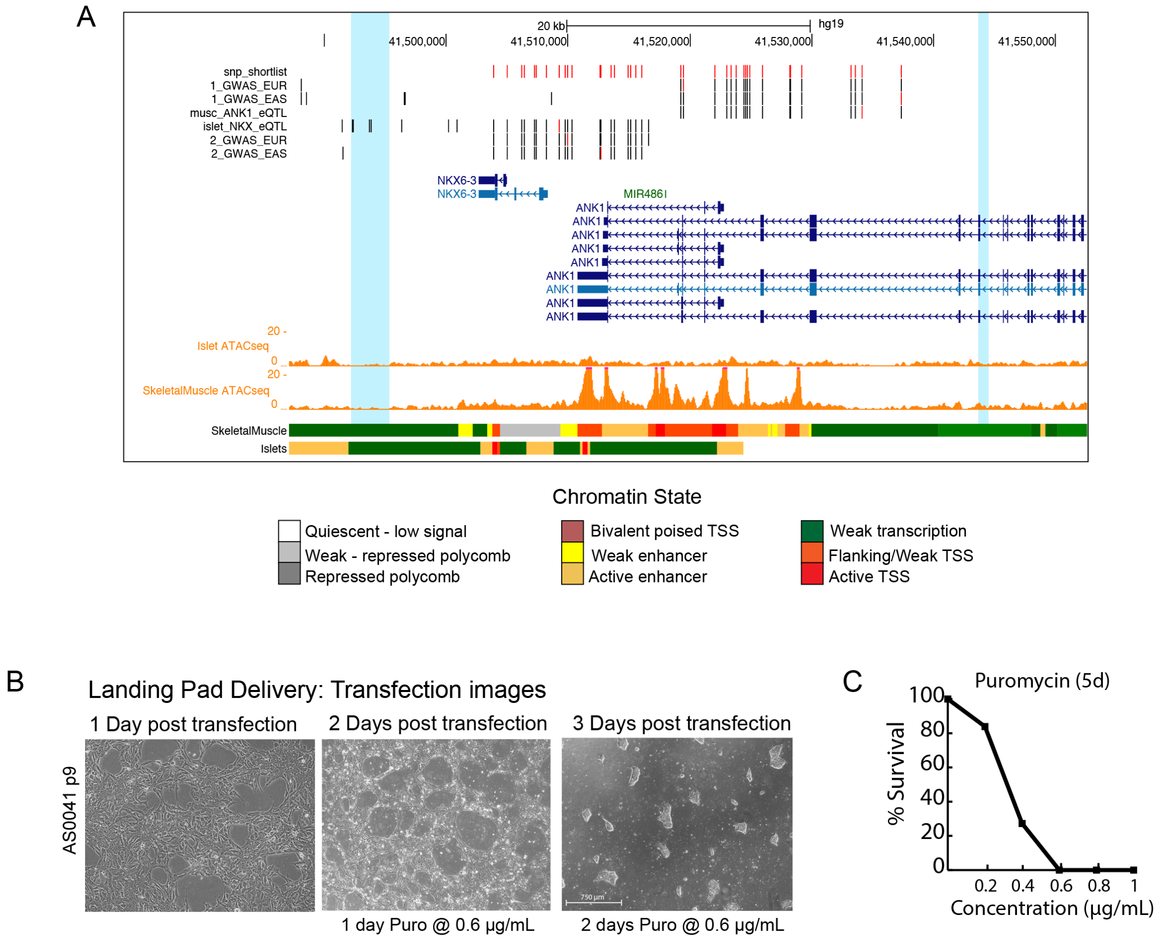

**Fig. S4. A**. Genome browser shot of the *NKX6-3/ANK1* locus with tracks of ATAC-seq peaks and chromatin states of islets and skeletal muscle tissue that correspond to those identified in Fig. S1C. Blue highlights show low activity regions with weak transcription, which were selected as regions for Cas9-gRNA cut sites for locus deletion and payload integration, as well as MC2 insertion. **B.** Images of AS0041 after MC1 transfection and selection. **C.** Puromycin kill curve to determine the optimal amount of selection to use on cells. Puromycin selection began 24 hours post MC1 transfection. For subsequent experiments, we used 0.6 µg/mL puromycin in selective medium.

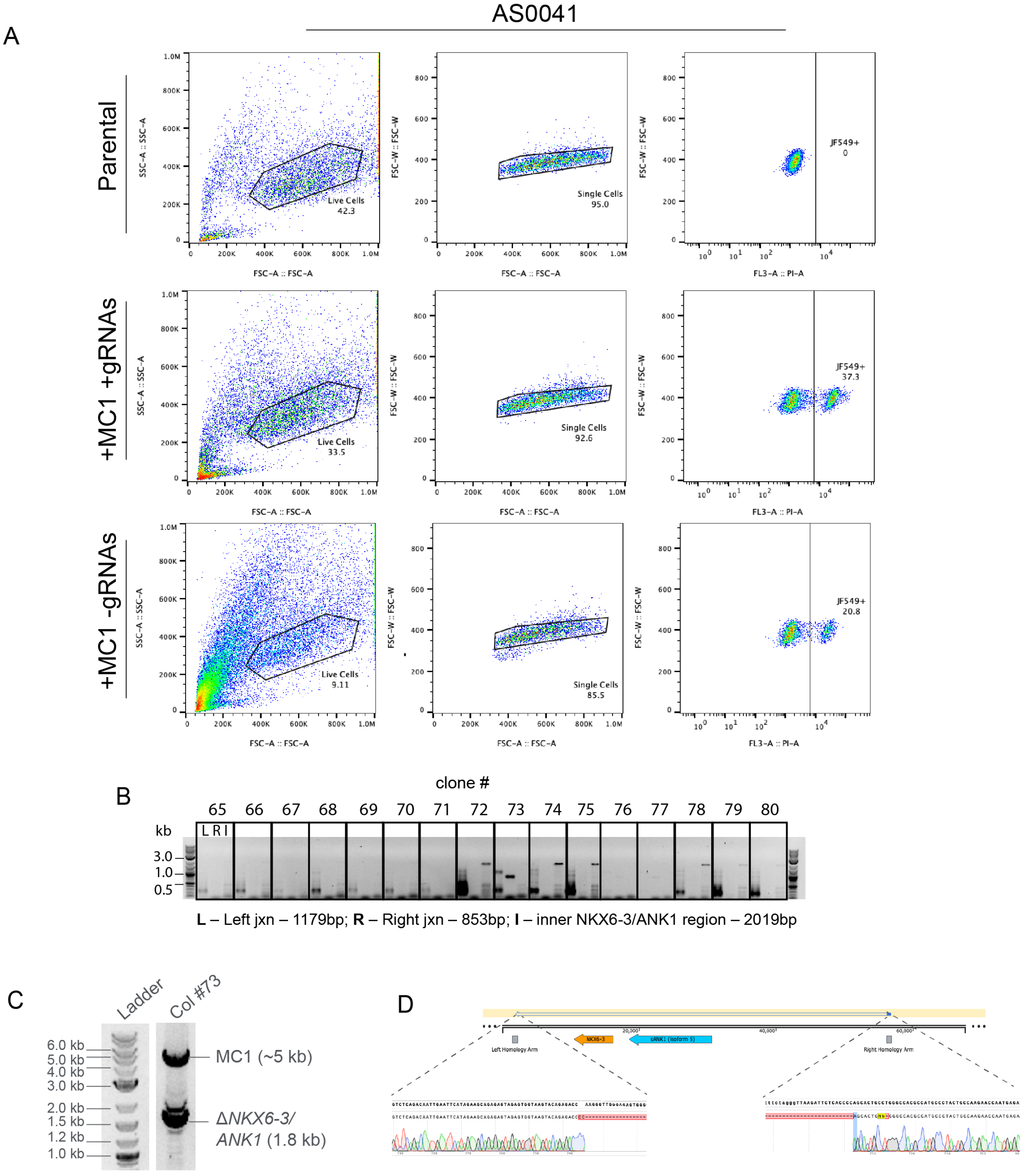

**Fig S5. A.** Single-cell sorting of AS0041 cells transfected with MC1 + gRNAs for JF549+ cells. 37.3% of cells in the population were shown to be JF549+. **B.** Junction PCR on expanded cells following single-cell sorting. One clone (#73) was found to have bands of both left and right junctions of MC1, and no band for *NKX6-3/ANK1*. **C.** PCR across MC1 on clone #73, showing two bands (1.8 kb for *NKX6-3/ANK1* deletion, and 5 kb for MC1), indicating hemizygous insertion of MC1 and a deletion removing both *NKX6-3* and *sANK1*. **D.** Sanger sequencing results of 1.8 kb band aligned to *NKX6-3/ANK1* flanking regions as indicated. Deletion sites align to gRNA cut sites, suggesting NHEJ following gRNA/Cas9 cutting.

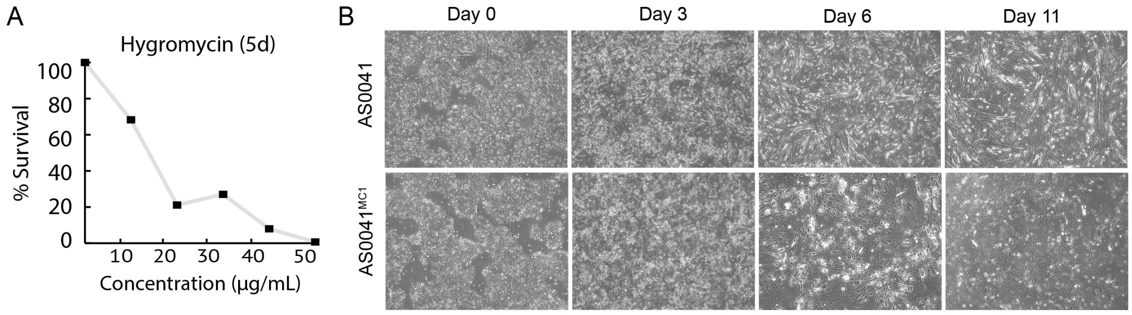

**Fig S9. A.** Hygromycin kill curve on AS0041 parental cells to determine optimal concentration of Hygromycin following *MYOD1*-lentiviral infection to select for cells that have taken up the lentivirus. The determined concentration utilized for subsequent experiments was 50 µg/mL **B.** Images of parental cell line AS0041 and MC1 cell line AS0041$^{MC1}$ at Days 0, 3, 6, and 11 post *MYOD1*-lentivirus infection. AS0041$^{MC1}$ did not properly differentiate as seen by the lack of myotubes compared to AS0041.

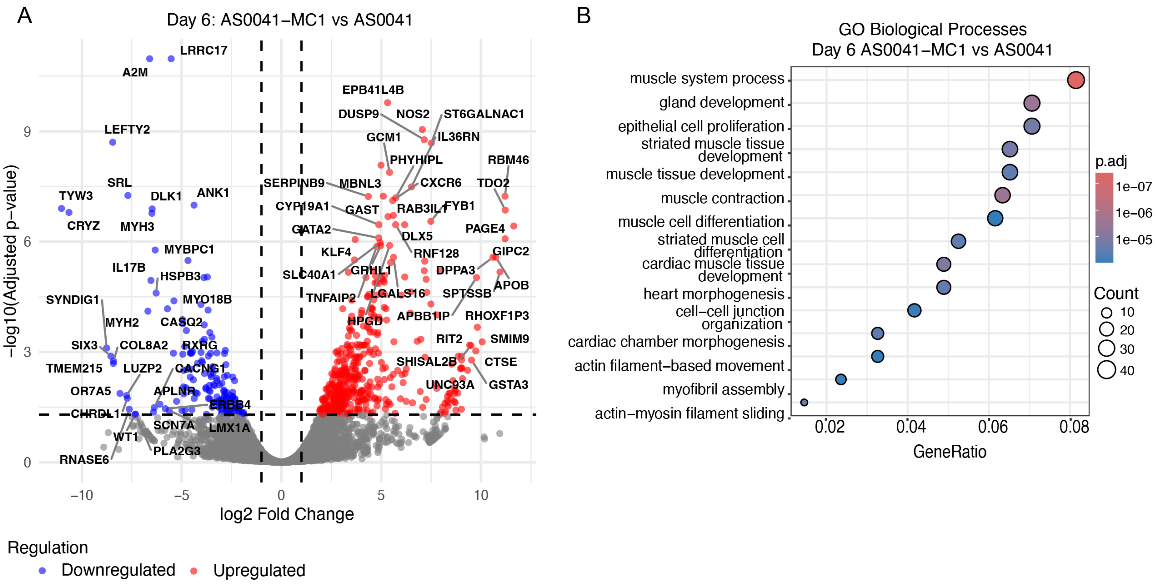

**Fig. S7. A.** Volcano plot of differentially expressed genes 6 days post-differentiation of AS0041$^{MC1}$ compared to parental cell line. Genes that play important roles in late-stage skeletal muscle development were shown to be downregulated. **B.** GO analysis for biological processes for AS0041$^{MC1}$ cells compared to AS0041 parental cells 6 days post-differentiation. Processes shown are related to muscle development and function

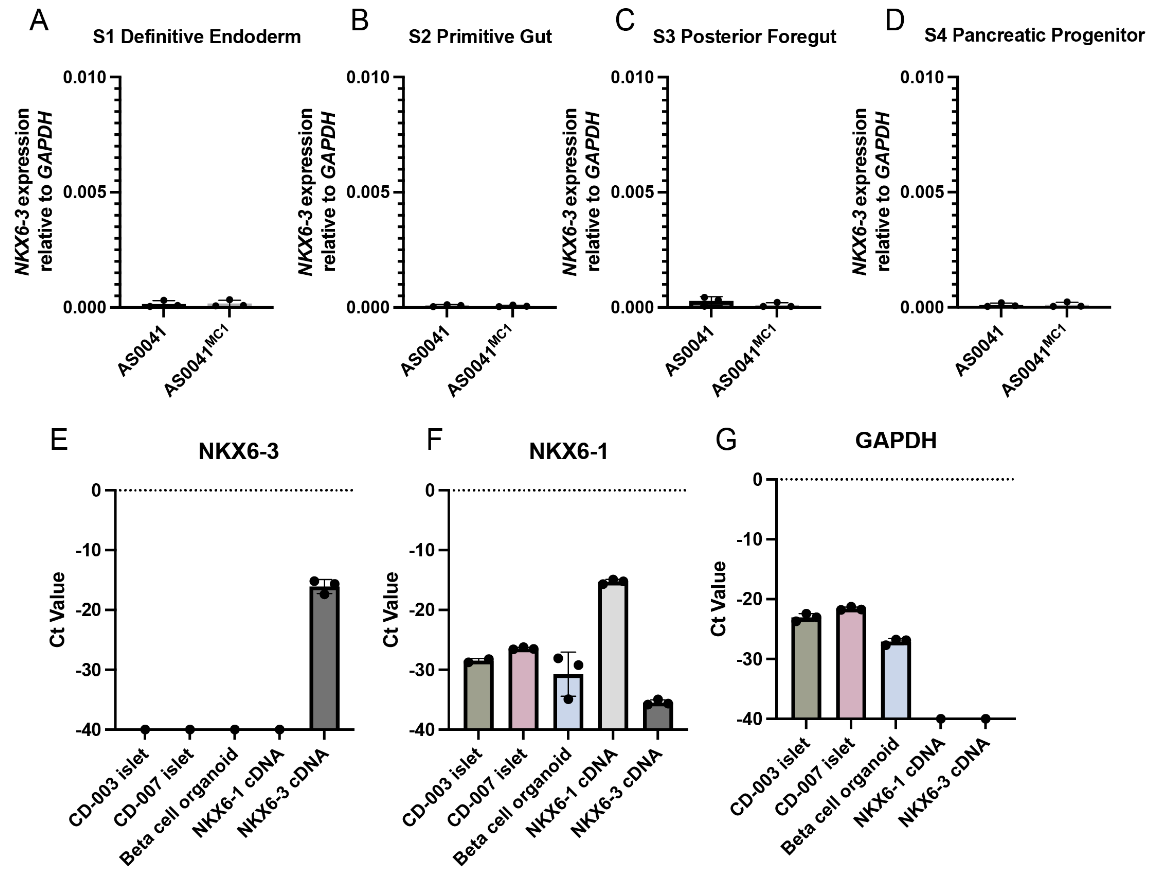

**Fig S8. A-D.** qPCR data quantifying *NKX6.3* mRNA from AS0041 and AS0041$^{MC1}$ at various stages of pancreatic differentiation. No *NKX6.3* transcription was detected. **E.** Raw Ct values quantifying *NKX6-3* mRNA plotted at a -Ct scale to visualize expression. NKX6-3 primers are functional al NKX6-3 cDNA was properly amplified. **F**. Raw Ct values quantifying *NKX6-1* cDNA plotted at a -Ct scale to visualize expression. NKX6-1 primers properly amplified *NKX6-1* cDNA suggesting primer specificity. **G.** Raw Ct values quantifying *GAPDH* mRNA plotted at a -Ct scale to visualize expression GAPDH primers amplified islet and organoid samples, but not the NKX6-1 and NKX6-3 cDNA samples.

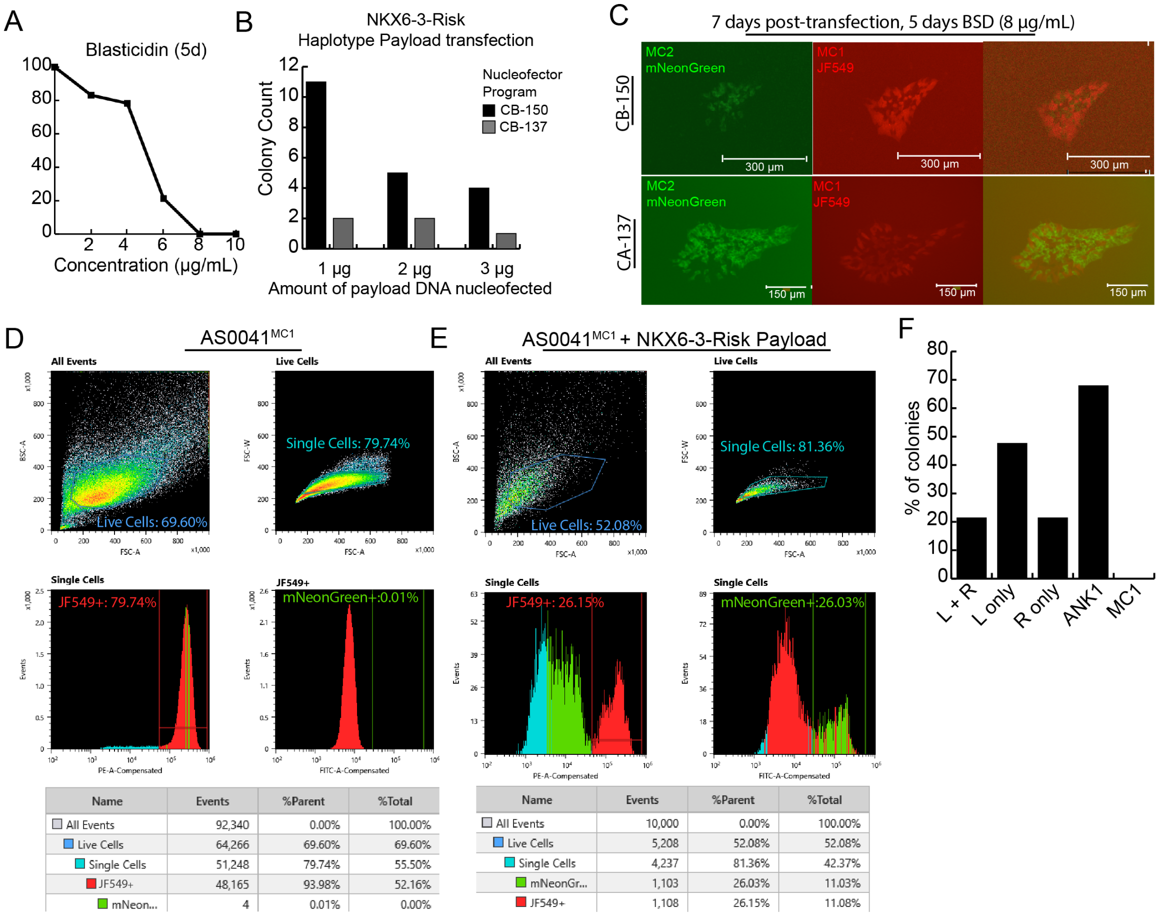

| Name | Events | %Parent | %Total |
|---|---|---|---|
| All Events | 92,340 | 0.00% | 100.00% |
| Live Cells | 64,266 | 69.60% | 69.60% |
| Single Cells | 51,248 | 79.74% | 55.50% |
| JF549+ | 48,165 | 93.98% | 52.16% |
| mNeon... | 4 | 0.01% | 0.00% |

| Name | Events | %Parent | %Total |
|---|---|---|---|
| All Events | 10,000 | 0.00% | 100.00% |
| Live Cells | 5,208 | 52.08% | 52.08% |
| Single Cells | 4,237 | 81.36% | 42.37% |
| mNeonGr... | 1,103 | 26.03% | 11.03% |
| JF549+ | 1,108 | 26.15% | 11.08% |

**Fig S9. A**. Kill curve to determine optimal concentration of BSD for payload selection, with 0.8 μg/mL showing total cell death after 5 days of selection. **B.** Colony counts of surviving AS0041$^{MC1}$ nucleofected with NKX6-3-Risk payload after 5 days of BSD selection (8 μg/mL), nucleofected using either program CB-150 or CB-137 on the Lonza 4D Nucleofector, and with either 1 μg, 2 μg or 3 μg of NKX6-3-Risk payload and 1 μg of a dual gRNA-Cas9 plasmid. The condition with the most surviving colonies was the 1 μg of payload DNA using CB-150. **C.** AS0041$^{MC1}$ cells following transfection with NKX6-3-Risk haplotype payload, and a dual gRNA-Cas9 plasmid after 6 days of blasticidin selection. Cells that did not integrate the payload remain tagged with JF549 HaloTag Ligand, shown in red, and cells that integrated the payload are shown in green, expressing mNeonGreen as a part of MC2. **D–E.** Single cell sorting data of AS0041$^{MC1}$ and nucleofected NKX6-3-Risk, showing 93.98% JF549+ cells in AS0041$^{MC1}$, and 26.15% JF549+ and 26.03% mNeonGreen+ cells sorted for NKX6-3-Risk. Gating for the AS0041$^{MC1}$ condition was performed on all events collected, thus the percentage of live cells shown is relatively high, whereas gating for the NKX6-3-Risk condition was performed on a subset of events collected, thus the percentage of live cells shown is relatively moderate. **F.** Percentage of PCR-screened colonies after sorting. A total of 84 colonies were PCR screened for left (L) and right (R) junctions, and inner regions of *ANK1* and MC1 to ensure proper payload integration.

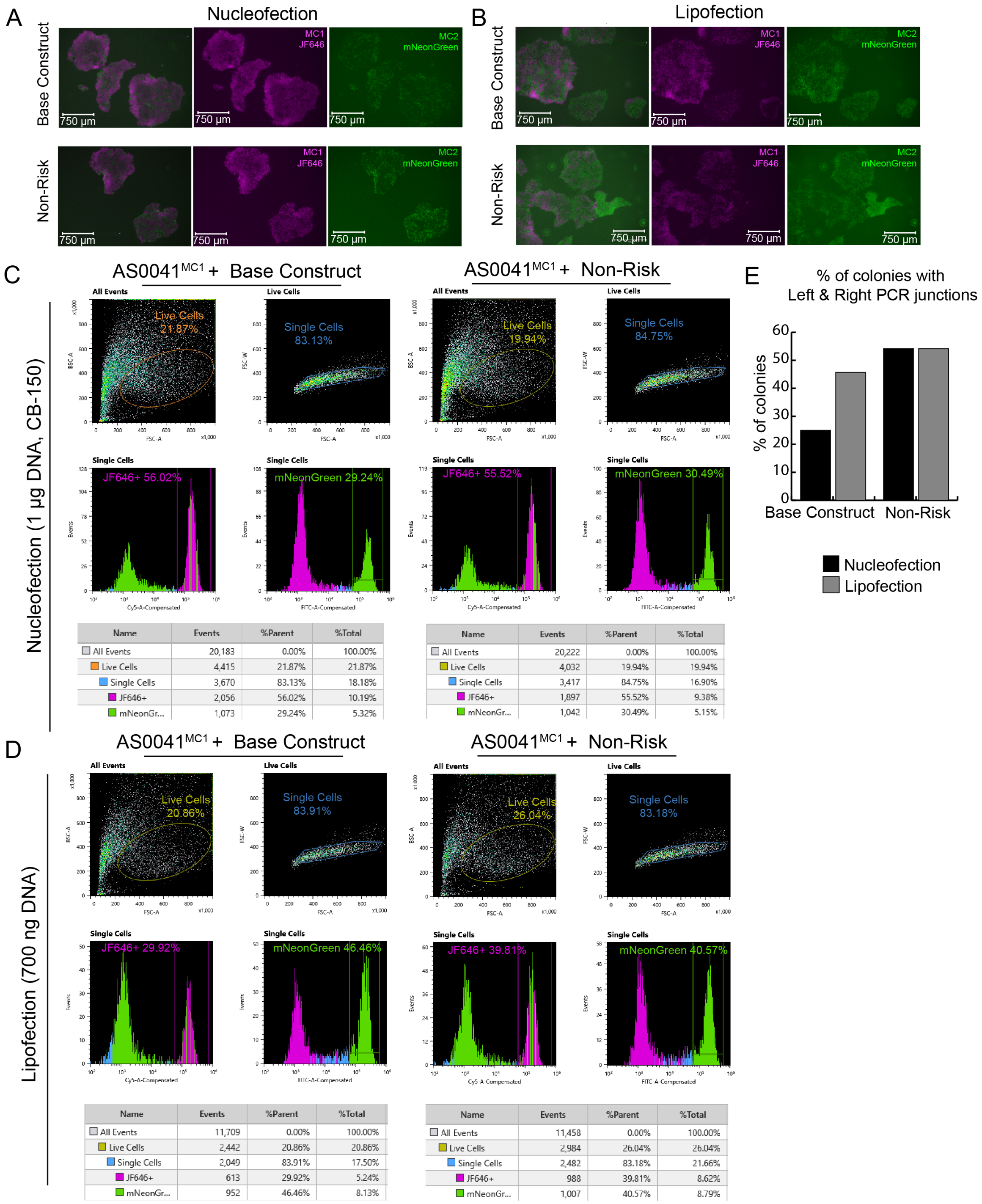

**Fig S10. A.** AS0041$^{MC1}$ cells following nucleofection with program CB-150 and 1 µg of Base Construct payload + dual gRNA-Cas9 plasmid, and 1 µg of Non-Risk payload + dual-gRNA-Cas9 plasmid after 6 days of blasticidin selection. Cells that did not integrate the payload remain tagged with JF646 HaloTag Ligand, shown in magenta, and cells that integrated the payload are shown in green, expressing mNeonGreen as a part of MC2. **B.** AS0041$^{MC1}$ cells following lipofection with 700 ng of Base Construct payload + 1 µg dual gRNA-Cas9 plasmid, and 700 ng of Non-Risk payload + 1 µg dual-gRNA-Cas9 plasmid after 6 days of blasticidin selection. Cells that did not integrate the payload remain tagged with JF646 HaloTag Ligand, shown in magenta, and cells that integrated the payload are shown in green, expressing mNeonGreen as a part of MC2. **D–E.** Single cell sorting data of AS0041$^{MC1}$ and nucleofected NKX6-3-Risk, showing 93.98% JF549+ cells in AS0041$^{MC1}$, and 26.15% JF549+ and 26.03%% mNeonGreen+ cells sorted for NKX6-3-Risk. Gating was performed on a subset of events collected, thus the percentage of live cells shown is relatively moderate. **F.** Colony counts of NKX6-3-Risk transfected cells that grew up in 96-well plates post-single-cell sorting, and colony counts of NKX6-3-Risk transfected cells that survived after BSD selection (8 μg/mL) post sorting.

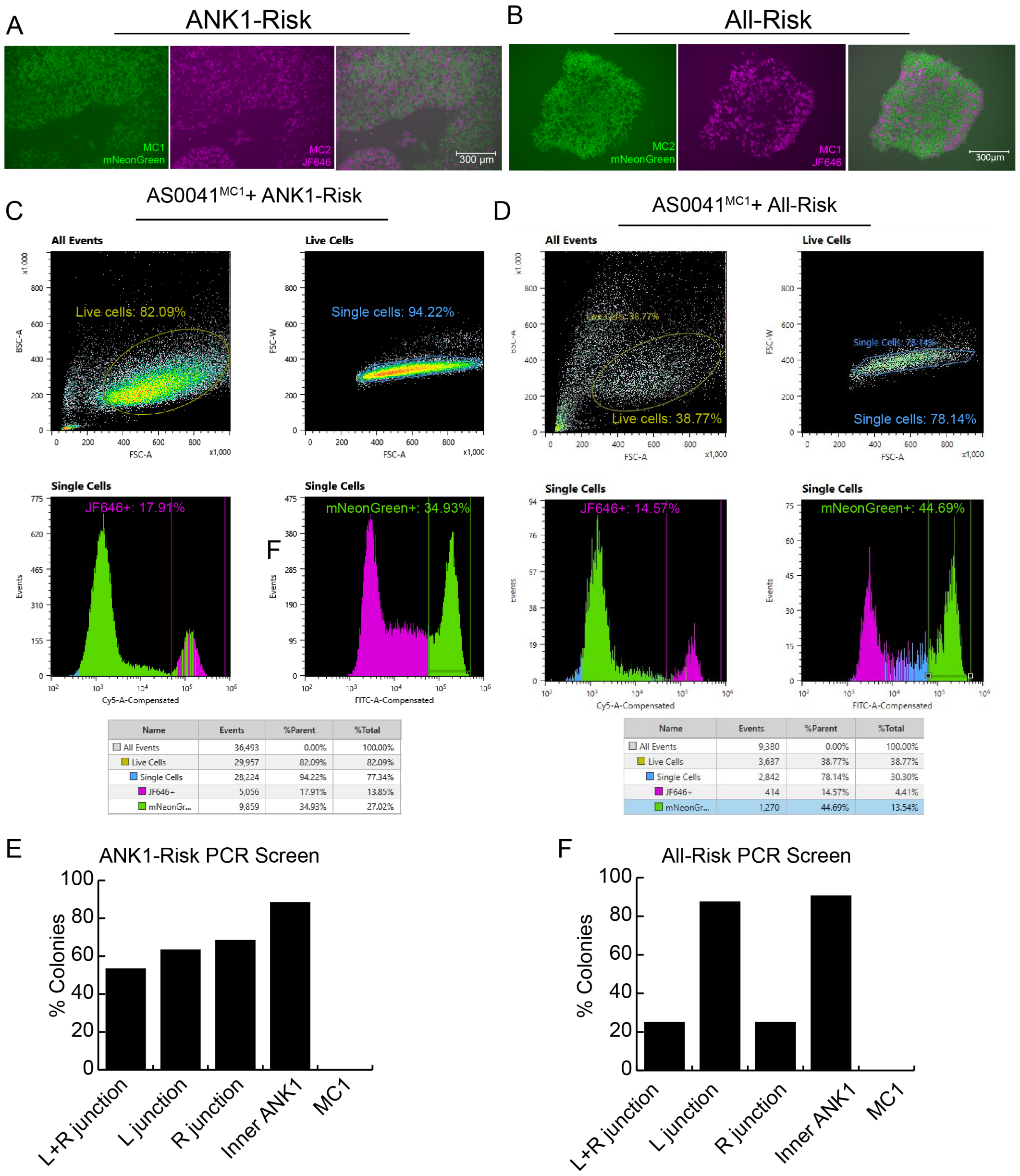

**Fig. S11. A–B.** AS0041$^{MC1}$ cells following lipofection with ANK1-Risk (A) and All-Risk (B) haplotype payload, and a dual gRNA-Cas9 plasmid after 6 days of blasticidin selection. Cells that did not integrate the payload remain tagged with JF646 HaloTag Ligand, and cells that integrated the payload are shown in green, expressing mNeonGreen as a part of MC2. **C.** Single cell sorting data of AS0041$^{MC1}$ lipofected with ANK1-Risk haplotype payload, showing 17.91% of JF646+ cells in the single-cell population, and 34.98% mNeonGreen+ cells in the single-cell population. Gating was performed on all the events collected, thus the percentage of live cells shown is relatively high. **D.** Single cell sorting data of AS0041$^{MC1}$ lipofected with All-Risk haplotype payload, showing 14.57% JF646+ cells in the single-cell population, and 44.69% mNeonGreen+ cells in the single-cell population. Gating was performed on a subset of events collected, thus the percentage of live cells shown is relatively moderate **E-F.** Percentage of PCR-screened colonies with both left (L) and right (R) junctions, left (L) junction only, right (R) junction only, inner *ANK1* region, and inner MC1 region in candidate ANK1-Risk and All-Risk cells. No cells retained MC1. A total of 60 colonies were screened for the ANK1-Risk haplotype and a total of 32 colonies were screened for the All-Risk haplotype.

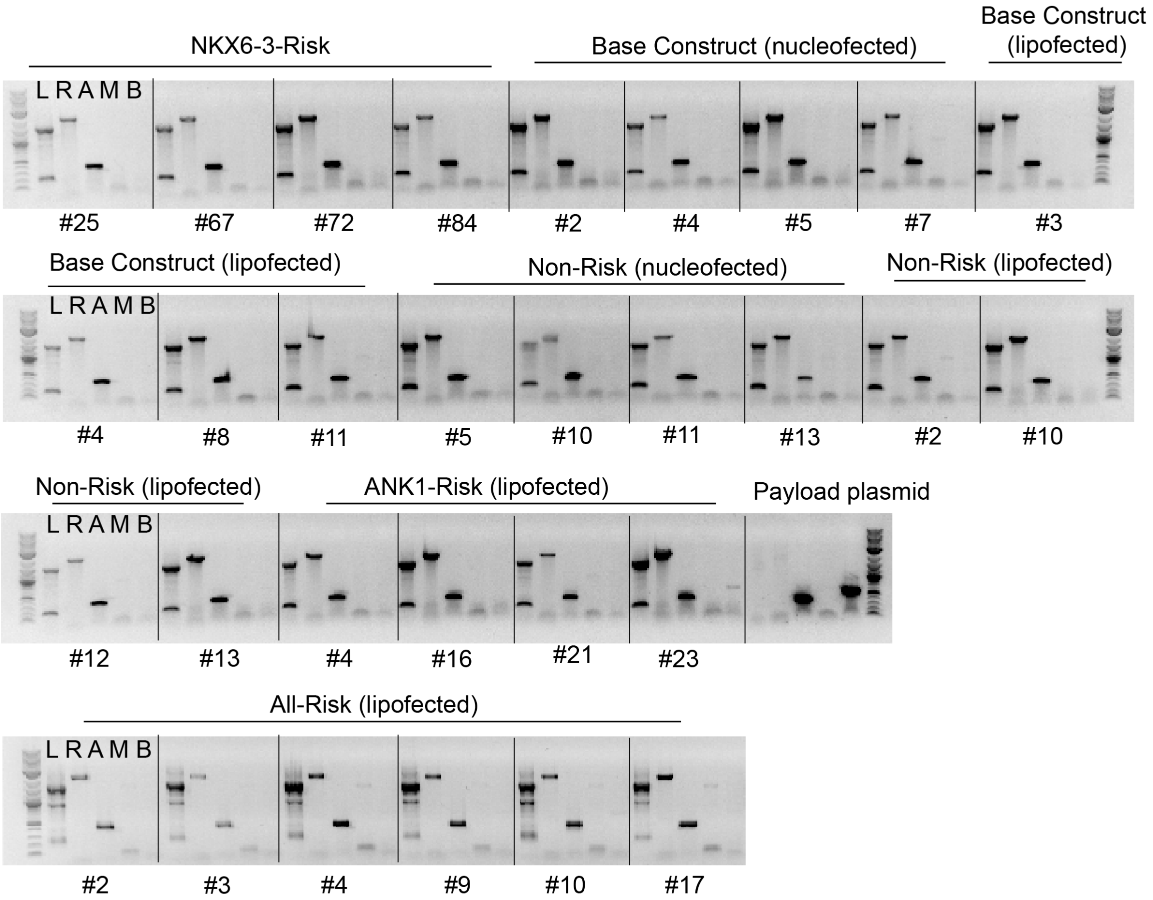

**Fig. S12.** PCR genotyping of candidates for Capture-sequencing. Primers targeting the left (L) and right (R) junctions used to check for correct payload integration, primers targeting the inner portion of ANK1 (A) were used to test for *ANK1* presence, primers targeting MC1 (M) were also used to ensure MC1 loss, and primers targeting the payload backbone (B) were used to test for backbone integration. All candidates shown had both left and right junctions, *ANK1,* and no MC1 or backbone integration. Primer sequences can be found in Table S4.

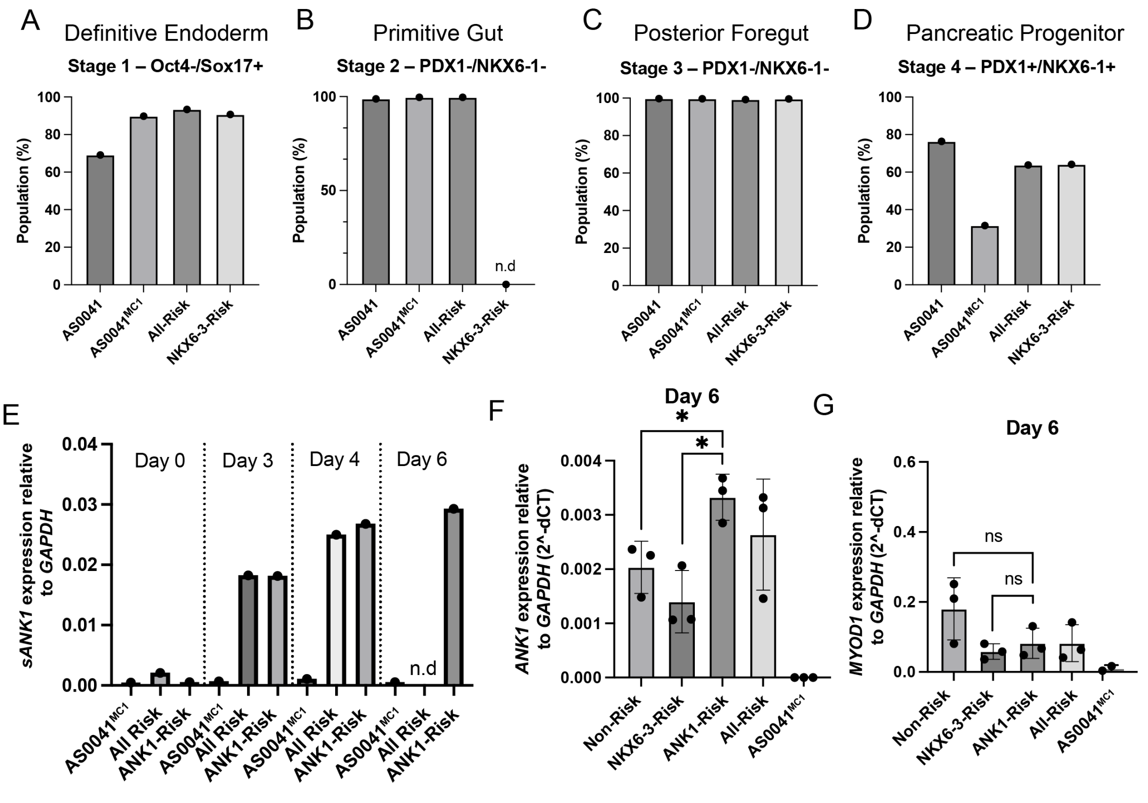

**Fig S13. A-D.** Flow data from pilot pancreatic differentiation of AS0041, AS0041$^{MC1}$, All-Risk, and NKX6-3-Risk haplotypes at Stages 1-4 of pancreatic cell differentiation. AS0041$^{MC1}$ showed lower levels of NKX6.1+ and PDX1+ cells at Stage 4 than the other haplotypes, but integration of haplotype payloads restored PDX1 and NKX6.1 levels to near parental levels. **E.** Time-course experiment of skeletal muscle differentiated cells at Days 0, 3, 4, 6 showing highest *ANK1* mRNA at Day 6. **F.** Quantification of *ANK1* mRNA at Day 6 across all haplotypes, showing highest *ANK1* expression in ANK1-Risk compared to Non-Risk and NKX6-3-Risk. Each data point represents a second biological replicate that has been differentiated three separate times. *P*-value ANK1-Risk vs. Non-Risk = 0.026; ANK1-Risk vs. NKX6-3-Risk = 0.026 (Student's t-test with Welch's correction). **G.** Quantification of *MYOD1* mRNA at Day 6 across all haplotypes, showing no significant difference in expression. Each data point represents a second biological replicate that has been differentiated three separate times.

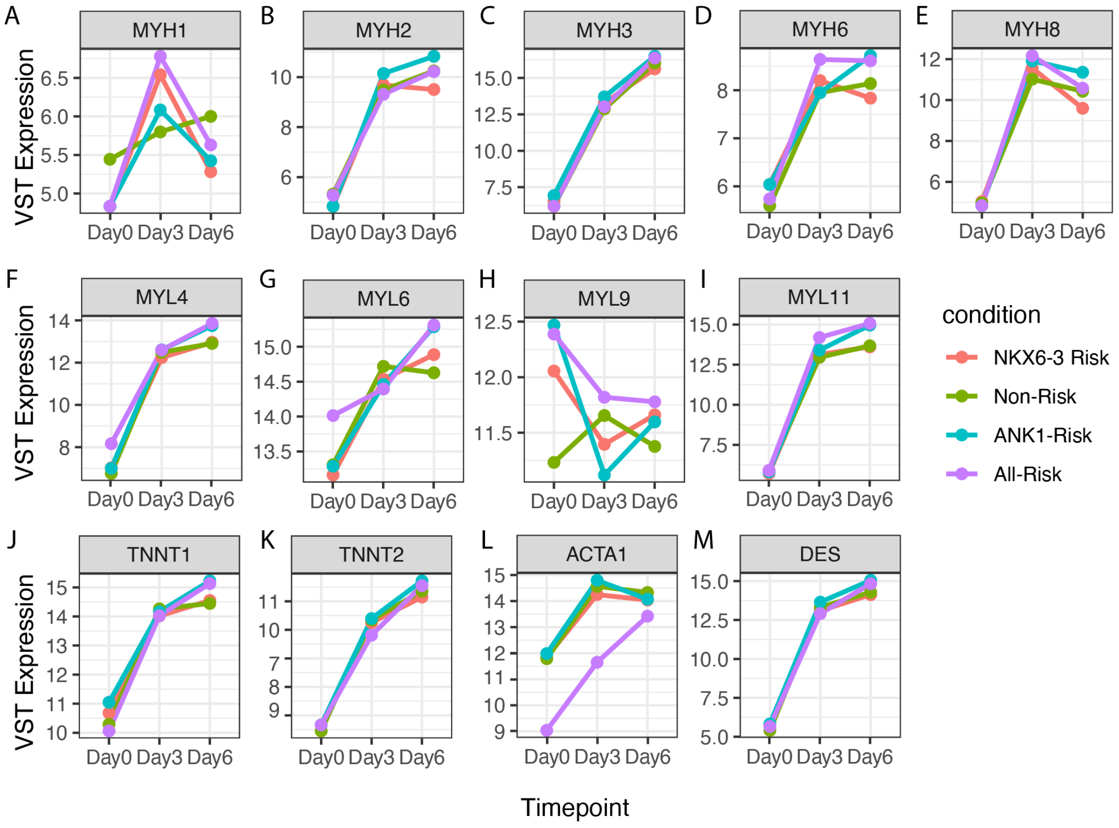

**Figure S14. A-E.** Gene trajectories for Day 0, Day 3, and Day 6 for myosin heavy chain genes showing dysregulation of *MHY1* in risk-SNP haplotypes compared to the Non-Risk SNP haplotypes, and no major differences in *MHY2, MYH3, MYH6,* and *MYH8.* **F-I.** Gene trajectories for Day 0, Day 3, and Day 6 for myosin light chain genes showing differences in *MYL6, MYL9,* but no major differences in *MYL4* and *MYL11* for the All-Risk haplotypes compared to the other haplotypes. **J-M.** Gene trajectories for Day 0, Day 3, and Day 6 for sarcomere structural proteins that are important for muscle development and function. The All-Risk haplotype showed a great difference in the expression of *ACTA1* compared to the other haplotypes.

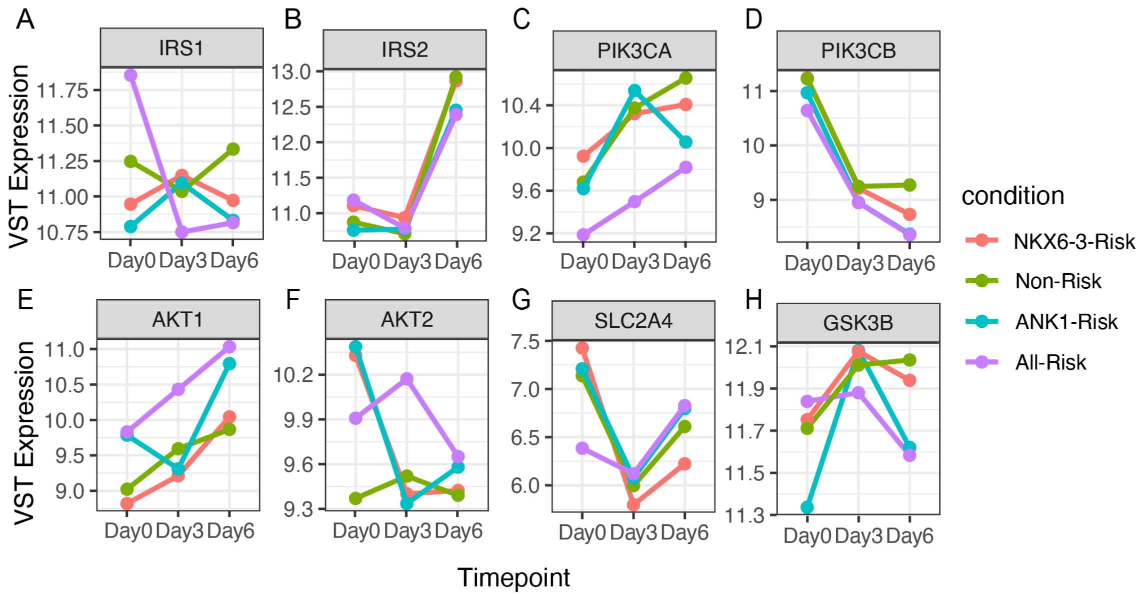

**Figure S15**. **A-D.** Gene trajectories for Day 0, Day 3, and Day 6 of genes *IRS1, IRS2, PIK3CA, and PIK3CB* which are important in the early stages of insulin signaling. All-Risk exhibits differences in *IRS1* and *PIK3CA* gene expression compared to other haplotypes. *P*-values were calculated using a two-way ANOVA with Tukey's multiple comparisons test. *PIK3CA*: *p*-value = 0.0217; *PIK3CB*: *p*-value = 0.0252; *AKT1*: *p*-value = 0.0346. **E-H.** Gene trajectories for Day 0, Day 3, and Day 6 of genes *AKT1, AKT2, SLC2A4, and GSK3B* which are important in later stages of insulin signaling. The All-Risk haplotype shows gene expression differences in *AKT1, AKT2,* and *GSK3B* compared to the other haplotypes.

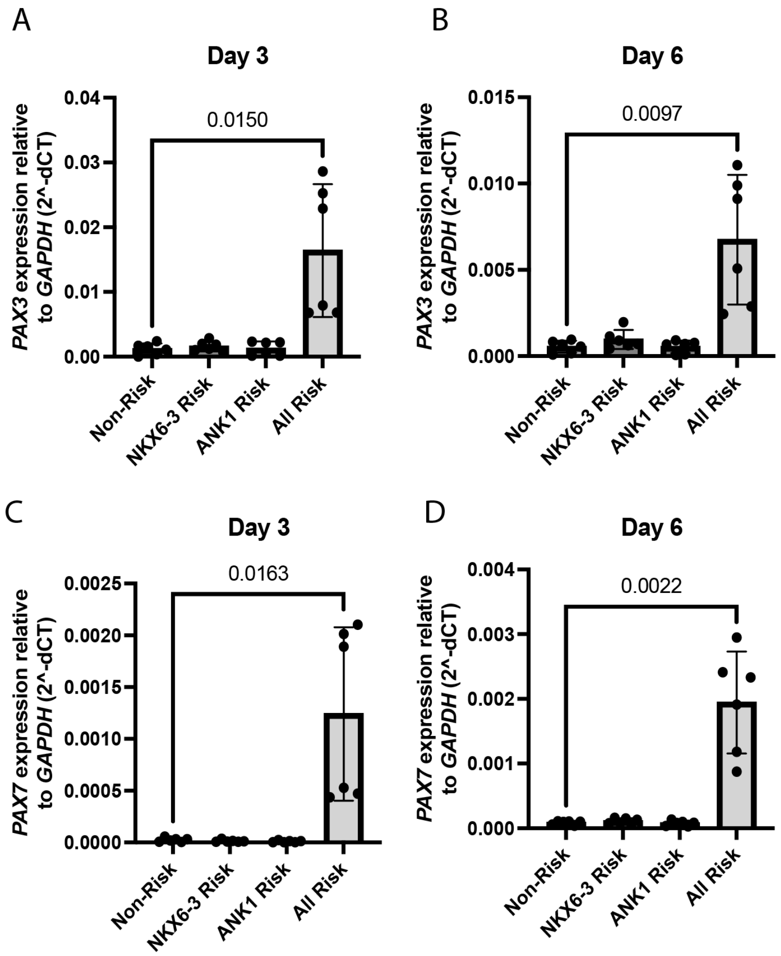

**Fig. S16. A.** Quantitative measurements of *PAX3* mRNA expression relative to *GAPDH* 3 days post *MYOD1* infection, showing greater *PAX3* levels in the All-Risk (p = 0.015). **B.** Quantitative measurements of *PAX3* mRNA expression relative to *GAPDH* 6 days post *MYOD1* infection, showing greater *PAX3* levels in the All-Risk (p = 0.0097). **C.** Quantitative measurements of *PAX7* mRNA expression relative to *GAPDH* 3 days post *MYOD1* infection, showing greater *PAX7* levels in the All-Risk (p = 0.0163). **D.** Quantitative measurements of *PAX7* mRNA expression relative to *GAPDH* 6 days post *MYOD1* infection, showing greater *PAX7* levels in the All-Risk (p = 0.0022). P-values were calculated using Student's t-test with Welch's correction. Dots represent two biological replicates, which have been differentiated three separate times each. Technical replicates cluster together.

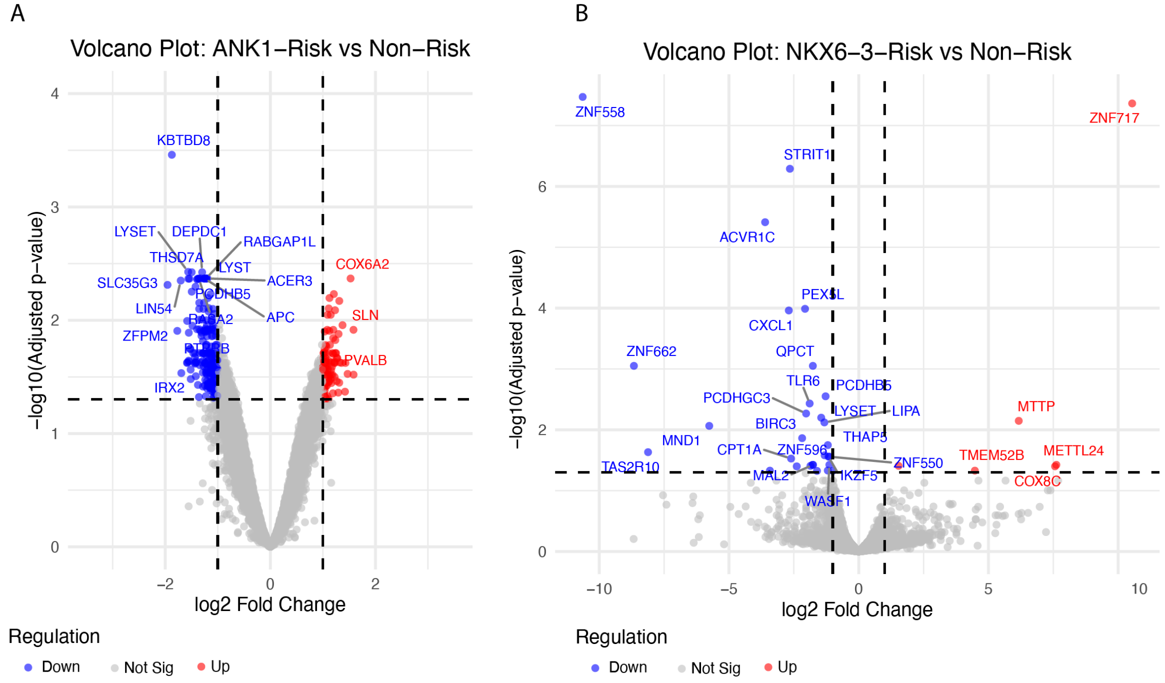

**Fig. S17. A**. Volcano plot for ANK1-Risk compared to Non-Risk 6 days post *MYOD1* infection showing, downregulated genes (blue) and upregulated genes (red), plotted with adjusted p-value cut-off of 0.05 and absolute log-fold change of 1. **B.** Volcano plot for NKX6-3-Risk compared to Non-Risk 6 days post *MYOD1* infection, showing downregulated genes (blue) and upregulated genes (red), plotted with adjusted p-value cut-off of 0.05 and absolute log-fold change of 1.

### Tables

**Table S1.** Proposed SNPs for each Haplotype Build

| | RS_Number | All-Risk | ANK1-Risk | Non-Risk | NKX-Risk | Base Construct |
|---|---|---|---|---|---|---|
| 1 | rs5891165 | - | AAAG | AAAG | - | AAAG |
| 2 | rs35659384 | G | - | - | G | - |
| 3 | rs4736999 | G | C | C | G | C |
| 4 | rs12549294 | G | A | A | G | A |
| 5 | rs7825337 | T | C | C | T | C |
| 6 | rs7825494 | T | C | C | T | C |
| 7 | rs6981607 | C | A | A | C | A |
| 8 | rs12549902 | A | G | G | A | G |
| 9 | rs4737000 | C | A | A | C | A |
| 10 | rs4736819 | T | C | C | T | C |
| 11 | rs12550613 | C | G | G | C | G |
| 12 | rs57963234 | - | ACAGCAGA GTCTAT | ACAGCAGA GTCTAT | - | ACAGCAGA GTCTAT |
| 13 | rs3063734 | GT | - | - | GT | GT |
| 14 | rs17659386 | C | T | T | C | T |
| 15 | rs17659428 | C | T | T | C | T |
| 16 | rs12544241 | A | G | G | A | G |
| 17 | rs6990613 | G | A | A | G | A |
| 18 | rs716199 | G | A | A | G | A |
| 19 | rs10504042 | A | G | G | A | G |
| 20 | rs516946 | C | C | T | T | T |
| 21 | rs515071 | G | G | A | A | A |
| 22 | rs146025758 | AAG AG | AAGAG | - | - | AAGAG |
| 23 | rs508419 | G | G | A | A | A |
| 24 | rs9694034 | A | A | G | G | A |
| 25 | rs6989203 | G | G | A | A | G |
| 26 | rs11354309 | T | T | - | - | T |
| 27 | rs28591316 | A | A | G | G | A |
| 28 | rs28602970 | A | A | G | G | A |
| 29 | rs35273426 | C | C | T | T | C |
| 30 | rs10109812 | A | A | G | G | A |
| 31 | rs750625 | C | C | T | T | C |
| 32 | rs3802315 | G | G | T | T | G |
| 33 | rs3802316 | G | G | C | C | G |
| 34 | rs59864562 | A | A | G | G | A |

| | | | | | | |
|---|---|---|---|---|---|---|
| 35 | rs10106563 | C | C | A | A | C |
| 36 | rs13266210 | A | A | G | G | A |
| 37 | rs11995075 | T | T | C | C | T |
| 38 | rs62508166 | G | G | A | A | G |

**Table S2.** SNP Positions and Allele Frequencies

| RS_Number | Position (hg38) | Allele Frequency |
|---|---|---|
| rs5891165 | chr8:41646335-41646338 | -=0.580, AAAG=0.42 |
| rs35659384 | chr8:41647443-41647445 | GGGG=0.549, GGG=0.451 |
| rs4736999 | chr8:41648633 | G=0.549, C=0.451 |
| rs12549294 | chr8:41648861 | G=0.581, A=0.419 |
| rs7825337 | chr8:41649718 | T=0.549, C=0.451 |
| rs7825494 | chr8:41649831 | T=0.549, C=0.451 |
| rs6981607 | chr8:41650649 | C=0.578, A=0.422 |
| rs12549902 | chr8:41651740 | A=0.570, G=0.430 |
| rs4737000 | chr8:41652208 | C=0.570, A=0.430 |
| rs4736819 | chr8:41652396 | T=0.538, C=0.462 |
| rs12550613 | chr8:41652741 | C=0.571, G=0.429 |
| rs57963234 | chr8:41655068-41655081 | -=0.527, ACAGCAGAGTCTAT=0.473 |
| rs3063734 | chr8:41655152-41655153 | GT=0.53, -=0.470 |
| rs17659386 | chr8:41656012 | C=0.559, T=0.441 |
| rs17659428 | chr8:41656290 | C=0.559, T=0.441 |
| rs12544241 | chr8:41657369 | A=0.559, G=0.441 |
| rs6990613 | chr8:41657611 | G=0.559, A=0.441 |
| rs716199 | chr8:41658012 | G=0.559, A=0.441 |
| rs10504042 | chr8:41658474 | A=0.559, G=0.441 |
| rs516946 | chr8:41661730 | C=0.769, T=0.231 |
| rs515071 | chr8:41661944 | G=0.767, A=0.233 |
| rs146025758 | chr8:41664499-41664514 | AAGAG=0.791, -=0.209 |
| rs508419 | chr8:41665473 | G=0.757, A=0.242 |
| rs9694034 | chr8:41665840 | A=0.759, G=0.241 |
| rs6989203 | chr8:41666227 | G=0.759, A=0.241 |
| rs11354309 | chr8:41666242-41666243 | T=0.759, -=0.241 |
| rs28591316 | chr8:41666907- | A=0.759, G=0.241 |
| rs28602970 | chr8:41666987 | A=0.759, G=0.241 |
| rs35273426 | chr8:41667170- | C=0.759, T=0.241 |
| rs10109812 | chr8:41667381 | A=0.759, G=0.241 |
| rs750625 | chr8:41668396 | C=0.759, T=0.241 |
| rs3802315 | chr8:41670660 | G=0.758, T=0.242 |
| rs3802316 | chr8:41670710 | G=0.758, C=0.242 |
| rs59864562 | chr8:41671654 | A=0.797, G=0.203 |
| rs10106563 | chr8:41675682 | C=0.784, A=0.216 |
| rs13266210 | chr8:41675996 | A=0.784, G=0.216 |

| | | |
|---|---|---|
| rs11995075 | chr8:41676550 | T=0.785, C=0.215 |
| rs62508166 | chr8:41679800 | G=0.798, A=0.202 |

**Table S3.** Gene fragments and Linked DNA Sequences

| Gene fragment ID | Sequence | Description |
|---|---|---|
| gNCH004 | ttccccactcacctgcccagctgagaccgaggtgctctgagataaca gccatcttcatctctgcctgctctgtcccactgagagaacctggggaga agcattagatttcatttagtagccagggcaggacacagttgacaggcc tgaggacacggcctgtgatggatgtggcttccgtgtgcactggggtgat tgtctagactctctgcaagatataacttcgtatagcatacattatacgaa gttattgcagtgcttcagccgctaccccgaccacatgaagcagcacg acttcttcaatttagggtagcatcaggaatctgaaccctcagaaagtgg ggatcccgggtatagacctttatctgcggttcaagttaggcataaggct gcatgctaccttgtcacacctacactgctcgaagtaaatatgggaagc gtgcgacctggctccaggcgttccgcgccgccacgtgttcgttaactgt tgattggtggcacat | Right linker for DNA for Base Construct assembly with homology to RP11-111B9 |
| gNCH006 | ggcctaaggtagctaccaatatttagtttctaagccttgcgacagacctc ccacttagattgccacgcatagagctagcgagtcagcgaaaagcat gacgcgctttcaagcgtggcgagtatgtgaaccaaggcttcggacag gactatatacttaggtttgatctcgccccgagaactgtaaacctcaacat ttatagattattgcagtgcttcagccgctaccccgaccacatgaagcag cacgacttcttcaaataacttcgtataggatactttatacgaagttattctc tactaaaaatacaaaaattagccaggtatggtggcttgtgtctgtagtcc cagctactcgggaggctgaggcaggagaatcgcttgaacccagga ggtggaggttgcagtgagccgagatggtgccactgcactccagcctg tgtgacagagcaagacttcatctctaaataaataaatagataaaataa aataaatttgcacattatacatctagtgaagggctatccaagatagata aagaactcaaacagaaggccgggcgcggtggctcatgcctgtaata ccagcactttgggaggctgaggcaggtggatcacctgaggtcagga gtttgagaccagcctggccaatacggtgaaacccatctctactaaaaa tacaaaaattagacgggcatagtggcgggtgcctgtagt | Left linker for base construct assembly with homology to RP11-111B9 |
| gNCH007 | aggagaactaggcgcccaggagggcctgatggaagaaacggacg ggccgcctgtggtgaggagtgcacacaggcgctggccctgcctggg gcacatacttggcaggggagcattctaaataaaacctgactgaattcc tagcagttctatggctggcagctcagggccttccttccagagccaccct agtcctggagtctaaggttgaggtaggaaagaagggagacaagga aagaaagagaggaaggagaggagaggaggaagcaccatgcctg gatgaggcgtgaaaacagagacggaacatgtcgttcatgtacatact aaacacacgcatgtggagaacagtggtgcatgttttataagaacaca aatggaaaagacacccattaatacgatggccaggatatgcttccaaa taatagtcagtgggagaaagtggggggtgcagatgaaacaagattg gccacgagtagatcatcatggggcctgggaatggggaa | Non-Risk and NKX6-3-Risk nonrisk SNPs rs35273426 rs10109812 |
| gNCH008 | gatacgttgtgacacatgtgtacccccatgaagccatcaattcaaaga agaaaaaggacatttgtcaccccagaaaggtctcctcaagcttctttgt aatctctccctgccctttccccctctgccaaccccaaaccattcccaggt gaccccgatctgctttttgccactacacatcagtttgcatttcccaggattt tacataaacggattcatacagtatgcactctctctttctttaaagcctggct tctttcactcaacgtagtcaatttgggtttcatccatgctgatgcatgtatc aataatttatttctgttgctaagccatagccctttggaaggatatatcacg gtttgtaaatttatttacctgccaatggacatttgggctgcttccagtttgtg gctattacaagtaaagatgctagtctgtatggacataatgctttcttttctct tgggtaaatagctaggaatggaatgacttgattgtacagtaggtatatgt ttaactttgtcagaaaatgatcaagtggttgtaccattttgcattcccacca gtggtgtatgagagttccagttcccccatatcctcaccaacacttgatat ggccagccttttttaattttaaccattcttttaggtatgtggtggtacctcctta tagttttaatttgtatttcccaaatcactaatgatgttgagcatattttcatgt gcttatttgccatgtgtatatattctgggtgaaaagtctgttcgaatctattg ctcaatctaaaaactgagttgttttcttattattaagttttgagagttctttgta | Non-Risk/NKX6-3-Risk rs10106563, rs13266210, rs11995075 nonrisk SNPs |

| | | |
|---|---|---|
| | tattccaatactatagcagatgattatttgccaatattttctcccagtctgtagctcgtcttttcattttcttaacagtgcattttgaagagcagaagttttgattttgatgaagcccaacttgtcaatttgttcttttatggattgtgcttttggtgttgtatctgagaaat | |
| gNCH009 | ggctgacgtgcagatcatctgaggtcaggagtttgagaccagcctggccaacatggtgaaaccctgtctctactaaaaatacaaaaattagcccggtgtggtggcaggtgcctgtaatcccagctactcaggaggctgtggcaggagaatcacttgaacctggtggatggaggttgcagtgagctgagattgcatcactgcactccagcctggttgacagagcaacactctgtgtcaaaaaaaaaaaggcggggggctggtcaagccctcaagacagagcttcagaacagcagttcccaggaatgagctccgagggggaggtagcttttgtcctaggagcccctccttgagaggccaagaagagggacagcctttgccccattgcttaatggaaaaccctctgctagacagccctgaaggggccttggcagtgtcatttcatgaccaaggtgtgactgggtgtgacctccccctcgagcccacagccttctgttttgctttttgatatggatgggagtttgtgctcttccccattctactcccatgagcctaagggcagagggcatctctttttcacccacccattcccgtattcacctgtccaaatcttcaaaccctttactctgtatctgccgtgtgctctgagtgggaggtgaaacatcagtgagacactgtccctgcctgcaagcagctcacagtgcaggaggcaagacagacgcacaaggagccttgctcaagggtgatgagagctgcgcctaagcagaagcacattcgagatcctgcctcagctcctcctctattttacccatagctagcaccgtgccccacacttgacaggcaaggtcatgtctctgagttaatatctgctaccaatccaatgcattttcacttccttttaattatttaattattgtgatggggtctcactctgtcacccaggctggagtgcagtggctcaatcatagcttatgcagacttgaactgctgggtttaagtgatccttctgcctcagcttcctgagtagctgtgactacaggcatgcgccaccatgcctggctaattttgtgtgtgtgtgtgtctctctctctctctgtgtgtgtgtgtgtgtgtgtgtgtatttatttatttttttaaaattttttttagagatggagtcttgccatattgcccaggctggtctaaatgtctgggctcaagcgatcctcctgccttggcctcccaaagtgctgggatcacaggtgtgagccagcacacccggcccacattctttttatttgaggaggcccatggtaggccagggatgctcaaactgcttgtttaacagctgttttcagggttaggcatcatgggactaggtctgggatgtttcctgggggctggcccttctggaaaagagcagcagaagctattcaggttgactcagacaaagcggttgtgaggacaccatgactggactgcagatgaccaaagctcccagctcctgggccctccatccctcctgagcacgggggcagccagcctggctgtgctcacctgctgtgagggcagcagggagaagccactggcttctaccaccttttagcttctagcccacctgcctctccccagcacagaggggaacactctgcacccaccttcttggtgacaatgttgccctgctcatccgtgaattgctcctctgtcacctgctcccctggaatattctgaaactcattcccctggaattagagaaagggagaaaatgcaagattggtgagtgggagtctgggaggaagggtggaggacgaggaaatgaaacagcagtagccacaatagccatggcccaataactcccccagcaggaaacccggctcagcatgtagcaggagccatgccagggctcccagggcgtgggcggcgcccctgagggtctgggaggcctgaagacgaacggtcgagctcacaattgtgcacttggccctcacaacacccccctgggaacctgcaaaatgggggcaattatgcaagtctgacggaggaggccctgagagaggaagtgaggcacctgtttcctatttttacagcactttgcagtatagaaaagcccttcgacaggcagcatcacccagaatgtgacttgcttgaaaccccagagccagttggtgacagaaacaggacgatccgggtgcaatcccatggtggct | Non-Risk/NKX6-3-Risk Nonrisk SNPs for rs146025758 |

| | | |
|---|---|---|
| | cacgcctgctatcccagcactttgggaggccaaggccggaggacca cttgagcccaggagttcaagaccaatttgggcaacatagcgagcccc catctctacaaaaatttttttaaaatagccaggcatggtggtgtgtgcctg tagtaccagctactcagaaggctgaggcaggaggatcacttgagcc caggaatttacagtaagcccaggttacagtgagccatgagcatgcca ctgcactccagcctggactaaacagcaagacttcatctctaaaaaca aaagaaaaaagagaagagagagaaagaagaaagaaacaaag aaagaaacaggatgagcgcagcctcccgagacccaaagctgagt gtttaatgtcactgtcacaatgca | |
| gNCH010 | aatgtcactgtcacaatgcactgggcaccagagggcaggttctggtg ggcgcacacacactctggctcccagggtggaacatgatccctgggg gcctcctggtccttttcctccggcgtctcccaactctgggtccggagctc cccacagcagcgtcccctcagccccggcctccggcgcccacacca ccagccctgccggcctcctgccaggcggttaccttcaggaagacccg ccgccggaccaccctggtggagatggtctcctcgtcgtcactgaggcc ctcgctctcccccagctccttgtccagctcctggtggatgtgctttagcac aaagcacagggacccccctgacaatgtgcatcacgttctgacagctga ccaggaagaagctcagcagcaccagcgtgaccaacagctgggtga cgaaagtccacatcctcgcctcctcaccagcccggtcctctggcagcc agggccccggccaccacggggggcctgcctctcattctccactggga ctcctgagtcccccagggacccccttgcattccccctccctatctctctggt ttgctctcttggtctctcactgtttctctctctgtgggcaggacaccgaatg gccctctcggctgaagcagcccggcccaagtgcccttctgcccagca gccaggctctgccctgagtggcccgggtgacatcacgctcctataattt tacctccccaaaccagctgccggagctgtccaaccgggctagaagg aagccggcaggtgcccatgtcctgagcactgaaagcacaagtgaa agtcactgtgtgagtgacacccgctgctgcctaaccccgccctgccca gagacctggaagtgggggctcggcctctcacctatcctgcccatcccc atgggaactagaaagagttctgcccagggaaagtttcttactttcagaa aataatacagtccgagaaagcagctacttgtgtgtctccttgggctgct actgcggtgatccaaatgctgttccccggcatggtagggatgagacc agctaataccctgggtacttagcggtctgtttcaaaaattgctccaaaa gggtgagtgatgggctcagggacagtcaagtgagcccagcagggg cccagcacaggtggtggggccacctggaagccggtctgtaattcctgt cagcccacaggcccttgcacccttcctttatggggtacagcctcccaa aactgaagcatggggctggtgtgtctatttaaggcctgttcttaaaggct cttccagggctgggctatcaacgcccactgagcaggaacaacgcgt attaaacttgtatttacagagctgattcacaaatgtagccgcagcagct gcggctggaaaaaaatttcactttctagatcaggttgttcctctctgagtgg caatgcagcatggcagaaagaacacagatctgggacacagcaga ctaaccctgcaactcacctactgtggggtcttgggaaggctggggacc ttctctgtgcctcagtttcctcatctgtgaagcacggattaaagtgctctcc ccaccctttctcaaggctgctgggacaatcatgccgcttatgaatgcta acactatgaataataatggcaaacgctgtgatagtgcatggttctatgt ggatttcgcattaattcactgacacctcaccacagtcctaaccaagga ggttctatcgtcttcccattttacacaaaggaaacagaagcaaagtatt ctgcctaggtggaggagtaagtaaacggaggagtcacattccaaatg | Non-Risk/NKX6-3-Risk<br>Nonrisk for rs508419<br>rs9694034<br>rs6989203<br>rs11354309 |

| | |
|---|---|
| | cctctcactccacaatgctgcttatttcaacaagaaagcacggggaaa<br>agtatgtgcaaagaagccggggaagcatctggttttggcccagtcaa<br>gaagagtccaacttcagatatttcctcctaagtcctcagagaaatcagc<br>tgggtcacagtcatttcaggccacctcccaagtcccaagttacatgca<br>atctctctatctccatcactgttttcagaggtccagtggaaatactaataa<br>gtgggtgctttgcaaaccatcaatcaccttcaagggcatccttatcagc<br>cctcggggcctgtgagatgggcaccggcagcagcctgccaggccct<br>ctgatcggcgtctccagttccaggctgggcat |

| | | |
|---|---|---|
| gNCH011 | tccctcctttaccagtgaggaccctgaggtgctgggaagagagcgcct<br>tgggcacgtcacccaggtcaccataggaggagagtaagcccaacg<br>cccagtgctcggttcctagccctttctaaaacgcgacagaacaagaa<br>aaccaacaaaagccacaactttggagtcagcgagacccagcttaga<br>gcccacttctgaaactgtgatcttgagcaagttgctcagccactctgatc<br>agcagctgcccaacaggtacaggcgccttagcagggatccaggag<br>gcagaaaggcaggccttgcccttgacttctagagaaacggaagcac<br>caccacaagtgttaaaggaaggggatatgcttagctcagggcttgagt<br>gcctgagaggcagccctggagtcccggccccttccgatgctgggaa<br>ggaacagcagcacgcagctcccctcgggctctcacctgcaccacttg<br>ggtgaaggtgttcttggcctcctgcacctgctcttcttggccttgctgcctc<br>cggtccctgtcggcctgggagctctcagcctcgggctgttctgtccacgt<br>gtgctcacttacagataccaggaccttctcgtactcctgagatccaccg<br>ggctctagcccttcagtcatggtacttgttccctggaatgagtgtggacct<br>tgcgtgacctcctcttgccaggaaccttgtgcagcatcttgcaggtatga<br>gacaatcgagccgtctgcatcccagtcctgcagtctggggtccagaa<br>gaagcagcagatggccggccggggagagagaaaagacacctggt<br>cacccatcagtctcattccttttagacactctccccacccagagctctgg<br>ctcatcaaggacacagaccgtcctcccatccctccaaaggtcagggg<br>cgcatgagcacattgctcacctcactgctgtggttcagaggggtacac<br>gtggcccatctaatgctgtccaggtggaaaggccactggtccctccct<br>gtgggtgaacagcttgcccttggggctcccagagcacttggctcaccc<br>taatggtccaaacactggggcctccacactctcatcccacatcctcaa<br>actctgccaccacctccttctgctctccaggcccttcccactcagcagtc<br>ctcatctgacaaggtggtcattctctccccctcaagacacattctcccgt<br>ggctcctggaaggtgctagcctggcgtctccccccttcctctctggccgct<br>gttgttccatgtcctctgcttgttcctcctagctttcctgacttctgaacatag<br>gagcaccccggggtgcagttctcagacctcaccgctgtccacatcca<br>cttcctaagcaagctcattcatcccatggttccaaagcccatctctacac<br>tcagcgtgcctggatgtgtgtctccagcttggctctcctcctagacctac<br>agctgtgtgcaccccgctgcctcctccacatctttactttgttgttgttttctg<br>agacagggcttcattctgtctcccgggctgcagtgcagtggtacaatca<br>cagctcactgcagctctgcactcctgggctcaggcgatcctcctgcctc<br>agccacttgagtagctgggactgcaggcacacgccaccacgtccag<br>ctatttaaaaaattttgtagagatgggttctggctcacaggtctactttgg<br>atctctacttggcatctccaggacagtgcgatccaatccacactcaaa<br>atcttctcccctagcctgtcccaccccatccctgctgccatccctggtatc<br>ttcatccttgcagtgggcaggtcgaggacaccactgccctccttgaccc<br>ctctcttcctctcttaccccacattgtatttgttagcagatgcttctgaaatat<br>gccctgcacccgcccacctctcagcacggccaccgctgccgtccca<br>gcccacaagcccacggccctttcctcccgacacacctcctgcttctctg<br>ccaggttgcgccagcctcctctcaaggcaggtgcagcttaccctgctc<br>accattccctccctagcttcatgggcttggttact | Non-Risk/NKX6-3-Risk Nonrisk for rs750625 |

| | | |
|---|---|---|
| gNCH012 | gcttcatgggcttggttactctgttagggcacgtgccaggcacggtctcc<br>ttcaaagggctaggccttctgtggggcatcctccctgagatgtcctcac<br>cctcaccctcaattccttgaagtctctcagctggcactcatgcaggcctc<br>ccctggccacccatctgtaattccagtcagccttctttccacacacacg<br>cctacctcctgtcctccttctctgcttgatttttctttttagcacgtattatctag<br>aatttctgttttaatgacttctaattttatttactttatcttgtttctagttagtttcc<br>ctgctagaaccacgcttcttgcaggtagagactttttctgtttcattcaat<br>gctggcacccaggcactctttattggtgcctggcacacagtaggcact<br>caataaatattgttgagtaagtggatgaaattcatctgtccccatgaga<br>aggctgaccaaggtcacagcaccaggaagtgacaaaataggccta<br>gagcccaggcgttcatgaacccagttagcaacactgctgcggtaatg<br>caggggccagcatgcagcccctgcaggtgctcaataaatgcttgttga<br>aggaatggccgtcccaggaaggtgtctctctcccgggaaggccacta<br>tttcactttaaaccagcctaagctccatctgttcaatcctccccatgaatc<br>caaagatttgtgtaggaaaaaaaattaaatctttgctgtctcaacttcca<br>aacgacttcctttcccttcctgttagggtattggggccatgctgcatggat<br>cctgtggaatggccccactagcctgggcctcagggtggcactgaggg<br>gtcacacaggtgggactgaggggtcacacaggtggggaaaggaca<br>tgagaacagctttcctttacctgaccagtcctgagctccttggggtgggg<br>gcctggaacagctgcctgggggcacaagaaccatgtccttactttagg<br>gaaaactcagcaagccaaggtgacctggaggaaaggggattcctct<br>gtcatccgcttcctgtgctagcccaggaggacaacgtgagccccatct<br>catgtctgtctcatctctttctacaatggcaagggcctggcttgtcttggca<br>tccttccctgcactcagcggaggggcatggtgtaggccctggcccag<br>atgggacgatgagagctccaaggtgagccggtcctcggccgcagac<br>ttgcagtgtctgggaaatgcatcaccttctcctgacctgccgacgctgct<br>tttctgctgcggcacagggcgggggcttctcagtgcgggaagcgtgg<br>ggaagagagcacccactgctctacagctgaaagctccgtcctgcccc<br>agttaagcccaccagcccctccttcaaagggggtaagtggggaggg<br>cgccattatcagctccgtcacatgcaggctgtgtgactgcacacaaat<br>aacttccctgggtactggtgcttctaaatatggtcttcaactaaatatggc<br>cttcaaccggaagcacgcaaggggagaggagctccgaacgccag<br>ctccagcagcagctgacccacggtcttccccagcctcatgattgctcttt<br>taagtgagccatcctggtgctctaatgtccctaagtgagcccttcccgtg<br>cactaatggctctaagtgagcactccaggtaccctaatgtccctaagtg<br>agctctcccggcaccccgatgtccctaagtgagacctcccggcaccc<br>cgatgtccctaagtgagccctcccggcaccccgatgtccctaagtgag<br>ccctcccggcgccccgatgtccctaagtgagtcttcccggcgc | Non-Risk/NKX6-3-Risk Nonrisk for rs3802315<br>rs3802316<br>rs59864562 |
| gNCH013 | agtggatattattaactctgtttccttgatgaggaagctggggttcagag<br>ccaactgttagcagggttgggaagtgagaatctctgcctcgagacac<br>aacaacaagttgtttgtcccacatcatgctggggtggctgggaggga<br>tagggtgagacagggagcagccactccactgaagaccaggccatg<br>cagaggggatgagaagggcagcgttacctcccgagaggctactcc<br>aaggagagcggctcggggtcactgtttccccctttttcaggctggcccg<br>cttcactatctgcgccccccttcctgccctctatcaggtccggctgtaggc<br>cactccctctcagctccacctgcagacagcagcagagacagaaagc<br>aggacagatcgcaaagacgggcagaacacaggcagaagatcga<br>aaggaggccacacggaggtggcagacacactgcccaccagggag<br>aagagagactggagagagagctcataaagaggtgagtggaggga<br>ggtgtcacgcagacggcccggcagagcaaggcagcagtgatggc<br>gctgggtcccccagggccgcgggcgcctcagtaccttggagtgtttca<br>taaagtaatcaatatcgacctccagctcactgggatcctccaggggcc<br>cctctacagtcacctcctcgtgctcctgggcggcatcggcgctggaca<br>agtctatctgtcgaaccaccttgcgaatgatctgggaaaggaaggga<br>aggaggaaagggctggtcaggccgggctcgggggctcatgtctgta<br>atctcagcactttgggaggctgacgtgcagatcatctgaggtcaggag | rs516946_rs515071_riskSNP |

| | | |
|---|---|---|
| | tttgagaccagcctggccaacatggtgaaaccctgtctctactaaaat<br>acaaaaattagcccggtgtggtggcaggtgcctgtaatcccagctact<br>caggaggctgtggcaggagaatcacttgaacctggtggatggaggtt<br>gcagtgagctgagattgcatcactgcactccagcctggttgacagagc<br>aacactctg | |
| gNCH014 | ttctgacagctgaccaggaagaagctcagcagcaccagcgtgacca<br>acagctgggtgacgaaagtccacatcctcgcctcctcaccagcccgg<br>tcctctggcagccagggccccggccaccacgggggggcctgcctctc<br>attctccactgggactcctgagtcccccagggaccccttgcattccccc<br>tccctatctctctggtttgctctcttggtctctcactgtttctctctctgtgggca<br>ggacaccgaatggccctctcggctgaagcagcccggcccaagtgcc<br>cttctgcccagcagccaggctctgccctgagtggcccgggtgacatca<br>cgctcctataattttacctccccaaaccagctgccggagctgtccaacc<br>gggctagaaggaagccggcaggtgcccatgtcctgagcactgaaa<br>gcacaagtgaaagtcactgtgtgagtgacacccgctgctgcctaacc<br>ccgccctgcccagagacctggaagtgggggctcggcctctcgcctat<br>cctgcccatccccatgggaactagaaagagttctgcccagggaaagt<br>ttcttactttcagaaaataatacagtccgagaaagcagctacttgtgtgt<br>ctccttgggctgctactgcggtgatccaaatgctgttccccggcatggta<br>gggatgagaccagctaataccctgggtacttagcggtctgtttcaaaa<br>attgctccaaaagggtgagtgatgggctcagggacagtcaagtgagc<br>ccagcaggggcccagcacaggtggtggggccacctggaagccggt<br>ctgtaattcctgtcagcccacaggcccttgcacccttcctttatggggtac<br>agcctcccaaaactgaagcatggagctggtgtgtctatttaaggcctgt<br>tcttaaaggctcttccagggctgggctatcaacgcccactgagcagga<br>acaacgcgtattaaacttgtatttacagagctgattcacaaatgtagcc<br>gcagcagctgcggctggaaaaaatttcactttctagatcaggttgttcct<br>ctctgagtggcaatgcagcatggcagaaagaacacagatc | 508419_riskSNP |
| gNCH018 | ggccaatacggtgaaacccatctctactaaaaatacaaaaattagac<br>gggcatagtggcgggtgcctgtagtcccagctactcgggaggctgag<br>gcaggagaattgcttgagcctgggaggcagaggttgcagtgagctg<br>agatcgcaccactgcactctggcctgggtgacagaacgagactctgt<br>ctcaggaaacaaaacaaaacaaaaaccccaaaacaaatcaaac<br>acttcgatagcaagaaaacaaataacccaattaaaaaatgggcctg<br>ggacgtgaatagacatttctgaaaataagacatacaaatgaccaaca<br>gacatgaaaaaacaacgctccacctcactaatcattagggaatgca<br>aattaaaaccacaatgagctatcgcctcacacctgtcagaatgactgtt<br>atcaaaaatgcaagtgttgttgaggatgtgaagtaaagaaaacccttg<br>cctgctcttggtaggaatgtaaattagattagc | Homology arm SNPs for<br>payloads |
| gNCH019 | gcccaagtgcccttctgcccagcagccaggctctgccctgagtggcc<br>cgggtgacatcacgctcctataattttacctccccaaaccagctgccgg<br>agctgtccaaccgggctagaaggaagccggcaggtgcccatgtcct<br>gagcactgaaagcacaagtgaaagtcactgtgtgagtgacacccgc<br>tgctgcctaaccccgccctgcccagagacctggaagtgggggctcg<br>gcctctcgcctatcctgcccatccccatgggaactagaaagagttctgc<br>ccagggaaagtttcttactttcagaaaataatacagtccgagaaagca<br>gctacttgtgtgtctccttgggctgctactgcggtgatccaaatgctgttcc<br>ccggcatggtagggatgagaccagctaataccctgggtacttagcgg<br>tctgtttcaaaaattgctccaaaagggtgagtgatgggctcagggaca | risk for rs508419,<br>rs9694034,rs6989203,rs113543<br>09 |

| | | |
|---|---|---|
| | gtcaagtgagcccagcaggggcccagcacaggtggtggggccacc tggaagccggtctgtaattcctgtcagcccacaggcccttgcacccttc ctttatggggtacagcctcccaaaactgaagcatggagctggtgtgtct atttaaggcctgttcttaaaggctcttccagggctgggctatcaacgccc actgagcaggaacaacgcgtattaaacttgtatttacagagctgattca caaatgtagccgcagcagctgcggctggaaaaaatttcactttctaga tcaggttgttcctctctgagtggcaatgcagcatggcagaaagaacac agatctgggacacagcagactaaccctgcaactcacctactgtgggg tcttgggaaggctggggaccttctctgtgcctcagtttcctcatctgtgaa gcacggattaaagtgctctccccaccctttctcaaggctgctgggaca atcatgccgcttatgaatgctaacactatgaatgataatggcaaacgct tgtgatagtgcatggttctatgtggatttcgcattaattcactgacacctca ccacagtcctaaccaaggaggttctatcgtcttcccattttacacaaag gaaacagaagcaaagtattctgcctaggtggaggagtaagtaaacg gaggagtcacattccaaatgcctctcactccacaatgctgcttatttcaa caagaaagcacggggaaaagtatgtgcaaagaagccgggg | |
| gNCH021 | Gcccaagtgcccttctgcccagcagccaggctctgccctgagtggcc cgggtgacatcacgctcctataattttacctccccaaaccagctgccgg agctgtccaaccgggctagaaggaagccggcaggtgcccatgtcct gagcactgaaagcacaagtgaaagtcactgtgtgagtgacacccgc tgctgcctaaccccgccctgcccagagacctggaagtgggggctcg gcctctcgcctatcctgcccatccccatgggaactagaaagagttctgc ccagggaaagtttcttactttcagaaaataatacagtccgagaaagca gctacttgtgtgtctccttgggctgctactgcggtgatccaaatgctgttcc ccggcatggtagggatgagaccagctaataccctgggtacttagcgg tctgtttcaaaaattgctccaaaagggtgagtgatgggctcagggaca gtcaagtgagcccagcaggggcccagcacaggtggtggggccacc tggaagccggtctgtaattcctgtcagcccacaggcccttgcacccttc ctttatggggtacagcctcccaaaactgaagcatggggctggtgtgtct atttaaggcctgttcttaaaggctcttccagggctgggctatcaacgccc actgagcaggaacaacgcgtattaaacttgtatttacagagctgattca caaatgtagccgcagcagctgcggctggaaaaaatttcactttctaga tcaggttgttcctctctgagtggcaatgca gcatggcagaaagaacacagatctgggacacagcagactaaccct gcaactcacctactgtggggtcttgggaaggctggggaccttctctgtg cctcagtttcctcatctgtgaagcacggattaaagtgctctccccaccctt tctcaaggctgctgggacaatcatgccgcttatgaatgctaacactatg aataataatggcaaacgctgtgatagtgcatggttctatgtggatttcgc attaattcactgacacctcaccacagtcctaaccaaggaggttctatcg tcttcccattttacacaaaggaaacagaagcaaagtattctgcctaggt ggaggagtaagtaaacggaggagtcacattccaaatgcctctcactc cacaatgctgcttatttcaacaagaaagcacggggaaaagtatgtgc aaagaagccgggg | risk for rs508419 |

**Table S4.** Primers for PCR-amplified gene fragments and junction verifications

| Primer Name | Sequence 5' -> 3' | Description |
|---|---|---|
| oNCH016 | gcctggtgacaaaagtgaga | Base Construct Assembly Left Junction Rev |
| oNCH043 | caacattttgcgcacggttat | Base Construct Assembly Left Junction Fwd |
| oNCH41 | aggtgagatggacaaagtgc | Base Construct Assembly Right Junction Fwd |
| oNCH42 | ctatatgaaaagccggttccgg | Base Construct Assembly Right Junction Rev |
| oNCH102 | tagggcatggcgtggcccaggcagtgctggggtgagaatcttaaccctagaaagataatcatattgtgacg | Fwd Amplify MC2 With Homology to Base Construct |
| oNCH103 | agccagcaagtccctgagccaggaggcagccgtacctgccttaaccctagaaagatagtctgcgt | Rev Amplify MC2 With Homology to Base Construct |
| oNCH104 | gacatgaactctcagcaagg | MC2 Junction Primer Left Fwd |
| oNCH105 | tttcaagaatgcatgcgtca | MC2 Junction Primer Left Rev |
| oNCH106 | cataagctcaggagtccatttct | MC2 Junction Primer Right Fwd |
| oNCH017 | gcattctagttgtggtttgtcca | MC2 Junction Primer Right Rev |
| oNCH049 | atggctgattatgatccggctg | MC1 Left Homology arm Junction Fwd |
| oNCH050 | gtAGCGGCTGAAGCACTGCA | MC1 Left Homology arm Junction Rev |
| oNCH051 | gtgtctctcactcggtgcgt | MC1 Right Homology arm Junction Fwd |
| oNCH052 | ctctgacttgagcgtcgatt | MC1 Right Homology arm Junction Rev |
| oNCH078 | actggagagagagctcataaagaggtgagtggagggaggtgtcacgcaga | ANK1-Risk Segment 2 FWDFwd |
| oNCH079 | ctgaccagccctttcctccttcccttcctttcccagatcattcgcaaggt | ANK1-Risk Segment 2 REVRev |
| oNCH080 | accttgcgaatgatctgggaaa | ANK1-Risk Segment 3 FWDFwd |
| oNCH081 | ctgggcagaactctttctagttcccatggggatgggcaggataggcgaga | ANK1-Risk Segment 3 REVRev |
| oNCH082 | tctgcgtgacacctccctccact | ANK1-Risk Segment 1 REVRev |
| oNCH083 | tgagaagggcagcgttacct | ANK1-Risk Segment 1 FWDFwd |
| oNCH084 | tcggcctctcgcctatcctg | ANK1-Risk Segment 4 FWDFwd |
| oNCH085 | tgcttcagttttgggaggctgt | ANK1-Risk Segment 4 REVRev |
| oNCH086 | gctggcccgcttcactatctgcgccccccttcctgccctctgtcggggctggcttaactatg | URA3 + Homology to NKX6-3/Ank1 for ANK1-Risk Construction Fwd |
| oNCH087 | accccataaaggaagggtgcaagggcctgtgggctgacagctccttacgcatctgtgcgg | URA3 + Homology to NKX6-3/Ank1 for ANK1-Risk Construction Rev |
| oNCH089 | cctaaagcaatccttttgcctcga | ANK1-Risk Seg 1 jxnJunction Fwd |
| oNCH090 | aaacactccaaggtactgaggc | ANK1-Risk Seg 1 jxnJunction Rev |
| oNCH091 | atcaggtccggctgtaggcca | ANK1-Risk Seg 2 Junction Fwdjxn |
| oNCH092 | agatgatctgcacgtcagcctc | ANK1-Risk Seg 2 Junction Revjxn |

| | | |
|---|---|---|
| oNCH093 | aatatcgacctccagctcactg | ANK1-Risk Seg 3 Junction Fwdjxn |
| oNCH094 | caagtagctgctttctcggactgt | ANK1-Risk Seg 3 Junction Revjxn |
| oNCH095 | agtcactgtgtgagtgacaccc | ANK1-Risk Seg 4 Junction Fwdjxn |
| oNCH096 | tgatagcccagccctggaaga | ANK1-Risk Seg 4 Junction Revjxn |
| oNCH097 | agccagcaagtccctgagccaggaggcagccgtacctgccttaaccct agaaagatagtc | MC2+40bp Homology Fwd |
| oNCH098 | gattatctttctagggttaagattctcaccccagcactgcctgggccacgcc atgcccta | MC2 +40bp Homology Rev |
| oNCH099 | tacagccggacctgatagag | ANK1-Risk Seg Jxn |
| oNCH100 | aagatcgaaaggaggccaca | ANK1-Risk Seg Jxn |
| oNCH101 | acagtccgagaaagcagctact | ANK1-Risk Seg Jxn |
| oNCH102 | tagggcatggcgtggcccaggcagtgctggggtgagaatcttaaccta gaaagataatcatattgtgacg | MC2 Primer With Homology To Ank1(Longer) |
| oNCH103 | agccagcaagtccctgagccaggaggcagccgtacctgccttaaccct agaaagatagtctgcgt | MC2 Primer With Homology To Ank1 (Longer) |
| oNCH104 | gacatgaactctcagcaagg | MC2 Junction Primer |
| oNCH105 | tttcaagaatgcatgcgtca | MC2 Junction Primer |
| oNCH106 | cataagctcaggagtccatttct | MC2 Junction Primer |
| oNCH107 | aactctttctagttcccatggggatgggcaggataggcgagaggccgag c | New ANK1-Risk Rs508419 Primer |
| oNCH108 | ccctcctcctcccctgcacctgctgctcctcctccacgtgtcggggctggctt aactatg | URA3 + Homology To NKX6-3/Ank1 For All-Risk Rev |
| oNCH109 | tttcagagagtgaaagctggagcttgtgagtctagaatcactccttacgca tctgtgcgg | URA3 + Homology To NKX6-3/Ank1 For All-Risk Fwd |
| oNCH110 | ctccacacgatgcctagaga | Segment 1 For All-Risk Rev |
| oNCH111 | gtctatacatcagagtagggtaggggattc | Segment 2 For All-Risk Fwd |
| oNCH112 | tgaagagtgcattcggggtt | Segment 2 For All-Risk Rev |
| oNCH113 | ttggtcaggctcgtctcgag | Segment 3 For All-Risk Rev |
| oNCH114 | cttggtggtacgaacatcc | URA3 Rev Junction Primer |
| oNCH115 | agcagaattgtcatgcaagg | URA3 Fwd Junction Primer |
| oNCH116 | gctcagacacctgtgtccaa | All-Risk Seg 1.2 Fwd |
| oNCH117 | gcaacagtgagtacgggtcc | All-Risk Seg 1.1 Rev |
| oNCH118 | tgatggggtcttgctctgttg | ANK1-Risk Segment 1 Jxn Fwd Updated |
| oNCH119 | ttaatacgcgttgttcctgctc | ANK1-Risk Segment 4 Jxn Rev Updated |
| oNCH120 | ttcttctgcagccagcagct | All-Risk Vector/Seg1.1 Jxn Fwd |
| oNCH121 | actgagtctttggatgaggg | All-Risk Vector/Seg1.1 Jxn Rev |
| oNCH122 | aaagacatgcaggagtgtctg | All-Risk Seg 1.1/1.2 Jxn Fwd |
| oNCH123 | gcccagattcatgcttgatgct | All-Risk Seg 1.1/1.2 Jxn Rev |
| oNCH124 | ttctctacatctggaactgccgg | All-Risk Seg 1.2/1.3 Jxn Fwd |
| oNCH125 | cagcccccaggtagggaatt | All-Risk Seg 1.2/1.3 Jxn Rev |
| oNCH126 | tctgctggagcaatggccat | All-Risk Seg 1.3/1.4 Jxn Fwd |
| oNCH127 | tgagagagcaagccctgcaga | All-Risk Seg 1.3/1.4 Jxn Rev |
| oNCH128 | cattggttcggctggtgaag | All-Risk Seg 1.4/Seg2 Jxn Fwd |
| oNCH129 | ctccagactccgctttggca | All-Risk Seg 1.4/Seg2 Jxn Rev |
| oNCH130 | actgaaagacaggccacagc | All-Risk Seg2/Seg3.1 Jxn Fwd |
| oNCH131 | tgttgtctctctctctgccaatttc | All-Risk Seg2/Seg3.1 Jxn Rev |
| oNCH132 | cagcagatccttgtacttaggaagag | All-Risk Seg 3.1/3.2 Jxn Fwd |

| | | |
|---|---|---|
| oNCH133 | ttcaccccaggttggaggct | All-Risk Seg 3.1/3.2 Jxn Rev |
| oNCH134 | attagtcgctgtttgaggctgtg | All-Risk Seg 3.2/Vector Jxn Fwd |
| oNCH135 | ccttaggagcagtggttcagt | All-Risk Seg 3.2/Vector Jxn Rev |
| oNCH136 | atcgaattcctgcagcccggggttaattaaaagatcgaaaggaggcca ca | Non-Risk/NKX6-3-Risk Chunk 1 Left Fwd Homology Arm |
| oNCH137 | gaggctgacgtgcagatcatctggcgcgcctccagttccaggctgggca t | Non-Risk/NKX6-3-Risk Chunk 1 Right Fwd Homology Arm |
| oNCH138 | ccaccgcggtggcggccgctctagaactagtgttaattaaaagtatgtgc cccaggcagg | Non-Risk/NKX6-3-Risk Chunk 1 Right Rev Homology Arm |
| oNCH139 | tcgaattcctgcagcccggggttaattaaggaacatgtcgttcatgtacat actaaacac | Non-Risk/NKX6-3-Risk Chunk 2 Left Fwd Homology Arm |
| oNCH140 | gactcacttaggcgcgcccctcactggtaaaggagggagtgattggaa | Non-Risk/NKX6-3-Risk Chunk 2 Left Rev Homology Arm |
| oNCH141 | accagtgaggggcgcgcctaagtgagtcttcccggcgc | Non-Risk/NKX6-3-Risk Chunk 2 Right Fwd Homology Arm |
| oNCH142 | cggccgctctagaactagtgttaattaagcacatacattcagtctggcaga tg | Non-Risk/NKX6-3-Risk Chunk 2 Right Rev Homology Arm |
| oNCH143 | Tgcacgtgcaaggccagcatgcccagcctggaactggagctccttacg catctgtgcgg | URA3 Chunk 1 Fwd |
| oNCH144 | aatctcagcactttgggaggctgacgtgcagatcatcttgtcggggctggc ttaactatg | URA3 Chunk 1 Rev |
| oNCH145 | tggcgacattccaatcactccctcctttaccagtgaggtgtcggggctggc ttaactatg | URA3 Chunk2 Fwd |
| oNCH146 | cttacttaggacatcggggcgccgggaagactcacttaggctccttacgc atctgtgcgg | URA3 Chunk2 Rev |
| oNCH148 | accgggaggactcagggacatc | Chunk2 URA3 Jxn Rev |
| oNCH149 | gggtcacacagtggaccaat | Chunk 1 URA3 Jxn Rev |
| oNCH150 | tggaagggccttctagatcacaca | Chunk 1 - Rev Primer To Amplify gDNA |
| oNCH151 | atcatcatggggcctgggaa | Chunk 2 - Fwd Primer To Amplify gDNA |
| oNCH152 | gtgctcacttagggacatga | Chunk 2 - Rev Primer To Amplify gDNA |
| oNCH157 | aatgtcactgtcacaatgcactgg | Chunk 1.2 Fwd From gNCH010 |
| oNCH158 | atgcccagcctggaactgga | Chunk 1.2 Rev From gNCH010 |
| oNCH159 | ctagcctggcgtctccccct | Part Of Chunk 2.1fwd gNCH011 |
| oNCH160 | agtaaccaagcccatgaagc | Part Of Chunk 2.1rev gNCH011 |
| oNCH161 | ggctccggtgcccgtcagtg | Ef1alpha Fwd |
| oNCH163 | aggagaactaggcgcccagg | gNCH007 Fwd |
| oNCH164 | ttccccattcccaggcccca | gNCH007 Rev |
| oNCH165 | ctggcagtgaggacagaggt | Non-Risk/NKX6-3-Risk Chunk2 Rev Junction Primer |
| oNCH169 | acgtctcagactttggatgggcaggataggcgaggttttagagctagaaa tagcaagtta | Primers For Creepy Assembly For All-Risk Rs516946, Rs515071, Rs508419 |
| oNCH170 | acgtctcccaaggtggttctgcgcaagcccggaatcgaaccggg | Primers For Creepy Assembly For All-Risk Rs516946, Rs515071, Rs508419 |

| | | |
|---|---|---|
| oNCH171 | acgtctccgaaccaccttggttttagagctagaaatagcaagtta | Primers For Creepy Assembly For All-Risk Rs516946, Rs515071, Rs508419 |
| oNCH172 | acgtctccaaactctgcgtgacacctccctcctgcgcaagcccggaatc gaaccggg | Primers For Creepy Assembly For All-Risk Rs516946, Rs515071, Rs508419 |
| oNCH173 | acgtctcacttgcgaatgatcgttttagagctagaaatagcaagtt | Primers For Creepy Assembly For All-Risk Rs516946, Rs515071, Rs508419 |
| oNCH305 | GactttGGATGGGCAGGATAGGAGAG | All-Risk Rs508419 gRNA Primer 1 |
| oNCH306 | AaacCTCTCCTATCCTGCCCATCCaa | All-Risk Rs508419 gRNA Primer 2 |
| oNCH181 | tacaaagaagcttgaggagacctttctggggtgacaaatgtcctttttcttct ttgaatt | Non-Risk/NKX6-3-Risk Last Snps Seg1 Fwd |
| oNCH182 | tgcagtaatgcaatctcagttcact | Non-Risk/NKX6-3-Risk Last Snps Seg1 Rev |
| oNCH183 | gctcacgcttgtaatcccagcactttgggaggccgaggtgggcggatcat gaggtcagga | Non-Risk/NKX6-3-Risk Last Snps Seg4 Rev |
| oNCH184 | tagtagagacggggtttcaccgtgttagccaggatggtctcgatctcctga cctcatgat | Non-Risk/NKX6-3-Risk Last Snps Seg5 Fwd |
| oNCH185 | gtagagacggggatctcaccatgt | Non-Risk/NKX6-3-Risk Last Snps Seg5 Rev |
| oNCH186 | tgcctcagcctcccaaagtgctgggattacagggttgagtgtgtcggggct ggcttaact | Non-Risk/NKX6-3-Risk URA3 + Homology Arm For Last Snps |
| oNCH187 | catgaatgagcctcaagtgttctgtggcgcaaagctgcccctccttacgc atctgtgcgg | Non-Risk/NKX6-3-Risk URA3 + Homology Arm For Last Snps |
| oNCH192 | agtcttgctctgttgcccag | Jxn Primer For Last Snps Non-Risk/NKX6-3-Risk |
| oNCH193 | tggccgggctcagtgactcaa | Jxn Primer For Last Snps Non-Risk/NKX6-3-Risk |
| oNCH194 | acttagggaggctgaggcag | Jxn Primer For Last Snps Non-Risk/NKX6-3-Risk |
| oNCH195 | agagtttcacccttgttgccc | Jxn Primer For Last Snps Non-Risk/NKX6-3-Risk |
| oNCH200 | cagctgtagggctcacttca | Right Jxn Primer For Last Snps Non-Risk/NKX6-3-Risk Fwd Updated |
| oNCH201 | gtggttgttgtggcaattaa | Left Jxn Primer For Last Snps Non-Risk/NKX6-3-Risk Rev |
| oNCH215 | ttctttgaattgatggcttc | Vswap-In Non-Risk/NKX6-3-Risk Seg 1 Rev |
| oNCH216 | acagtgcattttgaagagcagaagtt | Vswap-In Non-Risk/NKX6-3-Risk Seg 3 Fwd |
| oNCH217 | tagcagacaaaggactgggcatg | Vswap-In Non-Risk/NKX6-3-Risk Seg 3 Rev |
| oNCH218 | aggccgaggtgggcggatcatgaggtcaggagatcgagacca | Vswap-In Non-Risk/NKX6-3-Risk Seg 4 Rev Non Specific |
| oNCH219 | tggtctcgatctcctgacctcatgatccgcccacctcggcc | Vswap-In Non-Risk/NKX6-3-Risk Seg 5 Fwd Nonspecific |
| oNCH295 | ggtgcctgtagtcccagctactcgggaggctgaggcaggagtgtcggg gctggcttaact | Left Homology arm Payload Snp_URA3_Fwd |
| oNCH296 | aacaacacttgcattttgataacagtcattctgacaggtctccttacgcatc tgtgcgg | Left Homology arm Payload Snp_URA3_Rev |

| | | |
|---|---|---|
| oNCH297 | attagacgggcatagtggcg | Left Homology arm Payload Snp_URA3_Jxn_Fwd |
| oNCH298 | tttacattcctaccaagagc | Left Homology arm Payload Snp_URA3_Jxn_Rev |
| oNCH313 | gctgcctaaccccgccctgcccagagacctggaagtgggggtgtcggg gctggcttaact | URA3 Fwd For Rs508419, Rs9694034,Rs6989203,Rs113 54309 |
| oNCH314 | gtgtcagtgaattaatgcgaaatccacatagaaccatgcactccttacgc atctgtgcgg | URA3 Rev For Rs508419, Rs9694034,Rs6989203,Rs113 54309 |
| | **Genotyping Primers for hiPSC payload deliveries** | |
| oNCH267 | gagtggtaagtacagagacc | Lp400 Deletion Junction Fwd Primer |
| oNCH268 | tgaggcatttctaaccctcctt | Lp400 Deletion Junction Rev Primer |
| oNCH0053 | aagaactcaaacagaaggccgg | MC1 insertion Left junction FWD |
| oNCH0059 | cccaacttctcggggactgtgg | MC1 insertion Left junction REV |
| oNCH0051 | gtgtctctcactcggtgcgt | MC1 insertion Right junction FWD |
| oNCH0154 | gctgatccctgccttcaaag | MC1 insertion Right junction REV |
| oNCH0057 | tgtgcgtgattccatttgaatcttc | Inner NKX6-3/ANK1 region FWD |
| oNCH0058 | gtcattcaagataagtctggccctc | Inner NKX6-3/ANK1 region REV |
| oNCH0235 | aagcttctgcacagcaaagg | Left payload insertion junction FWD |
| oNCH0232 | aattgtgtgccctctgacca | Left payload insertion junction REV |
| oNCH0017 | gcattctagttgtggtttgtcca | Right payload insertion junction FWD |
| oNCH0307 | agcagctgcagagctgactc | Right payload insertion junction REV |
| oNCH134 | attagtcgctgtttgaggctgtg | ANK1 inner genotyping primer FWD |
| oNCH135 | ccttaggagcagtggttcagt | ANK1 inner genotyping primer REV |
| oNCH0054 | cgacgatatcgtctacgtacccg | MC1 inner region genotyping primer FWD |
| oWZ883 | gttagcctcccccatctcccgg | MC1 inner region genotyping primer REV |
| oWZ0111 | GGTGCCCTTAAACGCCTGGTTG | Payload backbone FWD |
| oWZ2065 | GATGCTGTCGCCGAAGAAGTTAAG | Payload backbone REV |

**Table S5.** All-Risk Haplotype SNPs relative to Reference Genome

| Chromosome | Position | rsID | Reference Allele | All-Risk Allele | Description |
|---|---|---|---|---|---|
| **chr8** | **41635620** | **.** | **C** | **A** | Additional SNP |
| *chr8* | *41636116* | *rs4736996* | *T* | *C* | Homology arm SNPs |
| *chr8* | *41636311* | *rs4736997* | *T* | *C* | Homology arm SNPs |
| *chr8* | *41636370* | *rs2354577* | *G* | *A* | Homology arm SNPs |
| chr8 | 41646334 | rs5891165 | CAAAG | C | |
| **chr8** | **41646919** | **.** | **C** | **A** | Additional SNP |
| **chr8** | **41646987** | **.** | **T** | **G** | Additional SNP |
| chr8 | 41647443-41647445 | rs35659384 | GGG | GGGG | |
| chr8 | 41648633 | rs4736999 | C | G | |
| chr8 | 41648861 | rs12549294 | A | G | |
| chr8 | 41649718 | rs7825337 | C | T | |
| chr8 | 41649831 | rs7825494 | C | T | |
| chr8 | 41650649 | rs6981607 | A | C | |
| **chr8** | **41651008** | **rs56300007** | **C** | **T** | Additional SNP |
| **chr8** | **41651010** | **rs554421053** | **C** | **T** | Additional SNP |
| **chr8** | **41651031** | **rs72168676** | **G** | **GTT** | Additional SNP |
| chr8 | 41651740 | rs12549902 | G | A | |
| **chr8** | **41652038** | **.** | **C** | **A** | Additional SNP |
| chr8 | 41652208 | rs4737000 | A | C | |
| chr8 | 41652396 | rs4736819 | C | T | |
| chr8 | 41652741 | rs12550613 | G | C | |
| chr8 | 41655068-41655081 | rs57963234 | ACAGCAGAGTCTAT | - | |
| **chr8** | **41655305** | **rs5891167** | **C** | **CT** | Additional SNP |
| chr8 | 41656012 | rs17659386 | T | C | |
| chr8 | 41656290 | rs17659428 | T | C | |
| chr8 | 41657369 | rs12544241 | G | A | |
| chr8 | 41657611 | rs6990613 | A | G | |
| chr8 | 41658012 | rs716199 | A | G | |
| chr8 | 41658474 | rs10504042 | G | A | |
| chr8 | 41661730 | rs516946 | T | C | |
| chr8 | 41661944 | rs515071 | A | G | |
| chr8 | 41665473 | rs508419 | A | G | |

**Table S6.** ANK1-Risk Haplotype SNPs relative to Reference Genome

| **Chromosome** | **Position** | **rsID** | **Reference Allele** | **ANK1-Risk Allele** | **Description** |
|---|---|---|---|---|---|
| *chr8* | *41636116* | *rs4736996* | *T* | *C* | Homology arm SNPs |
| *chr8* | *41636311* | *rs4736997* | *T* | *C* | Homology arm SNPs |
| *chr8* | *41636370* | *rs2354577* | *G* | *A* | Homology arm SNPs |
| chr8 | 41661730 | rs516946 | T | C | |
| chr8 | 41661944 | rs515071 | A | G | |
| chr8 | 41665473 | rs508419 | A | G | |

**Table S7.** Non-Risk Haplotype SNPs relative to Reference Genome

| Chromosome | Position | rsID | Reference Allele | ANK1-Risk Allele | Description |
|---|---|---|---|---|---|
| *chr8* | *41636116* | *rs4736996* | *T* | *C* | Homology Arm SNPs |
| *chr8* | *41636311* | *rs4736997* | *T* | *C* | Homology Arm SNPs |
| *chr8* | *41636370* | *rs2354577* | *G* | *A* | Homology Arm SNPs |
| chr8 | 41664498 | rs146025758 | AAAGAG | A | |
| chr8 | 41665840 | rs9694034 | A | G | |
| chr8 | 41666227 | rs6989203 | G | A | |
| chr8 | 41666241 | rs11354309 | CT | C | |
| chr8 | 41666907 | rs28591316 | A | G | |
| chr8 | 41666987 | rs28602970 | A | G | |
| chr8 | 41667170 | rs35273426 | C | T | |
| chr8 | 41667381 | rs10109812 | A | G | |
| chr8 | 41668396 | rs750625 | C | T | |
| chr8 | 41670660 | rs3802315 | G | T | |
| chr8 | 41670710 | rs3802316 | G | C | |
| chr8 | 41671654 | rs59864562 | A | G | |
| chr8 | 41675682 | rs10106563 | C | A | |
| chr8 | 41675996 | rs13266210 | A | G | |
| chr8 | 41676550 | rs11995075 | T | C | |
| chr8 | 41679800 | rs62508166 | G | A | |

**Table S8.** Non-Risk Haplotype SNPs relative to Reference Genome

| Chromosome | Position | rsID | Reference Allele | Non-Risk Allele | Description |
|---|---|---|---|---|---|
| *chr8* | *41636116* | *rs4736996* | *T* | *C* | Homology Arm SNP |
| *chr8* | *41636311* | *rs4736997* | *T* | *C* | Homology Arm SNP |
| *chr8* | *41636370* | *rs2354577* | *G* | *A* | Homology Arm SNP |
| chr8 | 41664498 | rs146025758 | AAAGAG | A | |
| chr8 | 41665840 | rs9694034 | A | G | |
| chr8 | 41666227 | rs6989203 | G | A | |
| chr8 | 41666241 | rs11354309 | CT | C | |
| chr8 | 41666907 | rs28591316 | A | G | |
| chr8 | 41666987 | rs28602970 | A | G | |
| chr8 | 41667170 | rs35273426 | C | T | |
| chr8 | 41667381 | rs10109812 | A | G | |
| chr8 | 41668396 | rs750625 | C | T | |
| chr8 | 41670660 | rs3802315 | G | T | |
| chr8 | 41670710 | rs3802316 | G | C | |
| chr8 | 41671654 | rs59864562 | A | G | |
| chr8 | 41675682 | rs10106563 | C | A | |
| chr8 | 41675996 | rs13266210 | A | G | |
| chr8 | 41676550 | rs11995075 | T | C | |
| chr8 | 41679800 | rs62508166 | G | A | |

**Table S9.** NKX6-3-Risk Haplotype SNPs relative to Reference Genome

| Chromosome | Position | rsID | Reference Allele | All-Risk Allele | Description |
|---|---|---|---|---|---|
| **chr8** | **41635620** | **.** | **C** | **A** | Additional SNP |
| *chr8* | *41636116* | *rs4736996* | *T* | *C* | Homology Arm SNPs |
| *chr8* | *41636311* | *rs4736997* | *T* | *C* | Homology Arm SNPs |
| *chr8* | *41636370* | *rs2354577* | *G* | *A* | Homology Arm SNPs |
| chr8 | 41646334 | rs5891165 | CAAAG | C | |
| **chr8** | **41646987** | **.** | **T** | **G** | Additional SNP |
| chr8 | 41647443-41647445 | rs35659384 | GGG | GGGG | |
| chr8 | 41648633 | rs4736999 | C | G | |
| chr8 | 41648861 | rs12549294 | A | G | |
| chr8 | 41649718 | rs7825337 | C | T | |
| chr8 | 41649831 | rs7825494 | C | T | |
| chr8 | 41650649 | rs6981607 | A | C | |
| **chr8** | **41651006** | **rs1356692773** | **TGCGCGCG CGCGC** | **T** | Additional SNP |
| **chr8** | **41651031** | **rs72168676** | **G** | **GTT** | Additional SNP |
| chr8 | 41651740 | rs12549902 | G | A | |
| chr8 | 41652208 | rs4737000 | A | C | |
| chr8 | 41652396 | rs4736819 | C | T | |
| chr8 | 41652741 | rs12550613 | G | C | |
| chr8 | 41655068-41655081 | rs57963234 | ACAGCAGA GTCTAT | - | |
| **chr8** | **41655129** | **rs373938111** | **G** | **GGT** | Additional SNP |
| **chr8** | **41655305** | **rs5891167** | **C** | **CT** | Additional SNP |
| chr8 | 41656012 | rs17659386 | T | C | |
| chr8 | 41656290 | rs17659428 | T | C | |
| chr8 | 41657369 | rs12544241 | G | A | |
| chr8 | 41657611 | rs6990613 | A | G | |
| chr8 | 41658012 | rs716199 | A | G | |
| chr8 | 41658474 | rs10504042 | G | A | |
| chr8 | 41664498 | rs146025758 | AAAGAG | - | |
| **chr8** | **41665538** | **.** | **G** | **A** | Additional SNP |
| chr8 | 41665840 | rs9694034 | A | G | |
| chr8 | 41666227 | rs6989203 | G | A | |
| chr8 | 41666241 | rs11354309 | CT | C | |
| chr8 | 41666907 | rs28591316 | A | G | |
| chr8 | 41666987 | rs28602970 | A | G | |
| chr8 | 41667170 | rs35273426 | C | T | |
| chr8 | 41667381 | rs10109812 | A | G | |
| chr8 | 41668396 | rs750625 | C | T | |
| chr8 | 41670660 | rs3802315 | G | T | |
| chr8 | 41670710 | rs3802316 | G | C | |

| | | | | | |
|---|---|---|---|---|---|
| chr8 | 41671654 | rs59864562 | A | G | |
| **chr8** | **41671710** | **.** | **CT** | **C** | Additional SNP |
| chr8 | 41675682 | rs10106563 | C | A | |
| chr8 | 41675996 | rs13266210 | A | G | |
| chr8 | 41676550 | rs11995075 | T | C | |
| chr8 | 41679800 | rs62508166 | G | A | |

**Table S10.** gRNAs used for Haplotype building, payload integration, and MC1 integration

| Oligo Name | Sequence (5' --> 3') | Description |
|---|---|---|
| orNCH001 | CACCGTAAGTACAGAGACCAAGGGT | Top strand oligo for gRNA-Cas9 cutting upstream of NKX6-3 (left side). Cloned into pX330. |
| orNCH002 | AAACACCCTTGGTCTCTGTACTTAC | Bottom strand oligo for gRNA-Cas9 cutting upstream of NKX6-3 (left side). Cloned into pX330. |
| orNCH003 | CACCGGTGCTGGGGTGAGAATCGGC | Top strand oligo for gRNA-Cas9 cutting downstream of ANK1 (right side). Cloned into pX330. |
| orNCH004 | AAACGCCGATTCTCACCCCAGCACC | Bottom strand oligo for gRNA-Cas9 cutting downstream of ANK1 (right side). Cloned into pX330. |
| orNCH005 | TAGACTCTCTGCAAGATCAG | For RP11-111B9 BAC digestion at chr8: 41,635,845 (crRNA) |
| orNCH006 | CACATTATACATCTAGTGAA | For RP11-111B9 BAC digestion at chr8: 41,688,341 (crRNA) |
| orNCH007 | GACTTTGTGCTGGGGTGAGAATCGGC | For golden gate assembly to insert MC2 |
| orNCH008 | AAACGCCGATTCTCACCCCAGCACAA | For golden gate assembly to insert MC2 |
| orNCH009 | CACCGGCTTCATGTGGTCGGGGTAG | Top strand oligo for dual gRNA-Cas9 cutting left side of MC1 and synthetic payload. Cloned into pX333 |
| orNCH010 | AAACCTACCCCGACCACATGAAGCC | Bottom strand oligo for dual gRNA-Cas9 cutting left side of MC1 and synthetic payload. Cloned into pX333 |
| orNCH011 | CACCGGAAGTTATGAACGCGCCTGC | Top strand oligo for dual gRNA-Cas9 cutting right side of MC1 and synthetic payload. Cloned into pX333 |
| orNCH012 | AAACGCAGGCGCGTTCATAACTTCC | Bottom strand oligo for dual gRNA-Cas9 cutting right side of MC1 and synthetic payload. Cloned into pX333 |

**Table S11.** qPCR Primers

| Name | Sequence (5' -> 3') | Description |
|---|---|---|
| oNCH0317 | GTCTCCTCTGACTTCAACAGCG | GAPDH Human Fwd |
| oNCH0318 | ACCACCCTGTTGCTGTAGCCAA | GAPDH Human Rev |
| oNCH0321 | TAGCCTCTCGGGAGGATCAC | sANK1 Fwd |
| oNCH0322 | CATGCCAAGAGGGGACTAGC | sANK1 Rev |
| oNCH0346 | GACGTGCCTTCTGAGTCG | MYOD1 Fwd |
| oNCH0347 | CTCAGAGCACCTGGTATATCG | MYOD1 Rev |
| oNCH0279 | GGAGGATGAAGCGGACAAGAAG | Pax7 Fwd |
| oNCH0280 | AGGTCAGGTTCCGACTCCACAT | Pax7 Rev |
| oNCH0340 | GGCTTTCAACCATCTCATTCCCG | Pax3 Fwd |
| oNCH0341 | GTTGAGGTCTGTGAACGGTGCT | Pax3 Rev |
| oNCH0342 | CAGTCCTGTCTGGTCCAGAAAG | MYF5 Fwd |
| oNCH0343 | GTCCACTATGTTGGATAAGCAATC | MYF5 Rev |
| oNCH0334 | CCTATTCGTTGGGGATGACAGAG | NKX6.1 Fwd |
| oNCH0335 | TCTGTCTCCGAGTCCTGCTTCT | NKX6.1 Rev |
